# Long-read sequencing and structural variant characterization in 1,019 samples from the 1000 Genomes Project

**DOI:** 10.1101/2024.04.18.590093

**Authors:** Siegfried Schloissnig, Samarendra Pani, Bernardo Rodriguez-Martin, Jana Ebler, Carsten Hain, Vasiliki Tsapalou, Arda Söylev, Patrick Hüther, Hufsah Ashraf, Timofey Prodanov, Mila Asparuhova, Sarah Hunt, Tobias Rausch, Tobias Marschall, Jan O. Korbel

## Abstract

Structural variants (SVs) contribute significantly to human genetic diversity and disease^1–4^. Previously, SVs have remained incompletely resolved by population genomics, with short-read sequencing facing limitations in capturing the whole spectrum of SVs at nucleotide resolution^5–7^. Here we leveraged nanopore sequencing^8^ to construct an intermediate coverage resource of 1,019 long-read genomes sampled within 26 human populations from the 1000 Genomes Project. By integrating linear and graph-based approaches for SV analysis via pangenome graph-augmentation, we uncover 167,291 sequence-resolved SVs in these samples, considerably advancing SV characterization compared to population-wide short-read sequencing studies^3,4^. Our analysis details diverse SV classes—deletions, duplications, insertions, and inversions—at population-scale. LINE-1 and SVA retrotransposition activities frequently mediate transductions^9,10^ of unique sequences, with both mobile element classes transducing sequences at either the 3′- or 5′-end, depending on the source element locus. Furthermore, analyses of SV breakpoint junctions suggest a continuum of homology-mediated rearrangement processes are integral to SV formation, and highlight evidence for SV recurrence involving repeat sequences. Our open-access dataset underscores the transformative impact of long-read sequencing in advancing the characterisation of polymorphic genomic architectures, and provides a resource for guiding variant prioritisation in future long-read sequencing-based disease studies.

---

Structural variants (SVs) make up most polymorphic basepairs (bp) in the genome^4,7^, and have been causally implicated in a wide variety of common and rare diseases, including diseases of the immune system, cancer and cognitive disability^2,11^. SVs show enrichment on haplotypes identified by genome-wide association studies, and are considerably enriched for expression quantitative trait loci when compared to single-nucleotide polymorphisms (SNPs)^4,12^. A subset of SVs, including variants implicated in diseases, exhibit population stratification^13–15^. Uncovering the full spectrum of SVs at a population-scale is therefore crucial for population genetics and genomic medicine.

Current population-scale catalogues of SVs are considered incomplete, owing to the predominant use of short-read sequencing for their generation^3,4,16,17^. Short reads resolve SV allelic sequences incompletely, and show poor sensitivity in SV-rich regions that coincide with repeats^5,6,18^. Furthermore, the reliance of prior population-scale SV studies on linear genomic references, such as GRCh38 developed by the Genome Reference Consortium, has limited SV analyses in areas of structural haplotype diversity^19,20^. These limitations remain important barriers to fully deciphering the biological underpinnings of SVs, including their formation mechanisms and phenotypic implications.

Recently, the Human Pangenome Reference Consortium (HPRC) released a draft pangenome from 44 diploid long-read assemblies, and showed how this graph-based reference enhances SV discovery^20^. This advancement, along with recent progress in genome assembly construction, underscore the transformative potential of long-reads for interpreting human genome variation^5,7,21–23^. Yet, while the integration of long-read sequencing in diagnostics and disease research^24–26^ is anticipated to become more widespread, these efforts are presently hampered by small sample sizes and a lack of long-read sequencing panels of normal individuals from diverse populations. Such data will be instrumental for variant prioritization^27,28^, with their absence currently limiting our understanding of SV diversity around the globe, which affects downstream applications in research and diagnostics.

Here we applied Oxford Nanopore Technologies (ONT) sequencing to analyse SVs from the 1000 Genomes Project (1kGP) sample collection, which allows unrestricted public data access, data sharing and reuse. This makes this cohort an ideal foundational resource for long-read sequence data analysis. Despite thorough characterization by short-read sequencing in prior studies^3,4,29–33^, the 1kGP sample collection has undergone only limited analysis with long-read sequencing so far^5,20^. This has restricted studies of SVs based on long reads mainly to common haplotypes, while variants unique to specific human populations have remained underrepresented. We performed ONT sequencing of 1,019 samples from 26 populations to intermediate coverage, analogous to how the 1kGP, in its phases I–III, conducted short-read sequencing^29,32,33^. This intermediate coverage approach allows for an improved representation of SVs across a diversity of structural haplotypes. We devised methods that combine linear reference and graph-based techniques for long-read based SV discovery, and engineered a computational framework that leverages graph augmentation for SV genotyping, yielding the most complete population-scale SV dataset to date with full sequence resolution and genotype information. We performed a comprehensive analysis of SV allelic sequences, uncovering SVs exhibiting population-specific patterns, and describing complex patterns of inversions. Our analysis into SV breakpoint junctions implies that a diversity of homology-mediated processes contribute to the formation of deletions, insertions and duplications. LINE-1 and SVA (for SINE-R/VNTR/Alu) retrotransposition activities frequently result in the mobilisation of adjacent unique DNA sequences via transduction. Whether transductions affect the 3′ or 5′ end appears to be determined by the mobile element class and the genomic location of the source element involved. The openly accessible data resource generated through this study establishes a pivotal framework for studies of SVs, and provides a reference dataset for future research into their biology and disease relevance at a population-scale.

## Results

### Long-read sequencing and graph-based SV determination in 26 global populations

#### ONT sequencing in 1,019 samples from the 1kGP cohort

We selected samples from the 1kGP collection available at Coriell, sourcing genomic DNA and discarding low-quality DNA (**Table S1**). Post size-selection for ≥25kb fragments (**Fig. S1**), the samples underwent sequencing using a PromethION 48 with R9.4.1 flow cells. Leveraging previously generated high-depth short-read sequencing data from the 1kGP cohort^3^, we implemented rigorous quality control measures to effectively eliminate instances of sample swaps and contaminations (**Methods**). After quality filtering, our assembled study cohort comprised 1,019 high-quality long-read genomes (**Fig. 1a**). These represent donors with self-identified ancestries from 26 geographic locations^34^ (referred to as populations for consistency with prior 1kGP literature^3,29,32,33^), each of which is represented by a minimum of 32 genomes. These populations represent five larger continental areas – comprising 189 genomes from donors with ancestries from Europe (EUR), 192 from East Asia (EAS), 199 from South Asia (SAS), 275 from Africa (AFR), and 164 from the Americas (AMR). The genomes exhibit a median ONT coverage of 16.9× as well as median ONT read length N50 of 20.3 kb, measures largely consistent across continental groups (**Fig. 1b, Fig. S2**).

**Figure 1:**
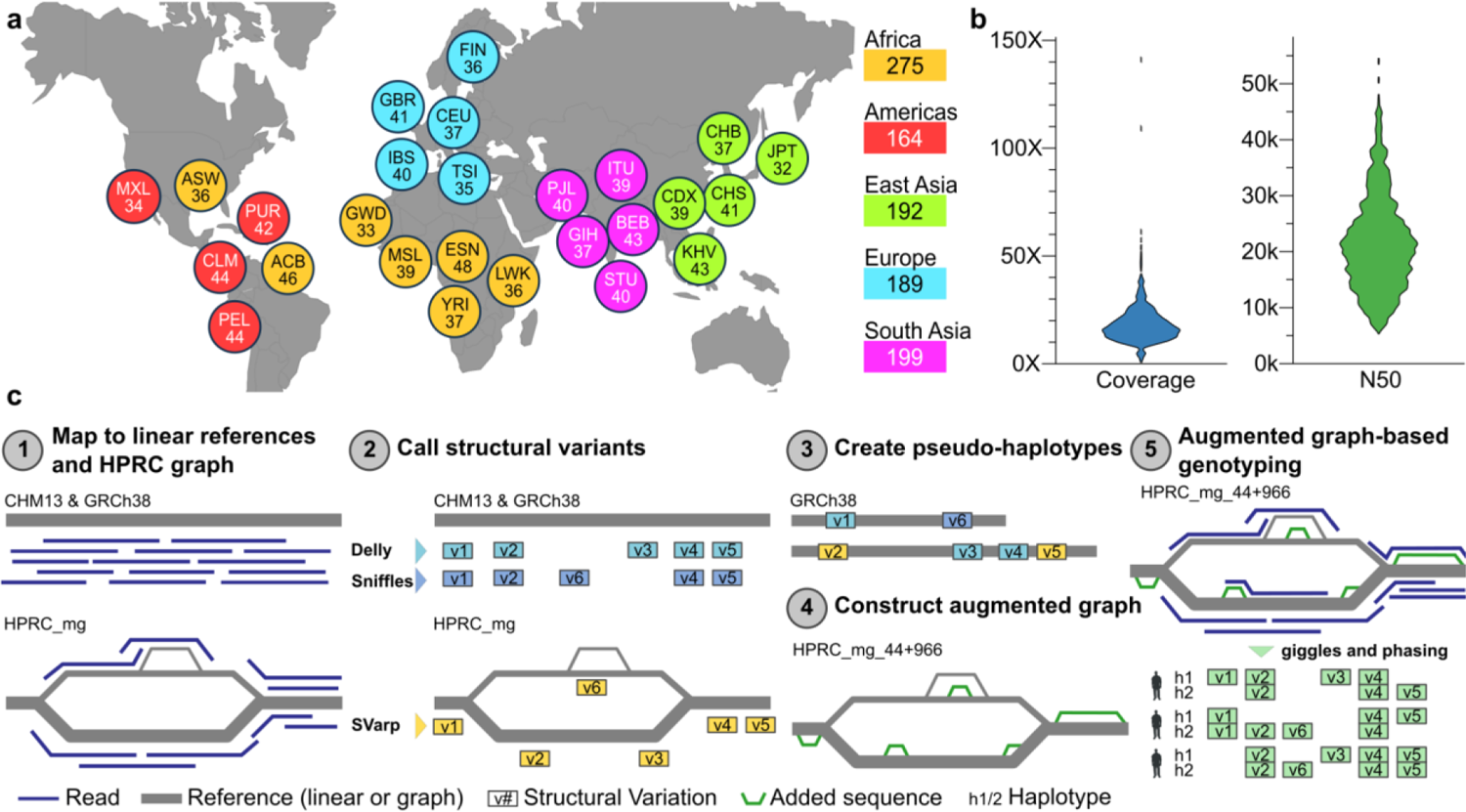
Long-read sequencing and SV analysis by graph augmentation (SAGA) framework. **a**) Breakdown of self-identified geographical ancestries, for 1,019 long read genomes representing 26 geographies (*i.e.*, populations) from five continental regions. The three letter codes used are equivalent to those used in the 1kGP phase III,^29^ and are resolved in **Table S2**. **b**) Long-read sequencing coverage per sample, expressed as fold-coverage (left), and ONT N50 read length in basepairs (right). **c**) Schematic of the SAGA framework devised for pangenome graph-aware discovery and genotyping of SVs, utilising a pangenome graph-augmentation approach.

While the HPRC recently achieved considerable advancements in pangenome graph construction and its utilisation harnessing short-read data^20,35,36^, there remains a gap in methods that leverage long reads for SV discovery and genotyping within a graph context. Addressing this gap, we devised the <u>S</u>V <u>A</u>nalysis by <u>G</u>raph <u>A</u>ugmentation (SAGA) framework (**Fig. 1c**), which capitalises on the advantages of long reads for a pangenome-aware approach to SV analysis. Outlined below, SAGA integrates read mapping to both linear and graph references, SV discovery in a pangenome context, and the assembly of pseudo-haplotypes followed by reference pangenome graph augmentation and genotyping – to enable graph-aware SV discovery and genotyping at a population-scale.

#### Mapping onto linear and graph genome references

As the first step of the SAGA framework, we aligned the long read data against both linear and pangenomic references, namely GRCh38^37^, the linear Telomere-to-Telomere (T2T) consortium reference (CHM13)^38^, and the pangenome minigraph reference from the HPRC (denoted HPRC_mg below)^20^ (**Methods**). We selected HPRC_mg as it comprehensively represents SVs while omitting SNPs and small (<50bp) insertions/deletions (InDels)^20^, thus comprising a compact graph structure facilitating analyses at the scale of a thousand long-read genomes. Comparative analysis of the alignments shows that HPRC_mg and CHM13 outperform GRCh38 in terms of average mapping identities by more than 0.5%, and that the HPRC_mg reference contains a more comprehensive collection of mobile element insertions and deletions (**Fig. S3**). Utilising CHM13, we find that on average 93.6% of the genome exhibits a coverage of 5× or more in each sample (**Fig. S4**). Haplotype phased SNPs and short (<50bp) insertions and deletions (InDels) that we extracted directly from the ONT reads (**Methods**) show excellent concordance with phased SNPs previously identified in the 1kGP cohort^3^, as evidenced by median switch error rates of only 0.69% seen in children from the six parent-offspring-trios included in our dataset, and 1.32% for unrelated (and parental) samples (**Fig. S5**).

#### Linear reference-based SV discovery

As the second step of SAGA, to allow for comprehensive SV callset construction, we applied long-read based SV callers tailored to linear reference genomes. Specifically, we utilised Sniffles^22^ and Delly^39^ version 1.1.7 optimised for long-read analysis, by applying both tools to the GRCh38 and CHM13 assemblies, respectively. Following callset integration across the respective linear references (**Methods**) this process resulted in an average number of 15,301 and 21,529 SVs per sample for Sniffles and Delly, respectively.

#### Graph genome-guided SV discovery

To complement these linear reference-based SV callers, we additionally harnessed HPRC_mg and the novel SVarp graph-aware SV caller^40^ (**Methods**) to allow for the discovery of SVs in haplotype contexts under-represented in CHM13 and GRCh38. We engineered SVarp to first discover SV signatures from graph-aligned reads, and then perform local long-read assembly to reconstruct ‘SV sequence contigs’ (svtigs). To maximise the accuracy of svtig assembly, we applied SVarp to the subset of 967 ONT genomes previously sequenced to high coverage with short reads^3^. This allowed efficient haplotype assignment (“haplo-tagging”) of 69.9% of the ONT reads using previously phased SNPs. Utilising these haplo-tagged read information, SVarp constructed on average 1,145 svtigs per sample, which we interpreted as SVs not previously represented in HPRC_mg (**Fig. S6**).

#### Construction of pseudo-haplotypes and graph-augmentation

To allow for non-redundant SV callset integration and to enable downstream analyses, the SAGA framework comprises a workflow to augment pangenomes by incorporating additional bubbles^20^ into the graph representing new SV sequences. This workflow comprises two key steps: pseudo-haplotype construction and graph-augmentation (Fig. 1c, steps 3 and 4). The pseudo-haplotype construction step generates chromosome-wide, haplotype-like sequences incorporating the discovered SV alleles in a non-overlapping manner (**Methods**). The graph-augmentation step uses the minigraph algorithm^41^ to integrate these pseudo-haplotypes into the original graph, giving rise to new bubbles at genomic loci previously not considered for SVs. By executing both steps, we augmented the HPRC_mg graph originally representing 44 genomes with pseudo-haplotypes derived from the abovementioned 967 long-read genomes. This process yielded the ‘HPRC_mg_44+966’ pangenome, which represents SVs from 1,010 individuals, taking into account that one sample from our sample set (HG01258) was already contained in the original HPRC_mg graph (**Fig. S**7**, Fig. S8**). HPRC_mg_44+966 comprises a total of 220,168 bubbles, considerably increasing SV representation compared to HPRC_mg which comprises 102,371 bubbles (**Fig. S9, Fig. S10**).

Out of the bubbles represented in HPRC_mg_44+966, we find that 105,744 (90%) are at least 1 kbp away from the nearest bubble in the original HPRC_mg, and thus most likely represent SV loci not previously considered in the HPRC pangenome reference^20^ (**Methods, Fig. S11**). To evaluate the quality of the augmented graph, we mapped ONT reads from HG00513, a sample not part of the HPRC_mg pangenome reference, onto HPRC_mg_44+966. This process yielded notably improved alignment metrics, with a gain of 33,208 aligned reads and an additional 152.5 Mb of aligned bases compared to aligning these ONT reads onto HPRC_mg (**Table S3**). This suggests that HPRC_mg_44+966 provides enhanced capabilities for genome variation analysis compared to the original pangenome reference.

#### Graph-aware SV genotyping and statistical haplotype phasing

Unified SV genotypes are a prerequisite for key downstream analyses of population-scale variant callsets, including population genetic and disease association studies. Currently, tools for constructing probabilistic SV genotypes from long-reads harnessing genome graphs are lacking. We incorporated such functionality as the fifth and final step into the SAGA framework, by devising Giggles^42^, a genotyping tool that harnesses graph-aligned ONT reads for SV genotyping. We applied Giggles (version: 1.0) to the set of 967 samples of HPRC_mg_44+966 for which our study generated ONT data, yielding a multi-sample VCF comprising genotypes for 167,291 primary SV sites after filtering (**Methods**).

Consistent haplotype-phasing elevates the value of SV resources, facilitating allele-specific analyses and detailed examination of haplotype blocks, and enhancing their utility as variant references^4,29^. We hence utilised a CHM13 haplotype reference panel recently generated based on SNP calls from 1kGP short-read data^3,43^ to carry out statistical phasing of each genotyped SV with SHAPEIT5^44,45^ (**Methods**). We prefiltered genotyped SVs to the 908 samples present in the haplotype reference panel^45^, set genotypes with a quality less than 10 to ‘missing’, and subsequently dropped all SV sites with allele count zero. We find that 164,571 (98.4%) of the genotyped SV sites are successfully phased through SHAPEIT5, which comprise our final integrated SV callset. We confirmed that allele frequencies in the callset after phasing are in excellent agreement with the primary genotypes before phasing (**Fig. S12, Fig. S13**). Our final callset includes 65,075 deletions, 74,125 insertions and 25,371 ‘putatively complex’ SV sites, where both the reference and alternative allele are larger than 1bp. Analysis of the unified genotypes show that out of all 164,571 SV sites, 148,794 (90.4%) are bi-allelic, with the remaining 6,528 representing multi-allelic SV loci typically residing in more repeat-rich areas of the genome.

#### Resource quality assessment

The quality of an SV callset is fundamental for its utility as a genetic reference, prompting us to utilise multifaceted approaches for resource quality assessment. To estimate callset accuracy, we subjected our deletion calls to intensity rank sum testing, using SNP microarray data generated in 1kGP samples^29,33,46^. We estimate a false discovery rate (FDR) of 8.06% for biallelic deletion sites, which improves to 6.97% in high-confidence regions of the genome (**Methods**), suggesting excellent SV callset quality (**Fig. S14**). To estimate the FDR for insertions, we augmented the CHM13 linear reference genome with each insertion, and then mapped sequences from compacted de Bruijn graphs generated from short reads^3^, yielding an FDR of 11.40% (8.43% in high-confidence regions) (**Fig. S15**).

We also examined the primary SV genotypes comprehensively. We used data from the six parent-offspring-trios contained in our resource to infer ‘Mendelian inconsistencies’, which could indicate either *de novo* SV formation events or genotyping errors. The average rate of such inconsistencies for biallelic SVs is only 3.87% for deletions, 4.44% for insertions, and 4.10% for putatively complex SV sites, implying high genotypic accuracy (**Table S4-S12**). For multiallelic sites the average Mendelian inconsistency is 15.05%, reflecting previously reported challenges in genotyping this variant class^4,29^. These estimates align with strong performance against Hardy-Weinberg equilibrium checks (**Fig. S16, Fig. S17**). When compared to short read based analysis of the 1kGP sample set^3^, our SV sites intersect 69.5% of insertions and 64.9% of deletions called from Illumina reads in these samples (**Fig. S18**). At these intersecting SV sites, we estimate a genotype concordance of 98.7% for deletions (non-reference genotype concordance: 77.6%) and 96.8% for insertions (non-reference concordance: 79.0%) when compared to Illumina based genotypes. Finally, using Yak^47^, we estimate an average quality value (QV) score of 37.6 for the haplotype blocks constructed with SHAPEIT5 when compared to sample-matched compacted de Bruijn graphs (**Fig. S19**). These scores, notably, are on par with QV scores previously derived from Pacific Biosciences continuous long read (CLR) genomic assemblies^7^, implying high resource quality. Taken together, these benchmarking results establish the high sensitivity, specificity and genotype accuracy of our SV resource.

### The long-read based SV landscape in genomes with ancestries from 26 populations

#### Landscape of SVs in 1kGP samples

Encouraged by these quality metrics, we conducted a comprehensive analysis of the ONT sequencing based SV landscape in 1kGP samples. We first analysed the degree to which the SV callset grows cumulatively after each new genome is added (Fig. 2a**, Fig. S10)**. We observe pronounced SV callset saturation effects when adding new samples to the resource^5^, with growth saturation being more limited for SV singletons, consistent with a large pool of rare SV alleles present in the 26 ancestries part of our resource. The addition of samples with AFR ancestries disproportionally increases SV yield, consistent with AFR exhibiting the highest levels of genetic variation among all humans^5,29^.

**Figure 2:**
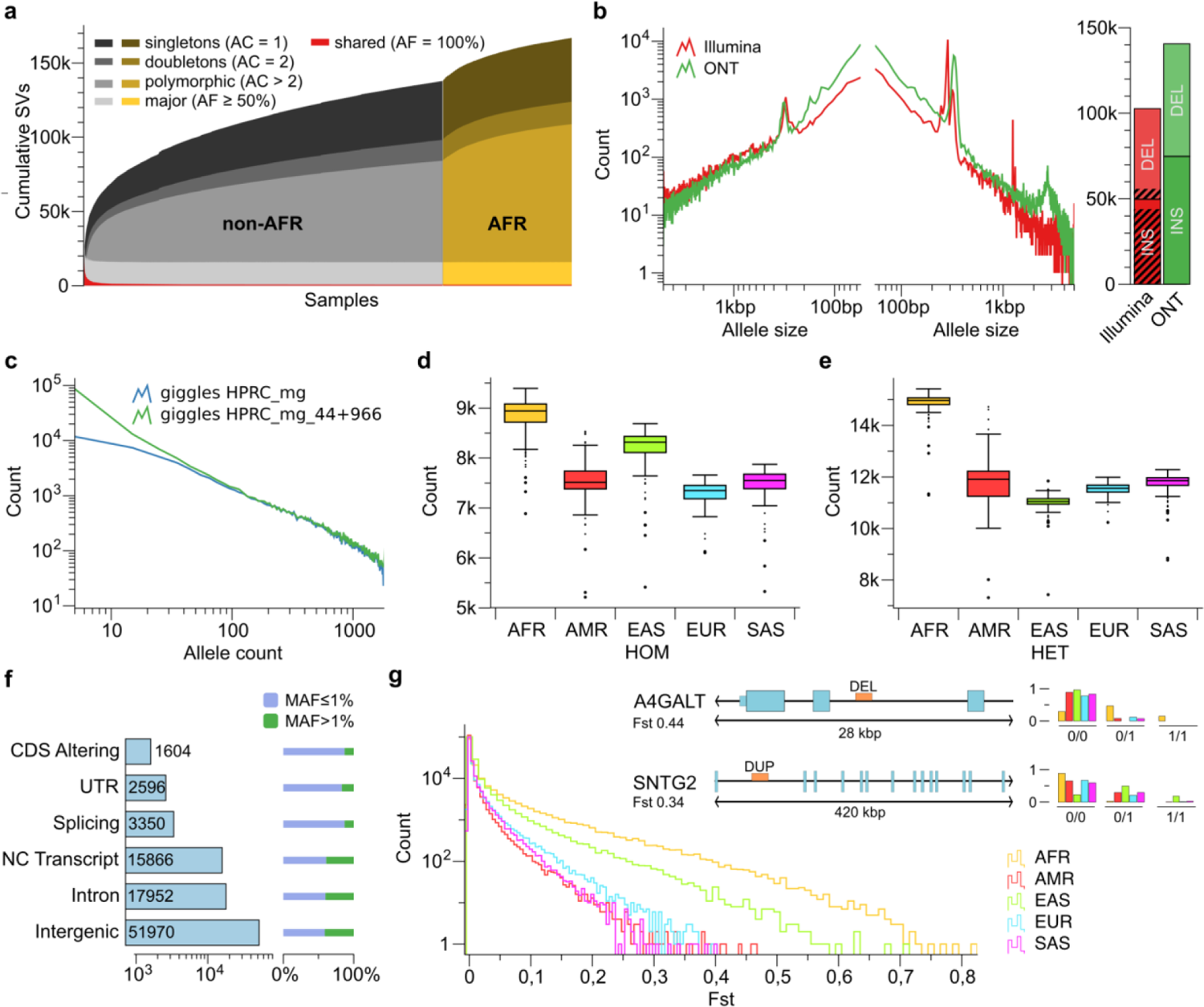
Callset properties and SV landscape across different geographical ancestries. **a**) Cumulative number of unique SVs when adding individual long-read genomes, from left to right. The rate of SV discovery slows with each new sample added. Colours denote singletons, doubletons, SVs seen with an allele count of >2 (‘polymorphic’), as well as major and shared alleles. **b**) Left: SV length distributions in population-scale 1kGP SV callsets in basepairs, versus the cumulative count of SV sites, with a comparison of ONT sequencing (‘ONT’; *N*=967 samples) to the high-coverage short-read based analysis of the 1kGP cohort^3^ (‘Illumina’) subsetted to the same (*N*=967) samples. Right: The 1kGP ONT callset is outnumbering short-read based SV calls both for deletions (DEL, upper sections) and insertions (INS, lower sections). SVs previously unresolved by their sequence, using short reads^3^, are depicted by shaded areas. **c**) SV allele count (x-axis) relative to the count of SV sites (y-axis), constructed by genotyping the original HPRC graph (HPRC_mg) and the augmented graph (HPRC_mg_44+966) using Giggles. Consistent with the considerably smaller panel size employed, HPRC_mg under-represents rare alleles. **d**) Homozygous SV count, and **e)** heterozygous SV count, per sample stratified into self-identified geographies. **f)** Impact of SVs on distinct genomic features. SVs affecting sequences of genes occur at lower MAF compared to non-coding regions. **g)** Histogram of SV Fst values for all five continental populations. Inset upper-right: deletion and duplication in the intragenic space of two medically relevant genes exhibiting strong differentiation in AFR and EAS samples, respectively.

Unlike prior population-scale surveys of 1kGP samples based on short read sequencing^3,4,30^, our resource captures SVs along their whole size spectrum, which includes 50 to several hundred basepair size SVs that are poorly captured^6,7^ by Illumina reads. Consequently, the median SV count per sample is 23,969 for AFR samples (19,297 for other ancestries, non-AFR) in our resource, more than doubling SV numbers when comparing to the most recent short-read based 1kGP study^3^ subsetted to the same samples (AFR: 9,963; non-AFR: 8,540). The proportionally largest gain compared to short read sequencing is seen for insertions, which our SV callset captures with increased sensitivity along the entire variant size spectrum (Fig. 2b). By comparison, large deletions are detected with a slightly higher abundance when using short reads^3^ (Fig. 2b), affirming the previously reported high sensitivity of short reads in the detection of large deletions via read depth analysis^6,7,48^.

#### A large resource of fully resolved SV allelic representations

Since short-read based surveys of the 1kGP sample set have utilised methods that discover SVs using inferential techniques, including by paired-end and read depth analysis^18,49^, SV allelic sequences have remained largely unresolved^3,4,30^. This shortcoming is particularly pronounced for insertions exceeding the Illumina read length, a large fraction of which lack allelic sequence resolution^3,4,30^. In contrast, our resource consistently provides sequence-resolved SV allelic representations, capitalising on the length of the ONT reads. We find a more than 10-fold increase in insertion sites with fully sequenced-resolved SV alleles compared to the most recent short read based 1kGP study, increasing from 5,609 to 74,125 in the same samples (Fig. 2b, bar chart). For deletions, which can be resolved via split-read analysis in short-read data^3,4,30^, our resource increases the count of nucleotide-resolved SVs from 46,895 to 65,812. These advancements underscore the potential of our resource to enhance the accuracy of SV analysis and variant interpretation, addressing a key prior limitation of population-scale SV studies.

#### Rare SV alleles

A more detailed analysis of the allele frequency spectrum in our resource provides a contrasting perspective relative to the original HPRC_mg graph^20^. Our dataset encompasses a broad spectrum of SV alleles, from common to very rare (Fig. 2c**, Fig. S20, Fig. S21**). This translates into a notable enhancement in capturing SV diversity over HPRC_mg, which, while comprehensively comprising common alleles, underrepresents those with an allele frequency (AF) of 2% or less (Fig. 2c). Further stratifying SVs based on their presence in the original HPRC assemblies verifies that as expected, the vast majority of rare SVs stem from genomes newly incorporated into the dataset, which were integrated into the augmented graph via pseudo-haplotypes (**Fig. S22**). This underscores that our SV callset represents an inclusive resource of SV alleles spanning a wide allele-frequency spectrum, vastly enhancing SV allelic representations across diverse human populations.

#### SV characteristics by self-identified continental ancestries

Facilitated by the diversity of samples captured, we investigated the characteristics of SVs in our dataset stratified by self-reported geographical ancestries from among the five continental groups – AFR, AMR, EAS, EUR and SAS. Most SVs occur with low frequency (59.3% have a minor allele frequency (MAF) < 1%). While these rare SVs are typically specific to individual continental groups, at an AF ≥ 2.5%, the majority of SVs are detected in samples from at least two continents (**Fig. S20**). Multiallelic SV loci generally have a higher propensity to be shared across continents than biallelic SVs, with the majority of multiallelic SVs with AF ≥ 1.5% shared across continents (**Fig. S23**), potentially explained by the recurrent rearrangements these latter loci experience. Furthermore, we find that the relative increase in SVs in AFR samples compared to other ancestries is more pronounced for heterozygous than for homozygous SVs, consistent with AFR samples displaying distinct SV site saturation statistics compared to non-AFR samples^5^ (Fig. 2a, 2d, 2e, Fig. S24).

Unlike previous 1kGP studies that have placed a strong focus on the characterisation of sequence-resolved deletions^3,4,30^, our findings reveal comparable population characteristics between SVs primarily identified as deletions and insertions (**Fig. S20)**. Measuring the degree to which SVs are in linkage disequilibrium (LD) to nearby SNPs, we find that 62.6% of deletions and 62.9% of insertions with at least 1% MAF are in LD with nearby SNPs (*r*^2^ ≥ 0.5) (**Fig. S25, Fig. S26)**. These metrics increase to 89.8% (deletions) and 91.7% (insertions), respectively, in high-confidence regions of the genome (**Methods**) that tend to be more depleted of repeats, and as such may be less likely to be subject to recurrent mutation or gene conversion events^50^. Additionally, we find that larger SVs, irrespective of being insertions or deletions, are significantly rarer in the population than smaller SVs (**Fig. S27**), a trend previously detected only for deletions^4^. This could be attributed to stronger negative selection against larger SVs, which are more likely to comprise, or disrupt, functionally relevant sequences. Consistent with this notion, evaluating the impact of SVs on distinct genomic features we observe a highly significant reduction (*P* < 7.4e-235; KS-test) in the MAF of SVs affecting functionally relevant elements (Fig. 2f).

#### Geographical stratification of SV alleles

Utilising our SV callset for a principal component analysis (PCA), we observe groupings of samples corresponding well with the donors’ self-reported geographic ancestries, consistent with prior short-read based 1kGP studies^4^ (**Fig. S28**). This is similarly reflected in an SV-based admixture analysis based on self-identified geographical ancestry (**Fig. S29**). Employing fixation indices (Fst) to quantify population differentiation, by comparing samples from each superpopulation against the remainder, the allelic diversity of AFR (followed by EAS) is clearly observable (Fig. 2g). Investigating single sites, we find evidence for differentiation (Fst > 0.2) for several thousand SVs – 6,427 for AFR, 106 for AMR, 1,673 for EAS, 295 for EUR, and 96 for SAS (**Table S13**). By intersecting these sites with the Genome in a Bottle Consortium’s list of medically relevant genes^51^, we find 105 candidate population-stratified SVs (affecting 61 distinct genes) of potential interest for disease studies (**Table S14**). Examples are a deletion and duplication affecting the intragenic regions of *A4GALT* (MAF 14%) and *SNTG2* (MAF 18%), enriched in AFR and EAS, respectively (Fig. 2g). We further note a complex SV near *LAMB1* exhibiting enrichment in AMR, and particularly among samples with self-identified ancestries from Peru in Lima (PEL) (**Fig. S30)**. Additionally, we observe a strong correlation between 1 Mb window averaged SV-based and SNP-based Fst values (Pearson *p* < 4.0e-16), deviations of which potentially serve as indicators for genomic areas with SV-driven differentiation. Requiring a confidence level of more than 5 standard deviations (*P* < 0.6e-6) we find 11 regions (AFR: 2, AMR: 3, EAS: 3, EUR: 1, SAS: 2) in which the differentiation is likely to be SV-driven (**Table S15**), including a deleted region near gene *ARHGAP24* with the highest MAF of 27.3% (Fst=0.31) seen in individuals with self-identified ancestries from the Mende in Sierra Leone (MSL).

### Detailed resolution of the spectrum of SV polymorphisms classes

#### Systematic annotation of SVs into distinct classes

Capitalising on the sequence-resolved allelic representations in our dataset, we conducted a detailed analysis of the genomic SV content by variant class. We built the SVAN computational pipeline to systematically group SVs into distinct classes, by utilising complete allelic representations and genomic annotations as an input (**Methods**). This is of particular relevance for distinct classes of sequence insertion and duplication, which have remained underexplored in studies dealing with diversity panels owing to difficulties in characterising these SVs classes using short reads^18,30^. Out of the 75,337 primary insertions, SVAN classifies 71,354 (94.7%) into a particular SV class, with only 3,983 remaining unclassified, offering detailed resolution of the spectrum of SV polymorphism classes in the human genome.

#### VNTRs and duplications

SVAN classified 26,904 (35.7%) of the primary insertion calls, and 16,338 out of the 66,237 primary deletion calls, as variable number of tandem repeat (VNTR) expansions, a class of variants with relevance to human diversity and disease^52,53^ that frequently escapes detection using short reads^52^ and thus has been under-represented in prior 1kGP studies (Fig. 3a). Out of the primary insertion calls classified as VNTRs, 26,060 (96.9%) represent simple VNTRs with just one motif, whereas 844 (3.1%) are complex VNTRs involving two or more repeat motifs. Overall, the VNTRs in our dataset comprise 15,867 distinct motifs, with motif length ranging from 1 bp to 481 bp (median 18 bp) and the length of the respective expansion/contracting ranging from 50 bp to 19.6 kbp (median 96 bp).

**Figure 3:**
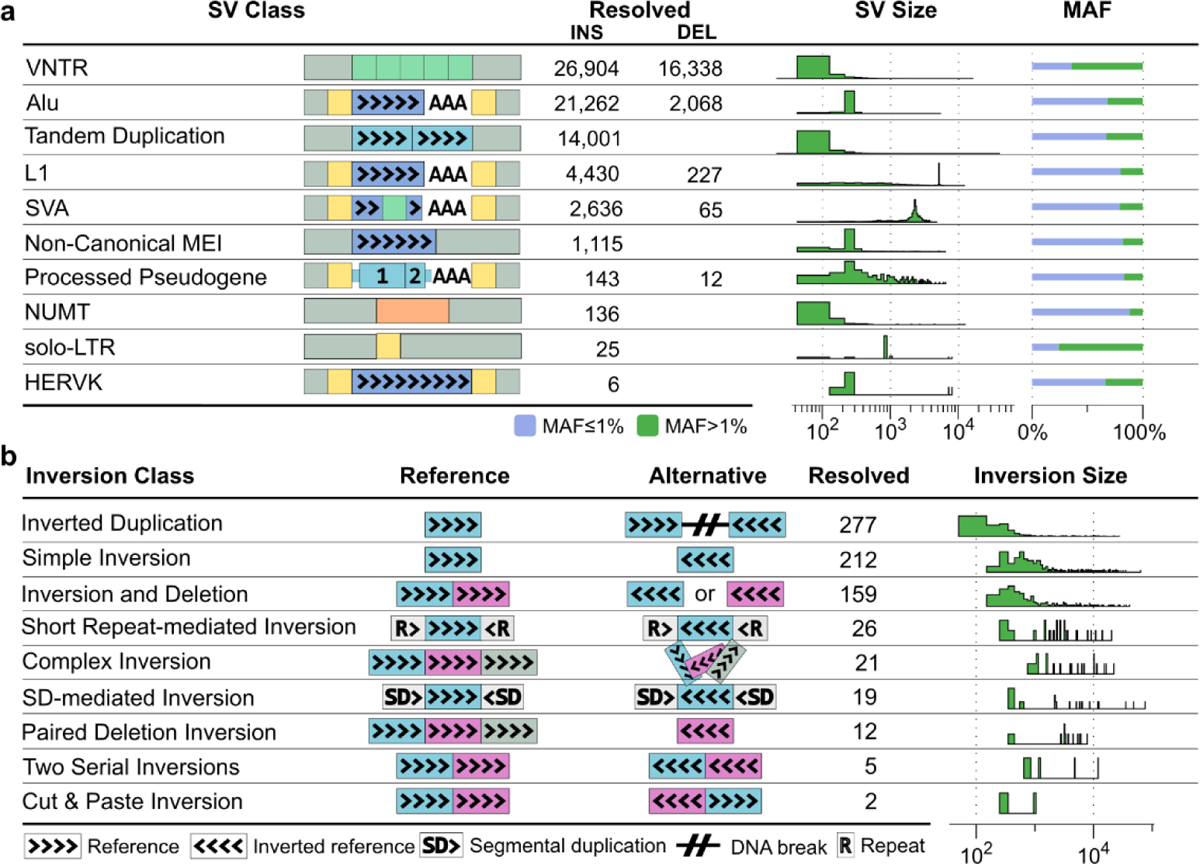
Prevalence of distinct SV classes in our long-read data resource. **a)** Polymorphic VNTRs, duplications and insertion classes classified by SVAN. The SV class is denoted by label and ideogram, the number of resolved class members is given for insertions (INS; insertion relative to the reference) and deletions (DEL; deletion relative to reference), in addition to size distribution and percentage of low and high frequency alleles. VNTR sequences depicted using green boxes. Target site duplications flanking retrotransposition insertion events and long terminal repeats (LTRs) flanking ERVK are represented as yellow boxes. Numbers for interspersed and complex duplications are shown in **Table S17**. Alu, L1 and SVA refer to canonical retrotransposition events per each of these MEI classes. **b)** Inversion classes identified. For each class, the reference structure is contrasted with the alternative allele structure, the number of resolved inversions is displayed, and the inversion size distribution is shown in logarithmic scale.

Notably, we find that VNTRs show a distinct allele frequency distribution than other SV classes initially categorised as insertions, with a pronounced tendency towards more common alleles (**Fig. S31**). When combining VNTR contractions and expansions, we observe that 64% of VNTR sites have MAF > 1%, a highly significant contrast to the 32% we detect at non-VNTR SV sites (*p* < 2.2e-308, two-tailed Fisher’s exact test). The excess of common alleles suggests repeated SV formation at these VNTR sites, which echoes patterns of SV recurrence in human populations recently identified for large inversions^54^, and is consistent with high mutational rates previously measured at VNTR loci^55^. Furthermore, we find that 15,509 (20,6%) of the primary insertion calls represent simple duplications. By assessing each duplication with respect to its source sequence we find that out of these 15,509, 14,001 (90.3%) represent tandem duplications, a class of variation considerably more abundant than inverted duplications (227; 1.5%). Furthermore, smaller subsets of the classified duplications are interspersed inserting into remote regions of the genome (1,214, 7.8%), or comprise complex events (67, 0.4%) that contain duplicated sequences both in forward and reverse orientation relative to the reference.

#### Mobile elements

We next turned our attention to numerous distinct classes of insertion classified by SVAN, most of which are the product of the activity of mobile elements (Fig. 3a). These include 28,358 non-reference mobile element insertions (MEI), including 21,262 Alu, 4,430 L1, and 2,636 SVA insertions, an increase of 7% (for Alu), 132% (L1) and 124% (SVA), respectively, over a recent 1kGP study pursued using short reads when subsetted to the same samples^3^. We further identify 2,360 reference MEIs primarily called as deletions (classified into 2,068 Alu, 227 L1 and 65 SVA events, respectively), a 4.4-fold increase compared to short read based analysis in the same donors^3^. Most of the detected MEIs (29,603/30,718; 96.4%) show distinctive hallmarks of retrotransposition^56,57^, including a target site duplication (TSD) ranging from 1-31bp in size, and a poly-A tail at the 3**′** end of the inserted sequence annotated by SVAN. A smaller subset of 1115 events represent MEIs that either lack poly-A tails, are truncated on the 3**′** end, or represent composite sequences spanning more than one retroelement family, and are thus classified as ‘non-canonical’. Missing TSDs, poly-A tails and 3**′** truncation could be due to sequence changes occurring after the MEI event or an endonuclease independent mechanism of integration^58^. Amongst the 4,202 non-reference L1 elements, 469 represent full length L1 elements, whereas 3,733 are truncated or inverted in their 5**′** end^54,59^. We also identify 143 (0.19%) non-reference processed pseudogenes, as well as twelve reference polymorphic processed pseudogenes, in our dataset. These carry a TSD and a poly-A tail, suggesting they have arisen via the trans-mobilisations of cellular mRNAs through the L1 machinery^60,61^. The majority of these (81, 56.6%) are monoexonic, with the remaining pseudogenes containing from 2 up to 21 exons. Additionally, we find evidence for human endogenous retrovirus (HERV) activity, with 2 full length HERVK, 4 truncated HERVK and 25 insertions of long terminal repeats (LTRs) likely resulting from HERVK integration followed by recombination^62^.

#### Nuclear mitochondrial DNA segments (NUMTs)

Finally, using SVAN we identify 136 insertions of NUMTs in our SV callset. NUMTs occur on all chromosomes (**Fig. S32**), with a median length of inserting mitochondrial fragments of 125 bp, similar to the NUMT size distribution previously reported in short read data^63^. 9 NUMTs are larger than 1 kb in size, including a near full-length mitochondrial insertion (14.8 kb) comprising 90% of the full mitochondrial length, on the q-arm of chromosome 9 (q22.31) that we identify in one carrier (**Fig. S32**).

#### Inversions

Inversions remain one of the most understudied SV classes owing to their balanced nature preventing detection through read-depth analysis, and with the common presence of flanking repeats additionally complicating their discovery^4,54,64^. While our initial SAGA-based joint SV callset does not comprise inversions, the notable N50 read length of this ONT dataset led us to explore its potential for inversion discovery. Our initial exploration revealed that inversions frequently escape automated detection in long reads owing to inaccuracies in read mapping particularly affecting inversions smaller than 1 kb, which appear as clustered mismatches in minimap2 alignments (**Fig. S33**). To allow capturing these inversions, we examined strategies for inversion discovery by simulating inversions of varied sizes (see **Note S1**). We generated simulated ONT reads from these augmented genomes, using SURVIVOR^65^, mimicking the sequencing coverage in our resource. Evaluating different aligners and SV callers led to two observations: firstly, we find that NGLMR^22^ provides superior read alignment capabilities at inversion sites. Secondly, we find Delly to be the most effective inversion caller when applied to NGLMR aligned reads (**Fig. S34**). We devised an inversion discovery workflow (**Methods**) leveraging these insights, followed by verification of each candidate inversion locus using alignment dotplots (**Fig. S35**), to construct an inversion callset across the 1,019 samples from our study.

Overall, this workflow identified 733 inversions, comprising 134 SVs identified through our custom workflow, 317 detected with Delly based on minimap2 alignments, 40 found via Sniffles, and 242 events primarily defined as insertions but later reclassified as inversions through SVAN. We conducted detailed analyses of these inversions, focusing on their size, complexity, and overlap with prior callsets^54^. We find that inverted duplications represent the most common inversion class, with 277 events detected; these display relatively small median length of 284 bp (Fig. 3b**, Fig. S36**). Additionally, we find 257 balanced (or “simple”) inversions with a median length of 1,565 bp, which are further broken down into 45 inversions bordered by various repeat classes in inverted orientation based on alignment dotplot analysis, implying formation through homology-directed repair (HDR) processes or non-allelic homologous recombination (NAHR)^66^. These flanking inverted repeats include Alu (*N*=10 inverted element pairs), L1 (8 pairs), long tandem repeat (LTR) (4 pairs) and low-complexity repeats (4 pairs), as well as segmental duplications (SDs) up to 75.5 kb in length in inverted orientation (19 pairs). We do not find inversions flanked by hundreds of kbp of SD sequence, in spite of their commonness in the genome, consistent with these SVs requiring long-read lengths in excess of the SD length for their detection^54^. Strand-seq^54,67^ or hybrid genomic assemblies using a combination of high-accuracy and ultra-long reads^68,69^ will be required to resolve this SV class at the population-scale in the future.

Prior studies of inversions have associated this SV class with an abundance of complex events exhibiting multiple SV breakpoints^4,16,54^. Facilitated by the ONT read length, we find a diversity of complex inversion patterns (Fig. 3b). These include 159 inversions with an adjacent deletion, and 40 more complex inversion structures, the latter of which comprise 5 SVs characterised by two serial inversions, 2 instances of “cut & paste” inversions with the locus structure characterised by an excised, inverted, and inserted segment into a nearby location, 12 inversion sites exhibiting two flanking deletions, and 21 particularly complex variants exhibiting a multitude of SV breakpoints, the structure of which could not always be resolved. These inversion structures suggest SV formation by a DNA replication associated process, such as microhomology-mediated break induced replication (MMBIR)^70^.

### Insights into polymorphic L1 and SVA transductions

MEI transductions occur when a full-length mobile element transports adjacent genomic sequences during retrotransposition^9,10^. This process can effectively generate the duplication of unique sequences in the genome situated in 5′ or 3′ relative to the source mobile element. Polymorphic transductions were previously reported in conjunction with polymorphic L1 and SVA elements^5^, as well as in association with somatic L1 activity in cancer^71,72^. Harnessing our basepair resolved dataset, we investigated polymorphic transduction events amongst all sequence-resolved MEI sequences. We find 880 transductions in total in our primary callset – with 497 (11,2%) out of the 4,429 primary polymorphic L1 insertions, and 382 (14,5%) of the 2,636 primary SVA insertions exhibiting a transduction (**Table S16)**. In addition to these transduction events containing a companion mobile element segment (denoted ‘partnered transductions’), an additional set of 229 transductions are highly truncated resulting in the integration of the transduced sequence alone (denoted ‘orphan transductions’). These numbers of transduction events exceed prior data on polymorphic transductions reconstructed to full sequence length by 5.3-fold (n=94, Human Genome Structural Variation Consortium (HGSVC) phase II) for L1 elements^5^ and 4.9-fold for SVAs^5^ (n=77). Furthermore, while orphan transductions have previously been reported in cancer^71,72^, to our knowledge this study provides the first report of this sequence class in a population-scale sequencing dataset, underscoring that transductions represent a prevalent source of duplicative sequence polymorphism contributing to structural variation in the germline.

Motivated by the proportionally high number of novel transduction events, we pursued a more detailed analysis of full-length transduction events. Consistent with prior studies^5,9,73^, we find that the relative proportion of 3′ or 5′ transduction events differs considerably between both MEI classes. Most (90.1%; 448) of L1 transduced sequences correspond to 3′ transductions, with 5′ transductions (9.9%; 49) seen much more rarely (Fig. 4a**, Table S16**). By comparison, SVAs generate 5′ and 3′ transductions at more similar ratios, with 223 (58.5%) events detected in 5′, and 1598 (41.5%) in 3′, respectively (Fig. 4b). Additionally, while we find L1-mediated 3’ transductions to be significantly longer in size than 3’ transductions (*P*=1.1e-15; two-tailed Mann-Whitney U test), we observe no such significant length difference for SVAs (Fig. 4a). These observations indicate the existence of class-specific determinants for 5’ and 3’ transduction occurrence within mobile elements, and motivated a more detailed analysis of the respective source elements of transductions in our dataset.

**Figure 4:**
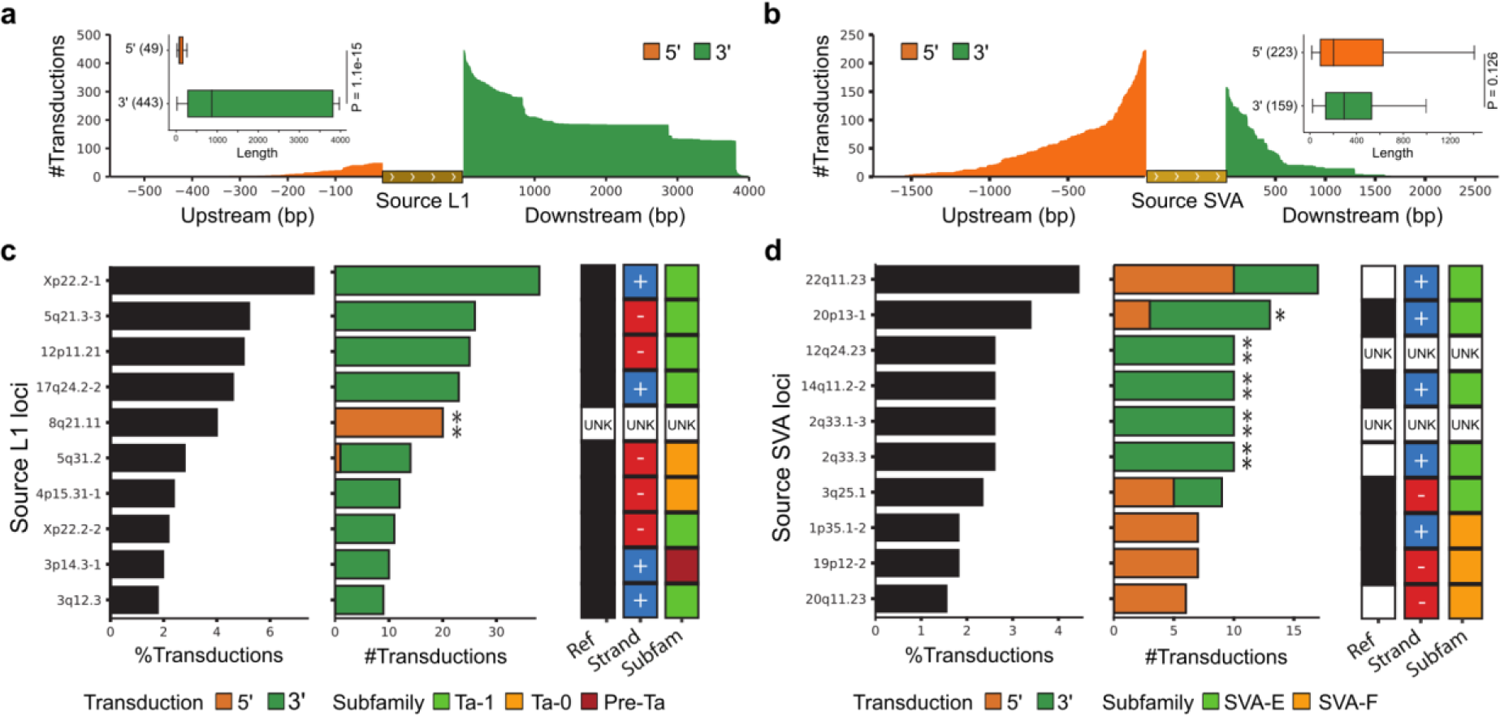
Polymorphic landscape of transductions for L1 and SVA source elements. **a-b)** Size distribution and number of 5′ and 3′ L1 and SVA mediated transductions (td) based on the analysis of flanking sequences. 5′ versus 3′ transduction length (*P*-values based on a two-tailed Mann-Whitney U test). Line, median; box, 25th to 75th percentile (inter-quartile range, IQR); whiskers, data within 1.5 times the IQR. **c-d)** Contribution relative to the total number of transductions and transduction counts for the 10 most active L1 and SVA loci. Source elements with significant bias (based on a Binomial test with P-values adjusted with the Benjamini-Hochberg method) towards 5′ or 3′ transductions are highlighted with a single asterisk (adjusted *P* < 0.05) or two asterisks (adjusted *P* < 0.005). Source loci are annotated by their presence or absence in the CHM13 reference, orientation (+ or - strand) and subfamily designation. Source loci without identified source L1 or SVA element and, therefore, with no annotation provided, denoted as “UNK”.

Specifically, we leveraged the transduced sequences as barcodes to identify their progenitor loci, revealing that L1 transductions originate from a limited set of 187 L1 source elements. Amongst these, 10 highly active source L1s are responsible for 38% (188) of all L1 transductions identified (Fig. 4c), with most (6/10) of them belonging to the youngest Ta-1 subfamily. By comparison, we detect 161 source SVA elements, with 10 loci alone mediating 26% (382) of all SVA transductions (Fig. 4d). All of these highly active SVAs belong to the recent human-specific SVA-E and SVA-F subfamilies. Interestingly, we find that for both L1 and SVA source elements the respective proportion of 5’ and 3’ transductions appears to strongly depend on the genomic locus. Out of the 188 L1 progenitors generating transductions, 159 exhibit only 3′ transductions. This includes the most active source L1 locus, a Ta-1 element residing at Xp22.2-1 on the reference, for which we find 38 transductions genome-wide – all in 3′ (**Table S16**). In contrast, the fifth most active L1 element at 8q21.11, showing 20 transductions, exhibited only 5′ transduction events (Fig. 4c). By conducting statistical testing, we uncover a highly significant bias towards transduction at the 5’-end for this L1 source element (*P=*1.98e-15; binomial test, controlled for the FDR with the Benjamini-Hochberg method). SVAs progenitors, by comparison, display a pronounced locus-specific pattern of transduction activity, with 3 amongst the 10 most active source SVAs mediating solely 5′ transductions, and 4 mediating solely 3′ transductions (Fig. 4d). Applying an 5% FDR threshold (**Methods**), we find five SVA source elements exhibiting a significant bias towards 3′ transductions, including elements at chr12q24.23, chr14q11.2-2, chr2q33.1-3, chr2q33.3 and chr20p13-1. Additionally, four additional SVA source elements, namely those at chr6p12.1, chr12p11.23 and chr20p13-2, as well as an orphan SVA element attributed to chromosome 12, exhibit a significant bias towards 5′ transduction. These data suggest strong locus-specific dependencies exist for L1 and SVA mediated 5′ and 3′ transductions in the germline.

#### SVs in a typical human genome

The overall abundance of SVs identified by SAGA compared to prior short-read based studies of the 1kGP samples motivated an analysis of the number of SVs per class seen in a typical human genome, whereby we distinguished between donors with ancestries from AFR and other donors (non-AFR). Utilising all classified SVs identified with the SAGA framework, we find a median of 8,989 deletions in AFR donors (non-AFR: 7,334), 9,921 insertions (non-AFR: 7,889) and 5,008 putatively complex SVs that cannot be directly resolved into either deletions or insertions (non-AFR: 4,048). Further resolving these SVs by class, AFR samples exhibit a median of 4,577 deletions affecting VNTR sequence (non-AFR: 3,795), 5,239 VNTR expansions (non-AFR: 4,203), 1,766 tandem (non-AFR: 1,485), 129 interspersed (non-AFR: 100) and 3 complex duplications (non-AFR: 2), 1,678 MEIs (non-AFR: 1,203), 86 non-canonical MEIs (non-AFR: 68), 1,226 deletions of mobile elements (non-AFR: 998), 6 insertions (non-AFR: 5) and 2 deletions (non-AFR: 2) of processed pseudogenes, 8 insertions of LTRs or LTR flanked HERVK elements (non-AFR: 5), as well as 4 NUMTs (non-AFR: 3) (**Table S17**).

### Breakpoint homology landscape and evidence for recurrent *Alu*-mediated SV formation

#### Breakpoint homology landscape suggests a continuum of homology-associated processes

SV breakpoint analyses enable exploring the molecular origins of SVs, ranging from homology-mediated SV formation to homology-independent DNA repair processes^4,5^. Prompted by the vast increase in nucleotide-resolved SVs over prior 1kGP studies, we comprehensively investigated SV breakpoint junctions, examining 66,198 deletions and 75,238 insertions from the primary callset, respectively. To allow for equivalent analyses between deletions and insertions, we implanted each insertion into the CHM13 reference genome, followed by an analysis of flanking sequences at the respective breakpoint junctions to identify homology (defined as ≥50 bp stretches of sequence homology) and microhomology (<50 bp; Fig. 5a**, Methods**). In examining insertions by class, we find that VNTRs and tandem duplications present extensive breakpoint homology (49.7% and 89.8%, respectively), with the homologous sequences flanking the SV frequently mirroring the inserted element in length (**Fig. S37, Fig. S38**). VNTRs typically form through processes like replication slippage, HDR, and NAHR, involving DNA sequence homology^74^. Nonetheless, like for simple tandem duplications, the emerging allelic structure of VNTRs generates homology at the SV flanks independent of the formation mechanism, thus necessitating separate consideration of both SV classes when analysing breakpoint junctions.

**Figure 5:**
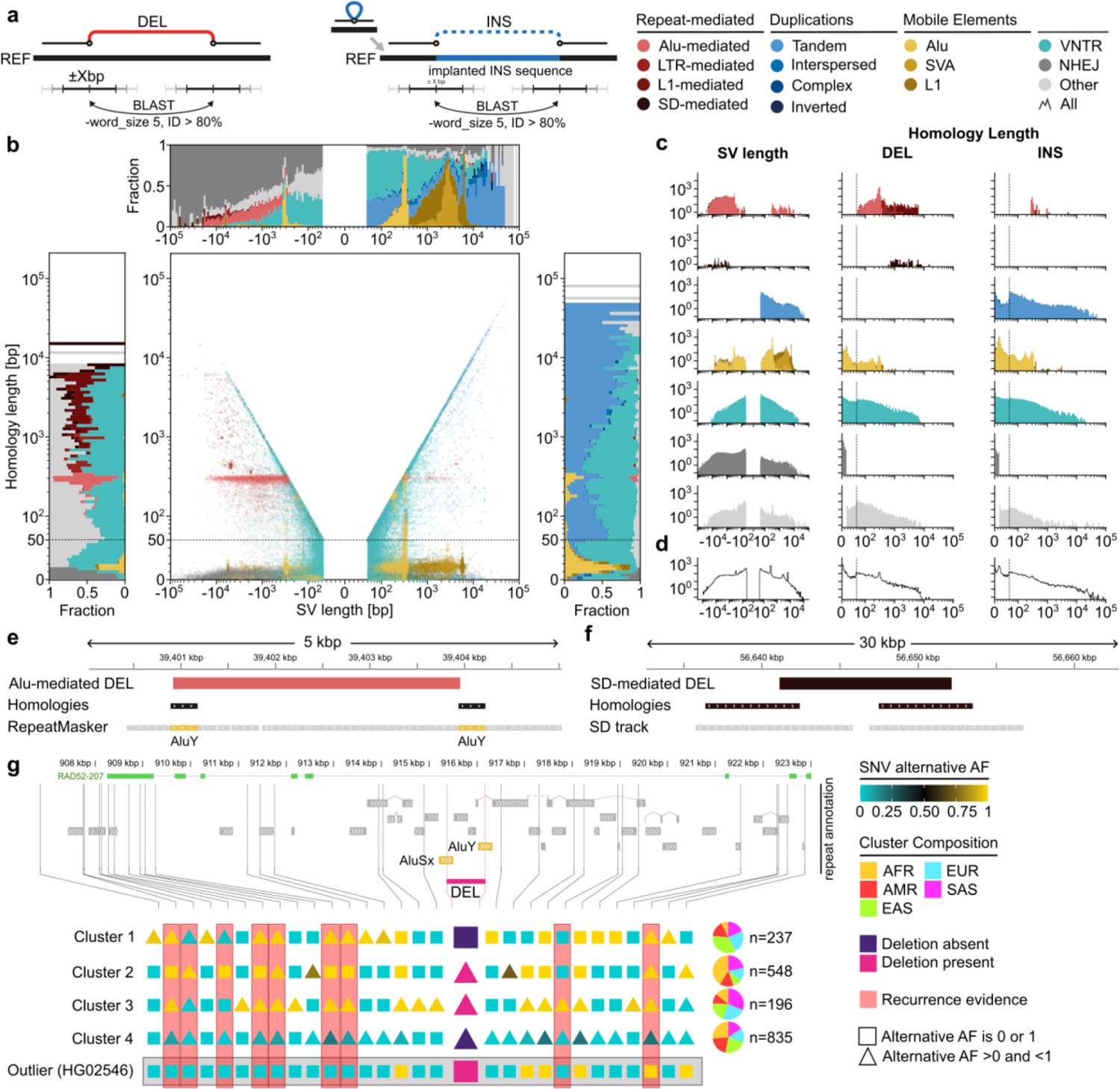
Breakpoint homology landscape and evidence for homology-mediated SV recurrence. **a**) approach for analysing homology at SV breakpoint junctions, which in the case of primary insertion calls (INS) is achieved after implanting the respective sequence-resolved SV into the CHM13 reference (REF). DEL, primary deletion call. **b)** Scatter plot of SV length versus homology length for all SVs coloured by the respective class (DELs depicted with negative length, and INS with positive length). Marginal plots show the size-binned fraction of SV classes perpendicular to both axes, depicting DEL and INS at the left and right, respectively. SVs directly flanked by paired repeats are shown in different shades of red, with SD-mediated SVs in black. SVs exhibiting microhomology ≤15 bp in length or blunt ended breakpoints are coloured in dark grey. Further colouring using SVAN annotations denotes the following SV classes: duplications (shades of blue), mobile elements (shades of yellow) and VNTRs (cyan). SVs not classified by SVAN are coloured in light grey. **c)** Histograms showing SV length and homology length separated by DEL and INS for distinct SV classes. **d)** Homology length trend lines for INS and DEL classes combined. **e, f)** Schematic view of a 3 kb Alu-mediated DEL and a 11 kb SD-mediated DEL. For visualisation purposes, scales of scatter plot axes in a) and x-axes in b) are linear up to 50 bp (representing microhomology) and logarithmic afterwards (representing homology); this split is denoted by the dashed line. **g)** A 806bp DEL at 12p13.3 mediated by an AluSx-AluY pair with evidence for recurrent formation. Clusters 1-4 were obtained from SNP based clustering of the haplotypes in a 100kbp window centred around the DEL, of which an interval of 18kbp is shown. Pie charts represent continental ancestries for each of the four clusters. For visualisation purposes, a consensus haplotype in a 20kbp window centred around the DEL is represented for each cluster. Squares are used to represent an allele frequency of 0 (yellow) or 1 (cyan) within a cluster while triangles represent allele frequencies in between. A haplotype from sample HG02546, grouping with Cluster 4, is represented in the figure as an outlier. Red bars represent SNPs indicative for recurrent deletion formation.

Leveraging the annotations provided by SVAN, our analysis of breakpoint junctions verifies that most MEIs exhibit TSDs of 10-20 bp at the respective insertion site (Fig. 5b**,c**). When analysing SVs neither annotated as VNTR, tandem duplication, or MEI, 35.0% of deletions and 33.8% of insertions exhibit homologous flanks exceeding 50 bp, indicative of SV formation through HDR processes. A considerable subset thereof – 11.1% of the deletions and 6.7% of the insertions – are flanked by ≥200 bp of homology, and are more likely mediated by NAHR^75^.

We further identify several clusters of SVs flanked by Alu, L1, and LTR elements annotated in the reference genome (Fig. 5b**,c****,e**), the formation of which is likely to be mediated by HDR as well as NAHR^76^. Amongst these, we find Alu-flanked SVs to be much more common in deletions (N=3546) than insertions (N=146). This group of SVs predominantly harbours pairs of full-length Alu elements at their flanks (89.5%), visible as a sharp breakpoint homology length peak of 295 bp in the breakpoint homology landscape (Fig. 5b). Notably, these presumably Alu element-mediated SVs display a wide distribution of SV lengths, ranging from ∼300 bp up to 25.2 kb for deletions and 9.5 kb for insertions. AluY and AluSx elements in all combinations comprise 23.5% of all SV flanking Alu pairs, with AluY– AluY pairs (8.9%) followed by AluSx-AluSx pairs (7.0%) representing the most common configurations. The usage of the different subfamilies of Alu elements at breakpoint junctions highly correlates with their counts in the reference genome (**Fig. S39**) – with the exception of the AluJ family, one of the oldest Alu families, whose members appear several times less frequently at the flanks of SV than expected by their reference genome count (e.g., with an 18-fold reduction seen for AluJb). Generally, the frequency of Alu elements at breakpoint junctions is consistent with Alu-based transposable element-mediated rearrangement representing a prevalent SV formation process^76^.

We also find a number of L1-flanked SVs, albeit with a lower abundance than Alu-flanked SVs, potentially attributed to the more ubiquitous presence of Alu elements in the genome. Like for Alu-flanked SVs, L1-flanked SVs are much more common in deletions (N=227) than in insertions (N=1). These L1-L1 pairs mediate larger SVs on average than Alu-Alu pairs (Fig. 5c), resulting in events up to 62.9 kb in size. Pairs of L1PA family members are most often represented, partaking in 76.8% of L1-L1 mediated SVs. Contrary to Alu-mediated SVs, we find L1 elements at deletion flanks to be typically truncated, with only 4.8% of L1 sequence homology-mediated deletions flanked by full length L1 elements, and with these truncated L1 pairs showing a median length of 1.7 kb.

We also find three distinct clusters of SVs of LTR flanked deletions and insertions, reflecting integrations or deletions of members of the HERVK (deletion: 40, insertion: 7), HERVH (deletion: 112, insertion: 9), and mammalian apparent LTR retrotransposon (deletion: 46, insertion: 4) families. Apart from truncated events seen in 30 cases, the homology-lengths correspond to the length of the LTRs flanking the respective integration site, with the SV length corresponding to the size of the viral integration plus one LTR. Moreover, we find 60 SD-mediated deletions (e.g. Fig. 5f), which are up to 31.8 kb in size and exhibit a homology length of up to 14.9 kb. We note that our approach is likely to exhibit decreased sensitivity in identifying SD-mediated SVs flanked by SDs larger than the read length of our study, suggesting larger SD-mediated SV events could be underrepresented.

Furthermore, we analysed 30,648 deletions not categorised into a particular subclass by SVAN and lacking homology of at least 50 bp at their breakpoints. Out of these, 2836 (9%) exhibit blunt ended-breakpoint junctions whereas 25,424 (83%) exhibit microhomology from 1-15 bp in size, indicative of SV formation by non-homologous end-joining, alternative end-joining, or MMBIR^77^. The remaining 2388 (8%) events show microhomology 16-49 bp in size, and are likely to have formed through homology-independent processes as well. We further find that inverted duplications rarely show homology at their breakpoints (9.3%), with most exhibiting microhomology of 1-15 bp at their respective breakpoint junction, suggesting a replicative formation origin of this SV class.

Lastly, we analysed the distribution of SV breakpoint homologies with increasing SV allelic length as a whole. Notably, we find that this distribution lacks appreciable inflection points, notwithstanding peaks in homology length corresponding to the sizes of mobile elements at the deletion flanks (Fig. 5c**,d**). This implies that a wide spectrum of homology-mediated SV formation processes is operating in the germline, generating variants of different classes and scales and representing a continuum of flanking homology lengths driving SV formation.

#### Evidence for recurrence of Alu-based transposable element-mediated rearrangements

Given the abundance of nearby mobile element sequences in the genome and the sequence homology they share, we postulated that occasionally SVs may form recurrently in human populations, mediated by the same flanking element pair. While precise mutation rate estimates will require larger cohorts sequenced with long reads, we conducted a semi-manual screening of Alu-mediated deletions (**Methods**), and selected a 806 bp deletion at 12p13.3 mediated by an AluSx-AluY pair to explore this phenomenon (Fig. 5g). After clustering haplotypes using SNPs in a 100kbp window centred around the deletion (**Methods, Fig. S40**), the four major emerging clusters recapitulate the geographical ancestries (Group 1: majority EAS; Group 2+4: majority AFR; Group 3: majority SAS). The deletion allele, notably, is observed in different haplotype backgrounds of groups 2 and 3, as well as in an outlier haplotype that clusters with Group 4 and is otherwise composed of sequences not carrying the deletion. Similar to an analysis previously employed to establish inversion recurrence^54^, we searched for SNPs for which haplotypes with all four combinations of both SNP alleles with the deletion being absent/present are observed. We find nine such SNPs, which are characteristic of either recurrent deletions or extensive local recombination near the deletion. Given their close proximity and association with specific haplotype groups, the most plausible explanation appears to be recurrent formation of an Alu-Alu pair-mediated deletion, with scenarios based on homologous recombination on either side of the event appearing less likely. Together with the unique SV allele frequency spectrum observed at VNTR sites as discussed above, these data support the notion of repeat-mediated SV recurrence in the human genome.

## Discussion

### An open-access resource of 1,019 human genomes sequenced with long reads

By devising the SAGA framework, we have conducted the thus far most comprehensive analysis of SVs in 1kGP samples based on long-read sequencing, generating an open data resource providing significant advancement in understanding human genetic diversity. Unlike prior population-scale SV surveys, which conducted analyses relative to a linear reference genome, our study couples linear and graph-based approaches for SV discovery, along with variant refinement and genotyping enabled by graph augmentation. Following the paradigm devised in the 1kGP^29,32,33^, our study provides proof-of-principle that intermediate coverage long-reads enable the generation of comprehensive SV catalogues at nucleotide resolution comprising common as well as rare SV alleles from populations around the globe.

We envision that our resource will serve as a basis for the construction of panels of normal long-read sequences from individuals sampled around the globe. It will be instrumental for variant pathogenicity prioritisation, serving as a foundation for the anticipated shift in disease studies towards using long read sequencing. We further envision our open resource to serve as an important benchmarking dataset for methods development in long-read variant discovery, genotyping and pangenomics^19,78,79^, facilitating an informative choice of methods for adopting nanopore sequencing in genomic medicine.

#### Implications for SV biology

Facilitated by the nucleotide resolution allelic representation of SVs in our resource, we pursued a number of analyses with implications into SV biology. The elucidation of a spectrum of homology-utilising molecular mechanisms for SV formation – enabled by long reads capturing repeat-associated SVs escaping efficient detection with short reads^5^ – provides insights into genomic architectures shaping DNA rearrangements in the population. We detected breakpoint homology lengths from 50 to thousands of basepairs contributing relatively homogeneously to the SV landscape. Prior SV studies applied sequence homology length thresholds, informed through molecular studies, to examine SV breakpoints with respect to homology-mediated SV processes^5,80^ (such as 200 bp for NAHR). While we acknowledge the importance of thresholds determined through molecular research, our data imply that a wide variety of breakpoint homology-associated SV formation processes are operating in the germline. These appear to be mediated through a continuum of homology lengths, leveraging distinct genomic repeat-contexts for *de novo* SV formation. In the future, advancements in long-read sequencing may allow for a more quantitative analysis of homology spectra at SV breakpoint junctions, and facilitate the accurate estimation of SV formation and SV recurrence rates^54^.

Leveraging the full SV allelic sequence representations, we undertook a comprehensive analysis of a diversity of classes of SVs primarily called as insertions, including VNTRs, which exhibit an unusual allele frequency spectrum indicative for mutational recurrence. We also comprehensively analysed mobile elements, which are accountable for 1.14-2.79 Mbp (median: 1.95 Mbp) of sequence variation detected per individual genome – involving MEIs, transductions, processed pseudogenes, and transposable element-mediated rearrangements driven by breakpoint homology. This represents a considerable portion of the 3.44-7.45 Mbp (median: 5.48 Mbp) of sequence variation per genome attributed to SVs in our study (**Table S18, Fig. S41**). Moreover, with respect to LINE-1 and SVA mediated transductions, our data indicates that whether the transduction affects the 3′ or 5′ end is influenced by the mobile element class as well as the genomic location of the respective source element. These findings suggest the existence of DNA sequence determinants for 3′ or 5′ transduction, respectively. It should be noted that other than in cancer genomes^71,72^, there is a possibility that the source mobile element does not co-segregate with the corresponding transduction event in the population, potentially leading to progenitor copies remaining undetected.

#### An ONT genetic diversity resource to foster SV phenotypic studies

Looking forward, we anticipate that our long-read resource will serve as a crucial tool for expanding research focused on the phenotypic implications of SVs, using data access-controlled biobanks in individual populations^81–86^. In the future, this will be particularly crucial for populations currently underrepresented in disease studies^85^, allowing to enhance inclusivity in genomic medicine. To illustrate the practical application of our data, we explored the genotyping of medically relevant genes in complex loci of the genome known to be difficult to ascertain by short read sequencing. This exploration was facilitated by the use of Locityper^87^, a tool designed to address the challenges of accurately genotyping regions that are traditionally difficult to ascertain. We analysed 265 genes identified by the Genome in a Bottle Consortium as challenging yet crucial for understanding genetic underpinnings of disease^51^, along with 20 polymorphic *MUC* family genes and the *LPA* gene, which are of particular interest due to their association with various structural haplotypes and health conditions^88,89^ (**Table S19**). Leveraging our ONT dataset, we observe a remarkable level of genotyping accuracy in the 1,019 samples of our study, particularly in regions fraught with structural complexity, considerably surpassing genotyping with short reads (**Fig. S42, Note S2**). Additionally, the haplotype data derived from our ONT resource are anticipated to facilitate enhanced variant imputation efforts, including in genomic areas previously difficult to analyse. To explore this aspect, a companion paper to this study demonstrates the utilisation of our ONT data resource for imputation and genome-wide association studies^90^. These data suggest broader value of our dataset for ascertaining disease relevant genetic variation.

#### Remaining limitations

While the data and computational methods developed in our study mark an important step forward in population-scale SV characterisation, certain challenges remain. The intermediate coverage study design^4,30^, while fostering the identification of SVs exhibiting a wide spectrum of allele frequencies, may limit the discovery power for extremely rare variants (e.g. with allele count=1), while at the same time introducing challenges in ensuring sequence consensus quality for such rare alleles in augmented graph references. Prior studies pursued by us and others as part of the HGSVC, HPRC and T2T consortia revealed the benefits of whole genome assemblies to obtain complete SV catalogues for individual donors^5,20,38^. Yet, sample numbers in these studies are presently limited, with high-quality whole genome assembly not currently scaling to cohort sizes of a thousand samples or more. Furthermore, achieving telomere-to-telomere assemblies for all human chromosomes is not yet possible in a scalable and fully automated manner, presenting an active field of research^78,91^. In the immediate future, the use of whole genome assembly in isolation is hence deemed less likely to significantly contribute to the advancement of precision medicine by facilitating the generation of comprehensive genetic variant data at a population-scale. This evaluation is further affirmed by the “All of Us” study’s^92^ choice of a future approach that incorporates intermediate-coverage long-read sequencing, thus resembling aspects of our study’s approach, albeit in a data access-restricted cohort. Additionally, future challenges include pursuing population-scale graph-based variant discovery with consideration of all small variant classes (<50bp) including SNPs and Indels. This approach is presently hampered by extensive computational demands in mapping long reads to a graph when including short sequence variants^35^, highlighting a further area for future methods development.

#### Considerations for completing the full 1kGP sample cohort using long reads

The results from our study suggest that in coming years, a coupled approach enhancing sample size at intermediate coverage – the approach employed by our study – and increasing sequencing depth on sample subsets, will represent a particularly promising middle ground to move forward. Such an approach will allow full completion of the sequence of more common structural haplotypes, while at the same time provide insights into rare structural haplotypes including those associated with diseases – enabling consideration and inclusion of large-scale clinical cohorts from a wide diversity of geographical locations. During early stages of our study, we thus coordinated sample sets with efforts piloting genome sequence assembly in smaller 1kGP sample subsets^5,20,93^, and provided open access to our data and callsets. This approach to open data sharing is guided by the principles of the 1kGP and is inspired by the potential to combine intermediate- and high-coverage techniques to advance the completion of the catalogue of human genomic variation encompassing the entire 1kGP cohort^3,29,32,33^ in the near future. It also aims to ensure that donors and the communities contributing biological samples can gain a long-term benefit from the research outputs generated^94^.

#### Final conclusions

In conclusion, our study leverages the potential of ONT sequencing, along with a graph-augmentation framework, to provide the most complete population-scale SV dataset to date, shedding light on the complex human SV landscape, SV formation processes and population-specific patterns. Our study underscores the potential of continued development of pangenomic computational tools^19,78,79^, which in the future will be facilitated with the data from our resource. As the field moves forward, it is imperative to integrate the insights obtained into efforts aiming at unravelling the genetic basis of diseases, ultimately contributing to the advancement of precision medicine around the globe.

## Acknowledgements

We thank all participants of the 1000 Genomes Project, without whom this project would not have been possible. We thank Julien Charest and Klaus Ehrlinger for assistance with long-read DNA sequencing, Wolfram Höps for providing valuable advice on the analysis of inversions, and Peter Ebert for assistance with data management. We acknowledge members of the HGSVC for valuable feedback provided on our approach and the generated findings. Moreover, we acknowledge the EMBL IT services, the Centre for Information and Media Technology at Heinrich Heine University Düsseldorf and the IT services at the Institute of Molecular Pathology (IMP) for providing resources for data processing and analysis. Finally, we thank the International Genome Sample Resource (IGRS) for assistance with providing the open data releases for this study. Funding for sequence data production was provided by the MARVL initiative, a collaboration between the IMP, BI X and Boehringer Ingelheim. Additional funding came from the following sources: National Institutes of Health (NIH) (to J.O.K., T.M., grant no. U24HG007497), Ministry of Culture and Science of the State of Northrhine Westphalia (to T.M, grant no. PROFILNRW-2020-107-A), and the GraphGenomes project funded by the BMBF (to T.M., grant no. 031L0184A, and J.O.K, 031L0184C). B.R.-M. was supported by a Bridging Excellence Fellowship provided by the Life Science Alliance.

## Data Availability

Our open data resource is fully available to the community for download at the International Genome Sample Resource (IGSR), at the following IGSR file transfer protocol (FTP) repository: https://ftp.1000genomes.ebi.ac.uk/vol1/ftp/data_collections/1KG_ONT_VIENNA/. This repository contains the graph and linear reference genomes, alignments, input SV callsets, the augmented genome graph, genotyped and phased structural variants as well as auxiliary data used for evaluating SV callset accuracy. For archiving purposes only, we will also mirror the complete dataset in the European Nucleotide Archive before the paper is accepted for publication.

## Code Availability

The pipeline producing a merged, multi-sample VCF from the post-processed SVarp calls (Methods, Section “SVarp VCF postprocessing”) is available at: https://github.com/eblerjana/long-read-1kg/tree/main/prepare-SVarp-callset (commit: <u>44a1752</u>). Our graph augmentation pipeline (Methods, Section “Graph augmentation”) is available at: https://github.com/eblerjana/long-read-1kg/tree/main/graph-augmentation-pipeline (commit: <u>44a1752</u>). The genotyping pipeline using Giggles (Methods, Section “Graph-aware genotyping with Giggles”), the associated preprocessing pipelines (Methods, Section “Preparing phased VCF panel”), SV calling with SVarp (Methods, Section “Variant calling using pangenome graph”), phasing pipeline (Methods, Section “Phasing with the ONT reads” and “Comparison of the WhatsHap phased VCFs against NYGC statistical phasing”), and the haplotype-tagging pipeline (Methods, Section “Haplotype-tagging of ONT reads”) are available at: https://github.com/marschall-lab/project-ont-1kg/tree/main/snakemake_pipelines (commit: <u>e42b63a</u>) Analysis script for genotyping are available at: https://github.com/marschall-lab/project-ont-1kg/tree/main/analysis_scripts/genotyping (commit: <u>e42b63a</u>) Further analysis scripts related to the base-calling, alignment to linear and graph-based references and SV calling are available at: https://github.com/1kg-ont-vienna/sv-analysis The structural variation annotation pipeline (Methods, Section “SVAN”) is available at: https://github.com/REPBIO/SVAN The inversion analysis scripts are available at at: https://github.com/celiatsapalou/Simulations_ONT_Data – for the simulations used, and https://github.com/celiatsapalou/Small_Inversions_Remap – for the pipeline for selecting regions for remapping and inversion calling.

## Methods

### Samples and DNA extraction

We initially selected 1,064 samples from the 1kGP collection, sourcing genomic DNA from Coriell and discarding low-quality DNA. The study complied with all relevant regulations for work with human subjects. DNA was extracted from lymphoblastoid cell lines (LCLs) and resuspended in TE buffer (10 mM Tris, pH 8.0, 1 mM EDTA).

### Quality Control Prior to Sequencing

DNA concentrations were determined with the Quant-iT™ dsDNA Broad-Range assay Kit (Q33130), using Varioskan™ LUX multimode microplate reader (VL0000D0). The purity evaluation of genomic DNA was performed on a DeNovix DS-11 Series Spectrophotometer. OD 260/280 and 260/230 ratio of 1.8 and 2 was maintained, respectively, for sequencing. Fragment length was measured on Femto Pulse system (Agilent, CA, United States, Cat N° M5330AA) using the Genomic DNA 165 kb Ladder Fast Separation assay with a separation time of 70 min (Agilent, CA, United States, Cat N° FP-1002-0275).

### Size Selection

All DNA samples were size selected using the Circulomics Short Read Eliminator Kit (Circulomics, MD, United States, SS-100-101-01). According to supplier information, the kit uses size-selective precipitation to reduce the amount of DNA fragments below 25 kbp in length. The kit was used according to the manufacturer’s recommendations (handbook v2.0, 07/2019). Briefly, 60 μl of Buffer SRE were added to the sample tube (60 μl volume), gently mixed and the tube centrifuged at 10,000 × g for 30 min at RT. After supernatant removal, two washing steps were performed with 200 μl of 70% EtOH and a centrifugation at 10000 × g for 2 min at RT. Finally, 50 μl Buffer EB was added and the tube was incubated at 50 °C for 30min, followed by overnight incubation at RT to ensure efficient DNA elution.

### ONT library preparation and sequencing

Sequencing library preparation was carried out following the general guidelines from Oxford Nanopore Technologies, with modifications proposed by New England Biolabs (NEB) to ensure high yield data generation and long-fragment sequencing. For library preparation, the following reagents were used: Ligation Sequencing Kit SQK-LSK110 (Oxford Nanopore Technologies), NEBNext Companion Module for Oxford Nanopore Technologies Ligation Sequencing (NEB, MA, United States, Cat N° E7180S), and AMPure XP beads (made in-house by the Molecular Biology Service, IMP). A DNA amount of 3 μg as input material was transferred into a 0.2 ml thin-walled PCR tube and the total volume was adjusted to 48ul with Nuclease free water (ThermoFisher, cat # AM9937). DNA fragments were repaired and end-prepped as follows: 3.5 μl NEBNext FFPE DNA Repair Buffer, 2 μl NEBNext FFPE DNA Repair Mix, 3.5 μl NEBNext Ultra™ II End Prep Reaction Buffer, and 3 μl NEBNext Ultra™ II End Prep Enzyme Mix were added to each tube. After mixing and spinning down, the samples were incubated at 20°C for 30 min, followed by a second incubation at 65°C for 5 min. The prolonged incubation time allowed recovery of longer fragments. The solution from each tube was then transferred to a clean 1.5 ml Eppendorf DNA LoBind tube (Eppendorf AG, Hamburg, Germany) for clean-up. First, 60 μl of AMPure XP Beads were added to each tube. The samples were then incubated on a HulaMixer™ sample mixer (Thermo Fisher Scientific, MA, United States, 15920D) for 5 min at RT. Bead clean-up was performed with two washing steps on a magnetic rack, each time pipetting off the supernatant and adding 200 μl of freshly prepared 70% EtOH. The pellet was resuspended in 61 μl nuclease-free water and incubated for 5 min at RT. Tubes were placed on a magnetic rack to collect the final eluate (1 μl was then taken out for quantification). For adapter ligation and clean-up, 60 μl DNA from the previous step was combined with 25 μl Ligation Buffer LNB, 10 μl NEBNext Quick T4 DNA Ligase and 5 μl Adapter Mix AMX in a 1.5 ml Eppendorf DNA LoBind tube. The reaction was then incubated for 20 min at RT. A second AMPure bead clean-up step was carried out by adding 40 μl of bead solution to each tube, followed by incubation on a HulaMixer™ for 5 min at RT. After pipetting off the supernatant on a magnet rack, the beads were washed twice with 250 μl Long Fragment Buffer LFB. Finally, the supernatant was discarded, the pellet resuspended in 25 μl Elution Buffer EB, and incubated for 10 min at 37℃ to collect the final library. Samples were quantified using a Qubit fluorimeter and diluted appropriately before loading onto the flow cells. The final mass loaded on the flow cells was determined based on the molarity, dependent on average fragment size. Sequencing was carried out using FLO-PRO002 (R9.4.1) flow cells from ONT on the PromethION 48. The sequencing run was stopped after 24h, flow cell washed using the Flow cell wash kit XL (EXP_WSH004-XL, Oxford Nanopore Technologies) and then the library was reloaded.

### Base calling and adapter trimming

Guppy 6.2.1 was used for base calling the Fast5 input files in “sup” accuracy mode with adapter trimming and read splitting disabled and the output was converted to FASTA. For adapter trimming and read splitting, we subsequently used Porechop^95^ version 0.2.4 on the generated FASTA files in chunks of 300,000 reads with default parameters.

### Reference genome alignments

Reads were aligned to the GRCh38 and CHM13 linear reference genomes as well as the prebuilt human genome graph (https://doi.org/10.5281/zenodo.6983934) constructed from HPRC year-1 samples^20^. For the GRCh38 and CHM13, we used minimap2^96^ to map the ONT reads using the options “-a -x map-ont --rmq=yes --MD --cs -L”. Samtools^97^ was used to sort the alignments and convert to CRAM. Multiple ONT runs for the same sample were tagged using different read-groups. Minigraph^41^ version 0.20-r559 was used to map the ONT reads against the graph genome using the options “--vc -cx lr”. The “--vc” flag enables the output of the alignments in vertex coordinates and the “lr” option enables long-read mapping. The resulting graph alignments in GAF format^41^ were sorted using gaftools^98^ and compressed using bgzip.

### Sample and alignment quality control

Using SNP calls of the high-coverage short-read data from the 1kGP cohort^3^, we investigated all samples for cross-contamination and potential sample swaps. We first genotyped all SNPs from the short-read haplotype reference panel with an allele count (AC) greater or equal to 6 in all samples using bcftools^99^. Individual VCF files were merged using bcftools^99^ into a multi-sample VCF file that was then combined with the short-read haplotype reference panel. We then used VCFtools^100^ to calculate a relatedness statistic of the long-read sequenced samples compared to the short-read sequenced samples using the “--relatedness2” option. This analysis identified a sample swap between HG01951 and HG01983, which we relabelled afterwards. We excluded all samples that appear to be cross-contaminated during library preparation, namely HG00138, HG02807, HG02813, HG02870, HG02888, HG02890, HG03804 and HG03778.

Alignments were analysed using NanoPack^101^ to determine median and N50 read length and genome coverage (**Fig. S2, Fig. S4, Fig. S43**). To compare linear and graph genome alignments in terms of percent identity, number of aligned reads (bases) and largest Cigar I and D event, we used lorax^26^ (**Fig. S3**).

### Structural variant calling using linear reference genomes

We used Sniffles^22^ version 2.0.7 and a new long-read version of Delly^39^ version 1.1.7 to discover SVs using a linear reference genome. For Sniffles, we converted CRAM files to BAM format using samtools and then calculated for each sample candidate SVs and the associated SNF file using Sniffles. We then employed Sniffles population-calling mode on all SNF files to generate two multi-sample VCF files, one for GRCh38 and one for CHM13.

Similarly, we also used Delly’s population-calling mode which first calls SVs by sample using the new long-read (lr) subcommand. We then merged all candidate SV sites using Delly merge with the options “-p -a 0.05 -v 3 -c” to select PASS sites that are precise at single-nucleotide resolution with a minimum variant allele frequency of 5% and a minimum coverage of 3x. We then genotyped this SV site list in all samples using delly lr and merged the results by id with bcftools merge using the option “-m id”. We then applied sansa (https://github.com/dellytools/sansa), a newly developed, multi-sample SV annotation method that detects SV duplicates based on SV allele and genotype concordance, to remove redundant SV sites. The parameters for the sansa markdup subcommand were “-y 0 -b 500 -s 0.5 -d 0.3 -c 0.1” to mark SV duplicates for sites that show an SV size ratio > 0.5, a max. SV allele divergence of 30%, a maximum SV breakpoint offset of 500 bp and a minimum fraction of shared SV carriers of 10%. After removing duplicates, we generated the final multi-sample VCFs for GRCh38 and CHM13.

To ensure specificity and facilitate genome graph augmentation, we further generated for Sniffles and Delly separately a consensus SV callset of SVs shared between the GRCh38 and CHM13 reference genome. We therefore lifted the GRCh38 call sets to CHM13 using bedtools^102^ and UCSC’s liftOver tool^103^ with the GRCh38 to CHM13 chain file. We then compared the lifted VCF with the original CHM13 VCF file to identify shared SVs using sansa’s compvcf subcommand with the options “-m 0 - b 50 -s 0.8 -d 0.1” to identify SVs that have a size ratio >= 0.8, a max. SV breakpoint offset of 50 bp and an SV allele divergence of at most 10%. Since the SV allele divergence filter requires single-nucleotide precision (e.g. Delly’s consensus SV allele sequence), we did not apply this filter to Sniffles. For genome graph augmentation, we subset the final VCFs to deletions and insertions only. All inversion-type SVs called by Delly and Sniffles using the minimap2 alignments were integrated separately in the inversion analysis.

### Inversion analysis

We developed a multi-tiered analytical pipeline to comprehensively ascertain inversions using long-read sequencing data. By inspecting previously known inversions in our dataset, along with simulating a range of small inversions (<1 kbp) with a coverage mirroring our actual data (median 17X), we discovered that minimap2 frequently misaligned reads in small inversion regions, leading to increasing error rates in those genomic locations. Conversely, NGMLR demonstrated superior performance in accurately aligning reads within these challenging areas. (**Fig. S34).**

Therefore, using pysamstats (https://github.com/alimanfoo/pysamstats), we first calculated the mismatch rate per base pair in 50 bp intervals, initially in simulated datasets to tune parameters and subsequently in the real data to identify candidate inverted regions for realignment. This analytical process revealed that the majority of genomic regions maintained a mismatch rate below 20%; regions surpassing this rate were identified as having a significantly high mismatch rate and selected for realignment with NGMLR^22^ (**Fig. S44**). Post-exclusion of telomeric and centromeric regions, as well as of misaligned regions exceeding 1kb in size, was then conducted to restrict the number of regions requiring remapping, thereby enhancing computational efficiency. The selected genomic segments underwent realignment using NGMLR and we then interrogated all realigned regions with Delly^39^ to discover previously missed inversions. The final inversion calls in the remapped regions from all samples were consolidated using SURVIVOR merge^65^. For the rest of the regions not requiring realignment, inversion calling was conducted using both Sniffles and Delly. Manual verification of true versus false positive calls was performed by examining dot plots and IGV-like plots generated with wally^26^ for each candidate inversion location for the largest ten reads per candidate region, ensuring the accuracy of our findings. Ultimately, we generated a final comprehensive inversion callset by merging all unique instances from each dataset with ‘bedtools (v.2.31.1) merge^92^’. As the inversion analysis was conducted on the GRCh38 reference genome, regions were subsequently lifted over to the T2T-CHM13+Y genome, applying a 90% base remapping threshold to retain a region.

To evaluate our inversion dataset against two prior studies on 1kGP samples^11,55^, we utilized ‘bedtools intersect’ (v.2.31.1)^92^, defining inversions as ‘known’ if they exhibited a minimum of 50% reciprocal overlap with inversions from either prior dataset. Analysis against the HGSVC dataset^11^, which delineates 188 inversions through whole-genome assembly, underscored the efficacy of our methodology in detecting a diverse range of inversions, both repeat-mediated and non-repeat-mediated: our results showed that 65% of non-repeat-associated inversions and 41% of SD-mediated inversions were successfully identified. Furthermore, we refined our comparison to a recent 1kGP-derived short-read inversion dataset^55^, where we included only those inversions from the comparison dataset with quality scores of 30 or higher, to ensure the accuracy of our comparative analysis. This approach revealed an overlap of 289 inversions, or 36.5% concordance (median size of 530 bp). This significant concordance attests to the robustness of our analysis. In conclusion, our study enhances the current understanding of genomic inversions by identifying 401 previously uncharacterized inversion events within the 1kGP cohort.

Regarding the flanking repeats in repeat-rich inverted regions, we conducted a detailed analysis by manually inspecting the repeat types and their orientations at inversion breakpoints. Repeat data were acquired from the RepeatMasker track and the Segmental Duplication annotations of the CHM13 reference^5,20,38^ (obtained from https://github.com/marbl/CHM13); an inversion was classified as repeat-mediated if it was bracketed by repeats in reverse orientation relative to each other, detected through dotplot analysis.

### Phasing with the ONT reads

We conducted phasing experiments with the ONT reads to check how well they compare to the statistical phasing done previously in high-coverage 1kGP short read data generated by the New York Genome Center (NYGC)^3^. The NYGC phased VCFs and the NYGC raw genotypes were used. Using the NYGC raw genotypes, the phasing was done by WhatsHap^104^ (version 2.0) in three different ways: phasing with only the ONT reads (from hereon referred to as longread phasing), trio phasing, and trio phasing with the ONT reads (from hereon referred to as longread-trio phasing). The trio phasing and longread-trio phasing was done for the 6 complete family trios (Family IDs: 2418, CLM16, SH006, Y077, 1463 (paternal side), 1463 (maternal side)) for which our resourcce has long read data. The longread phasing was done for all the 967 samples in the intersection of our 1019 sample set and the NYGC sample set of 3202 (**Fig. S8, Table S20**). The phasing was done for only the autosomes and each chromosome was phased separately to allow parallel processing. The commands are as follows:

- Long-read phasing: whatshap phase -o <output phased vcf> -- chromosome <chromosome id> --sample <sample name> -r <reference fasta> <nygc raw genotypes for sample> <ont cram for sample>
- Trio phasing: whatshap phase -o <output phased vcf> --chromosome <chromosome id> -r <reference fasta> --ped <pedigree data for the NYGC samples> <NYGC raw genotypes for samples in a family>
- Long-read trio phasing: whatshap phase -o <output phased vcf> -- chromosome <chromosome ID> -r <reference fasta> --ped <pedigree data for the NYGC samples> <NYGC raw genotypes for samples in a family> <ont cram for samples in a family>

### Comparison of the WhatsHap phased VCFs against NYGC statistical phasing

The phased VCFs produced by WhatsHap were compared against the NYGC statistical phasing using WhatsHap’s compare function. The commands are as follows:

- For the samples without trio data: whatshap compare --sample <sample name> --names longread,nygc --tsv-pairwise <pairwise tsv file> -- tsv-multiway <multiway tsv name> <input longread phased vcf> <input nygc statistical phasing vcf>
- For the samples with trio data: compare --sample <sample name> --names trio,longread,trio-longread,nygc --tsv-pairwise <pairwise tsv file> --tsv-multiway <multiway tsv name> <input trio phased vcf> <input longread phased vcf> <input longread trio phased vcf> <input nygc statistical phasing vcf>

### Haplotype-tagging of ONT reads

The ONT reads were haplotype-tagged (or haplotagged) using WhatsHap^104^ (version 2.0). The New York Genome Center phased VCF^3^ was used as the reference for tagging the reads. The command used to tag the reads was: `whatshap haplotag --skip-missing-contigs --reference <reference fasta> --sample <sample name> --output-haplotag-list <output file> - -output /dev/null <NYGC phased VCF> <ont cram>`

Although the main output of `whatshap haplotag` is a tagged alignment file, downstream tools used in this study only required a file containing the tag for each read which is given in `--output- haplotag-list.` Due to the presence of pseudo-autosomal regions (PAR) in the phased VCF, the command was altered for the male samples. Instead of providing the entire NYGC Phased VCF, the non-PAR records were removed and the haplotagging was performed. Post-haplotagging, the list of reads aligning to the non-PAR on chrX were extracted, assigned manually as the maternal haplotype, and added to the haplotag list.

### Variant calling using pangenome graph

The aim of SVarp^40^ is to discover SVs on graph genomes. These are the variants that are missing in a linear reference, but are on top of alternative sequences present in the pangenome. With SVarp, we call novel phased variant sequences, called svtigs, rather than a variant call set, which we later use in the graph augmentation step. In order to discover phased structural variation assemblies (svtigs) on top of the pangenome graph, we utilised haplotag read information and the alignment file (i.e. GAF alignments) as input to SVarp algorithm using <`svarp -a GAF-FILE -s 5 -d 500 -g GFA-FILE --fasta READS-FASTA-FILE -i SAMPLE_NAME --phase HAPLOTAG-FILE>` command. With 967 samples, we found a total of 1,108,850 variants (∼1145 per sample).

In order to find specific SV breakpoint loci with respect to a linear reference genome, we used the PAV tool^5^ to call SV breakpoints, using svtigs that SVarp generated, as input. This yielded 1,258,880 and 1,241,252 SVs relative to the CHM13 and GRCh38 linear genomes respectively, that are >50bps. However, we realised that some svtigs give rise to multiple SVs in the output of PAV. To ensure that variants called from the same svtig end up on the same pseudo-haplotypes in the graph augmentation step (see Section “Graph augmentation”), we generated a script to combine records arising from the same svtig into a single VCF record. This is done by connecting multiple smaller such SVs into a single SV record by adding reference sequence in between. This yielded 564,661 and 562,311 SVs relative to CHM13 and GRCh38 respectively.

### SVarp VCF postprocessing

The single-sample VCFs (relative to GRCh38) generated with PAV from the SVarp svtigs were merged into a multi-sample VCF using `bcftools merge`^99^ (version 1.18) and then post processed using `truvari collapse`^105^ (version 4.1.0). The latter step merges SV records likely representing the same event into a single record removing redundancy. This reduced the number of SVs from 451,942 to 215,209. Finally, we filtered the resulting VCF by keeping only records present in at least two samples. This filtered set contained 70,932 SV.

### Graph augmentation

We developed a pipeline to add additional variants found by Delly, Sniffles and SVarp across the 1kGP ONT samples to the minigraph graph so that they can be genotyped by giggles. The main idea is to construct so-called “pseudo-haplotypes” by implanting sets of non-overlapping variant calls into a reference genome and then adding them to the graph using minigraph^41^ (**Fig. S7**). Our pipeline consists of the following steps. At first, we remove variant calls that fall into the centromere regions and mask the respective region in the reference sequence by Ns using the tool `bcftools maskfasta` (version 1.18). We used the GRCh38 reference genome and centromere annotations obtained from UCSC genome browser. In the next step, we generate the pseudo-haplotypes as follows. Each of the callsets (Delly, Sniffles, SVarp) contains variants overlapping across samples. Thus, inserting all of them into one reference genome will fail.

Therefore, we first group variants of each callset into sets of non-overlapping variants, and then generate a consensus sequence for each of these sets by implanting the variants into the reference genome using `bcftools consensus` (version 1.18). As a result, we obtain a whole-genome consensus sequence for each of these sets, which we call the pseudo-haplotypes. For the Delly calls, we obtained 26 such pseudo-haplotypes, for Sniffles we got 69, and for SVarp we generated 117 pseudo-haplotypes. In total, these pseudo-haplotypes carry 154,319 Delly SVs, 128,688 Sniffles SVs and 70,813 SVs detected by SVarp. In the last step, we insert all of these newly constructed genome sequences into the HPRC minigraph using the tool minigraph (version v0.20). We first inserted all SVarp haplotypes, then all Delly haplotypes and finally all Sniffles haplotypes into the graph using the command: `minigraph -cxggs -t32 <minigraph> <svarp pseudo-haplotype fastas> <delly Pseudo-Haplotype fastas> <sniffles pseudo-haplotype fastas> > augmented-graph.gfa`

In the augmented graph, the number of bubbles (**Fig. S9**) increases to 220,168 (102,371 in the original graph) and the total sequence represented in the graph increased from 3,297,884,175 bases to 3,477,266,061 bases during graph augmentation. In order to identify bubbles in the augmented graph representing variation that was previously not represented in the original graph, we created BED files with coordinates of bubbles present in both graphs using `gfatools bubble`. Then we used `bedtools closest` to compute the distance between each bubble in the augmented graph and their respective closest bubble in the original graph (**Fig. S11**). We defined all bubbles in the augmented graph whose distance to the closest bubble in the original graph is at least 1kb as “new” bubbles, representing novel SV sites.

In order to evaluate our augmented graph, we aligned ONT reads of human sample HG00513 to it using the command: `minigraph -cx lr -t24 augmented-graph.gfa HG00513- reads.fa --vc 2 > alignments.gaf`. For comparison, we aligned the same set of reads to the original HPRC minigraph graph using the same command. We then computed alignment statistics using `gaftools stat` (**Table S3**). We observed better alignment statistics when aligning reads to the augmented instead of the original graph. The number of aligned reads increased by 33,208, and the number of aligned bases by 152,454,715 base pairs.

### Preparing phased VCF panel

We developed a pipeline that can reconstruct the alleles of the samples in the graph using the graph and the assemblies for the samples. Firstly, the bubbles in the graphs are identified using gaftools^98^ (commit ID: 919bbecf51602161db7ba6e859cc5c26a7413c83). The function order_gfa tags the nodes of the graph to identify whether the nodes are bubble nodes (nodes inside a bubble) or scaffold nodes (nodes outside a bubble). The phased panel is created by aligning the HPRC assemblies back to the tagged HPRC graph using the tool minigraph^41^ (version: 0.20-r559). The resulting alignments are processed to identify the alleles using the node paths between scaffold nodes. The allele information from the haplotype assemblies is then converted to a phased panel VCF. For the augmented graph created with the addition of pseudo-haplotypes into the HPRC graph, the same pipeline discussed above works where the pseudo-haplotypes are considered as assemblies and aligned to the augmented graph. Based on the tagging of the augmented graph, the alleles on the pseudo-haplotypes are identified and separate columns are created in the phased panel VCF corresponding to the alleles of the pseudo-haplotypes. This pipeline creates a multisample phased VCF, containing multiple alternate alleles for each record of the VCF.

The phased panel VCFs are processed for its inclusion in the Giggles genotyping pipeline. Internal filter tags are set during the VCF creation step which allows for filtering the bubbles where allele information is not present for at least 80% of the samples. Additionally, bubbles are filtered out which don’t have any alternate alleles in any of the haplotypes. From the phased panel VCF, a “Giggles- ready” version is created where the reference alleles in the pseudo-haplotypes are masked with dots, since internally Giggles accounts for the allele frequency in the panels and the pseudo-haplotypes heavily bias the genotyping towards the reference used in the `bcftools consensus` step. The VCF records describe bubble structures in the graph. These bubbles often contain many nested and overlapping variant alleles, i.e. a bubble does not necessarily represent a single variation event. In order to identify variant alleles represented in these bubbles, we applied the same bubble decomposition approach as described by the HPRC^20^. In short, the idea is to compare node traversals of the reference and alternative paths through bubbles in order to identify nested alleles. As in the HPRC’s study^20^, we then annotate the bubbles in our VCF to encode nested variants, so that we can translate genotypes computed for bubbles to genotypes for the variants represented in the graph.

### Graph-aware genotyping with Giggles

Genotyping was done using Giggles (version: 1.0). Giggles is a pangenome-based genome inference tool which leverages long-read DNA sequences. It serves as a long-read alternative to PanGenie’s short-read based approach^106^. It works with a Hidden Markov Model (HMM) framework where each variant position contains states corresponding to all possible genotypes at that position. The transition probabilities are based on the Li-Stephens model^107^ and the emission probabilities are based on the alignment of reads around an interval window of the variant position. Forward-Backward computation of the HMM gives the posterior likelihoods of each state at each variant position which can be used to compute the likelihood of genotypes across the genome.

Giggles requires a phased VCF panel, sorted graph alignments, the input long reads, tagged graph, and the haplotype tagging of the reads. We use the command: `giggles genotype --read- fasta <input reads> --sample <sample name> -o <output vcf> --rgfa <tagged gfa> --haplotag-tsv <haplotype tags> <phased panel VCF> <sorted alignments>`. This outputs an unphased VCF with the genotypes for the given sample.

The VCFs were further filtered using bcftools^99^ to create a high-quality genotype set. This was done by masking genotypes of samples having genotyping quality of less than 10 and dropping variant positions where more than 5% of genotypes are missing. The commands are: bcftools view --min-ac 1 -m2 -M2 <vcf> | bcftools +setGT - -- -t q - n. -i ‘FMT/GQ<10’ | bcftools +fill-tags - -- -t all | bcftools filter -O z -o <filtered-vcf> -i ‘INFO/AC >= 1 && INFO/F_MISSING <= 0.05’

### Mendelian inconsistency statistics

For the 6 trios in the 967 samples of our dataset we calculated Mendelian inconsistency statistics (**Fig. S45**). We provide the confusion matrices and various statistics for each family, genotyped against HPRC_mg and HPRC_mg_44+966 and also report for various variant types depending on whether the variant came from a biallelic or multiallelic bubble (**Table S5-S12**). The definitions for each statistic can be found in **Table S4**.

### Statistical phasing using prior short-read data of 1kG samples

Using SHAPEIT5^44^, we statistically phased the multi-sample VCF file outputted by Giggles using a recently constructed CHM13 haplotype reference panel^45^. This panel uses SNP and InDel calls generated from 1kGP short-read data^3,43^ for CHM13. We first subsetted the unphased input VCF from Giggles with 167,291 SVs to the 908 unrelated samples present in the panel and then used SHAPEIT5’s common variant phasing mode to incorporate the new SV alleles into the SNP and InDel haplotype scaffold, yielding a phased VCF with 164,571 SVs. Prior to the phasing we splitted multi-allelic variants into bi-allelic variants using bcftools norm^99^ with the option “-m -any”. After phasing, we joined multi-allelic variants back into the original state using bcftools norm with the option “-m +any”.

Using plink^108^, we assessed linkage disequilibrium (LD) to nearby SNPs in a window of 1 Mbp. We first converted the VCF to plink input files and then ran plink to calculate r2 values of all SVs to nearby SNPs. For the LD analysis, we further subdivided the CHM13 genome into “difficult” regions and “high-confidence” regions using BED files provided by the Genome in a Bottle project, namely the CHM13_notinalldifficultregions.bed.gz (**Fig. S25, Fig. S26**).

### Compacted de Bruijn graphs of short-read data of 1kG samples

To facilitate efficient estimation of the average quality value (QV) of reconstructed haplotypes and the accuracy of novel insertions, we used the previously generated deep-coverage short-read 1kGP data^3^ to build compacted de Bruijn graphs from error-corrected reads using lighter^109^ and bcalm2^110^.

We first applied lighter^109^ to the short-read FASTQ files using a k-mer length of 23 and the options “- trim -discard” to allow trimming of reads and discarding reads that cannot be corrected. All successfully corrected reads were provided as input to bcalm^110^ to construct a compacted de Bruijn graph from the sequencing data using a k-mer length of 61 and a required minimum k-mer abundance of 3.

### Haplotype QV estimation and insertion accuracy estimation

To estimate QV values for reconstructed haplotypes, we first implanted for each sample all phased variants (SNPs, InDels and SVs) into the CHM13 reference using bcftools consensus^99^ for haplotype 1 (option “-H 1”), called sample.h1.fa, and haplotype 2 (option “-H 2”), called sample.h2.fa. We then used yak^47^ count on the matched compacted de Bruijn graph to build a short-read based k-mer hash table, called sample.dbg.yak, using “-K 1.5g -k 31”. With the reconstructed haplotype and the k-mer hash table, we then calculated a haplotype QV estimate using yak qv with the options “-p -K 1.5g -l 100k sample.dbg.yak sample.h[1|2].fa” for haplotype 1 and 2, respectively (**Fig. S19**).

To estimate the insertion accuracy, we augmented the CHM13 reference with all new insertions and then mapped all short-read derived compacted de Bruijn graph sequences against the extended reference using bwa mem^111^. For a singleton SV, occurring in only 1 sample at allele count 1, the expected coverage is 1x and we therefore computed for all insertions the average depth using samtools depth^99^ with the option “-aa”. For FDR estimation, an insertion with average coverage less than 1x was counted as a false positive (**Fig. S15**).

### Copy-number accuracy estimation using intensity rank sum (IRS) testing

The IRS test was previously used to estimate an FDR for copy-number unbalanced SVs^4^. The IRS test leverages differences in microarray probe-level intensities between samples expected to have different copy number states. From our final SV call set with 908 samples, microarray probe-level intensities were available for 901 samples. To use the IRS test, we first lifted all array probes to the CHM13 assembly using UCSC’s liftOver tool^103^ with the GRCh38 to CHM13 chain file. We then extracted all deletions and duplications and applied the IRS test from the SVAnnotator tool of Genome STRiP^46^ with the input VCF files and the previously used array intensity file^4^ lifted to CHM13. We further subdivided the CHM13 genome into “difficult” regions and “high-confident” regions using BED files provided by the Genome in a Bottle project, namely the CHM13_notinalldifficultregions.bed.gz (**Fig. S14**).

### Comparison to previous SV callsets

Since most previous callsets used GRCh38 as reference genome, we first lifted our SV calls to GRCh38 using UCSC’s liftOver tool^103^ with the CHM13 to GRCh38 chain file. As baseline callsets, we included the callset produced by Byrska-Bishop et al.^3^, Ebert et al.^5^ and Sudmant et al.^4^, labelled as nygc, pangenie and phase3, respectively **(Fig. S18)**. The comparison of SVs was restricted to autosomes and chrX for deletions and insertions separately. We used sansa compvcf to compare the SV VCF files using default parameters. We altered the base and comparison VCF to identify SVs that are distinct for one or the other callset and to identify potential 1:many SV overlaps among the shared SVs. We used the presence of INSSEQ and the absence of IMPRECISE in the INFO field as criteria for sequenced-resolved insertions in prior short-read SV callsets^3^ (**Fig. 1**).

### Population differentiation

We used ADMIXTURE (1.3.0) to compute admixture for K=[3..10]. For performing SV based PCA we used EIGENSOFT (8.0.0). Fst values were calculated using VCFtools for each continental population (against the remaining samples) and each population (against the remaining samples) per site and for 1 Mbp windows.

### SV impact estimation on genomic features

We used vep with annotation from the CHM13 rapid release of Ensembl (107) to estimate the impact of the SVs on genomic features. Processing was performed using the command: `vep --assembly T2T-CHM13v2.0 --regulatory --offline -i final- vcf.unphased.vcf`

We observe a large difference in the mean AF of SVs affecting coding (mean MAF 0.009) and non-coding (mean MAF 0.061) regions. This observation and the p-value of 0.0 reported by both t-test and KS-test thereby supporting the hypothesis that the two underlying distributions are different.

### Targeted genotyping of challenging loci

Locityper^87^ (version: 0.13.3) is a targeted WGS-based genotyper designed for challenging loci. Our initial set of target genes was preprocessed using `locityper add -e 300k`. ONT WGS datasets were preprocessed and genotyped with `locityper preprocess --tech ONT` and `locityper genotype`, respectively. A database of loci haplotypes was constructed based on the GRCh38 and CHM13 reference genomes, as well as based on the 44 diploid HPRC whole genome assemblies. In order to evaluate genotyping accuracy, we aligned haplotypes to each other using Wavefront alignment algorithm^112^, and calculated sequence divergence as a ratio between edit distance and alignment size. Local haplotypes were extracted from the NYGC call set using `bcftools consensus`.

### Annotation of SV classes with SVAN

Deletion and insertion SV calls were processed with the SV ANnotation (SVAN) tool to classify them in distinct classes based on distinctive sequence features:

#### Retrotranspositions

Candidate retrotransposition events were identified by searching for poly(A) and poly(T) tails at the 5’ end 3’ ends of the deleted and inserted sequences. Poly(A/T) tracts were required to be at least 10 bp in size, have a minimum purity of 90% and be at a maximum of 50 bp to the insert end or begin, for poly(A) and poly(T), respectively. The sequence corresponding to the target site duplication was detected and trimmed from either the 5’ or 3’ end of the L1 insert through the identification of exact matches with the genomic sequence at the integration position. In order to have all candidate retrotransposed inserts in forward orientation, the reverse complement sequence for every trimmed insert occurring in the minus strand was obtained.

Candidate 3’ partnered transductions were detected by searching for a second poly(A) stretch at the 3’ end of the trimmed sequences using the same criteria outlined above. Integrants with a secondary tail were annotated as ‘3’ transduction’ candidates with the sequence in between the poly(A) stretches corresponding to the transduced bit of DNA, while those with a single tail were classified as ‘solo’ candidates. In order to trace transductions to their source loci, transduced sequences were aligned onto CHM13 using BWA-MEM^111^. In order to maximise sensitivity for particularly short transduction events a minimum seed length (-k) of 8 bp and a minimum score (-T) of 0 were used. Alignment hits were filtered by requiring a minimum mapping quality of 10. Transduction candidates with at less than 80% of the transduced aligning on the reference were further filtered out.

The retrotransposition candidates were further trimmed by removing the poly(A) tails and transduced sequences. In order to identify processed pseudogenes, orphan transductions, 5’ partnered transductions and retroelement sequences, the resulting trimmed inserts were aligned onto CHM13 using minimap 2.1, as well as BWA-MEM 0.7.17-r1188, by employing the parameters described above and with the alignment conducted onto a database of consensus L1, Alu and SVA repeats.

Alignment hits were filtered by requiring a minimum mapping quality of 10, and chained based on complementarity in order to identify the minimum set of nonoverlapping alignments that span the maximum percentage of the trimmed insert. Based on the alignment chains, retrotranspositions are classified as processed-pseudogenes (≥75% aligns on the reference over a single or multiple Gencode 35 annotated exons^113^), orphan transductions (≥75% aligns on the reference outside exons), 5’ transductions (≥75% aligns both into the reference on its 5’ and into the consensus of a retrotransposon) and solo (≥75% aligns into a single consensus retrotransposable element).

#### Non-canonical MEI

Insertions and deletions without a poly(A) tail detected were aligned into a database of consensus L1, Alu and SVA repeats as described above to identify non-canonical MEIa. Alignment hits were filtered and chained and non-canonical MEI calls made if ≥75% of the sequence aligned into one or multiple repeat classes.

#### Endogenous retroviruses

Inserted and deleted sequences were aligned with BWA-MEM 0.7.17-r1188 into a database containing consensus ERV and LTR sequences. Alignment hits were filtered and chained as described above. SVs with hits spanning ≥75% of their sequence on the retrovirus database were classified as Solo-LTR and ERVK, based on the presence of an LTR alone or LTR plus retroviral sequence, respectively.

Expansions and contractions of VNTR loci were annotated by processing inserted and deleted sequences with TRF v4.04.^114^ TRF hits were processed and chained based on complementarity in order to determine the minimum set of non-overlapping hits that span the maximum fraction of the sequence. Based on the hit chains, VNTRs are classified as simple (≥75% of the sequence corresponds to a single repeated motif) and complex (≥75% corresponds to more than one repeated motif).

#### Duplications

Diverse classes of duplications, including tandem, inverted and complex duplication events, were annotated by realigning the inserted sequences onto the reference using minimap 2.1. Alignments were filtered to select only those located within a 2kb window around the insertion breakpoint. These were further chained based on complementarity to determine the minimum set of non-overlapping hits spanning the maximum percentage of the sequence. Based on the alignment chains, duplications are classified as tandem (≥75% of the sequence aligns at the insertion breakpoint in forward orientation), inverted (≥75% of the sequence aligns at the insertion breakpoint in reverse orientation) and complex (≥75% of the sequence aligns at the insertion breakpoint both in forward and reverse orientation). *NUMTs*. Insertions with ≥75% of its sequence having one or multiple minimap 2.1 alignments on the mitochondrial reference are classified as Nuclear Mitochondrial DNA (NUMT).

### Analysis of MEI transductions

L1 and SVA transductions were clustered based on the alignment position on the reference of their corresponding transduced sequences to identify their source regions. A buffer of 10 kbp was applied to their start and end alignment position prior to clustering. Source regions were further intersected with a database of full-length loci to determine the progenitor repeat. This database was generated by aggregating all the full-length L1s (>5.9 kb in size) and SVAs (>1 kb) in CHM13 and detected as non-reference insertions in our SV callset. Progenitor repeats were further annotated by determining their insertion orientation and subfamily. For L1 events, subfamily assignment was performed through the identification of subfamily diagnostic nucleotide positions on their 3ʹ end. L1 progenitors bearing the diagnostic “ACG” or “ACA” triplet at 5,929-5,931 position were classified as “pre-Ta” and “Ta”, respectively. Ta elements were subclassified into “Ta-0” or “Ta- 1” according to diagnostic bases at 5,535 and 5,538 positions (Ta-0: G and C; Ta-1: T and G).

Elements that did not display any of these diagnostic profiles could not be assigned to a particular category, and their subfamily status remained undetermined. For SVAs, the inserts were processed with RepeatMasker v4.0.7 to determine the subfamily. If multiple RepeatMasker hits were obtained, the one with the highest Smith-Waterman score was selected as representative. To assess a bias in source elements to generate 5′ or 3′ transduction events, respectively, we conducted two-tailed binomial tests followed by controlling for the FDR according to Benjamini and Hochberg. For SVAs, in which 5′ and 3′ transductions occur at approximately similar abundance across the dataset, we formally tested all source elements with at least 10 transductions for a statistical 5′ vs. 3′ bias. In the case of L1s, in which 5′ transductions are much rarer than 3′ transductions, we included all source elements with at least 5 transductions when conducting statistical testing for 5′ vs. 3′ bias.

### SV breakpoint junction analysis

The detection of homologous sequences and microhomologies flanking deletions and insertions was conducted on the primary callset, after removing calls less than 50 bp in length, using two approaches: Microhomology was quantified using the available homology output from SV calls generated by Delly^39^ (v.1.2.6). We systematically generated Delly calls for each SV by building synthetic reads carrying therespective SV allele, mapping those synthetic reads with minimap2^96^ (v.2.2.26) onto the genomic reference of the SV with 4 kb padding and calling the respective SV with Delly. The synthetic reads consist of 2kb reference sequence upstream of the SV start, the inserted sequence (only for insertions and modified for larger insertions) and 2kb reference sequence downstream of the SV end coordinate. For insertions longer than 100 bp we used at most the first and last 50 bp of the inserted sequence to avoid aberrant mapping of the insert, which was especially observed in the case of long insertions. Due to the use of truncated inserts for insertions only homologies with a length of 50 bp were considered, and larger values were set to 50 bp. The second approach aimed to capture longer stretches of homology using BLAST^115^ based detection of homologous stretches of DNA. For this various pairs of search windows around both breakpoints were defined. Search windows were either symmetric with a padding of 50 bp, 100 bp, 200 bp, 400 bp, 1 kb, 2 kb, 5 kb, 10 kb, 50 kb and 100 kb or asymmetric with a window size of “SV length”, which is shifted regularly along the breakpoint (0 bp upstream / “SV length” bp downstream; 1/6 “SV length” upstream / 5/6 “SV length” downstream; …; “SV length” upstream / 0 bp downstream). From the predefined windows only those leading to no overlap between windows were used. Potential homologies were detected using blastn 2.12.0 with -perc_identity 80 and -word_size 5, where the sequences inside the search windows were passed using the -subject and -query parameters. In cases of insertions the inserted sequence was inserted into the reference genome and the search windows were defined on the modified reference sequence. BLAST results were filtered to ensure flanking homology stretches show the same directionality, span the respective SV breakpoints, and contain the respective breakpoint at the relatively same position inside the homology segment.

For annotation of repeat-mediated SV the overlap of the homologous sequences with RepeatMasker (v.4.1.2) annotations was used. For deletions, overlap of the homologous regions with the elements of the RepeatMasker track of the CHM13 reference^5,20,38^ (obtained from https://github.com/marbl/CHM13) was calculated with bedtools^102^ (v.2.31.1) intersect. Deletions were classified as repeat-mediated if at least 85% of the bases of the homologous regions on both sides intersect a RepeatMasker annotation of the same class and at least one homologous region spans 85% of the RepeatMasker element. For insertions the homologous sequences outputted by blastn were annotated with RepeatMasker and insertions where both homologous flanks show a reciprocal overlap of at least 85% with a RepeatMasker annotation of the same class were deemed repeat-mediated. To analyse segmental duplication (SD) mediated deletions, all deletions were intersected with the SD track of CHM13 and hits spanning more than 85 % of the homologous region and spanning more than 200 bases or 50 % of the SD were termed potentially SD mediated.

### Deletion recurrence analysis

To detect potential Alu-mediated deletions recurrence, we analysed deletions shorter than 5kbp in length exhibiting a flanking sequence homology of 300-400 bp. Additionally, to focus the analysis on potentially recurrent events, we limited our analysis to deletions with allele frequency between 40-60%. Furthermore, we used Hardy-Weinberg equilibrium and Mendelian consistency statistics to exclude events with potential genotyping errors. Lastly, we filtered out the deletions for which the phasing information was unavailable. Using the above mentioned criteria, we selected a subset of 11 deletions for our analysis. To find the evidence of recurrence, we used an approach similar to one previously devised to detect recurrent inversions^52^. We screened SNPs with at least 10% allele frequency lying within a 20kbp window centred around the deletion in search for positions where we observe both SNP alleles in haplotypes with and without the deletion. Additionally, we used centroid hierarchical clustering to cluster the SNP haplotypes in a 100kbp window centred around the deletion, in an effort to find whether the haplotypes with and without the deletion appear together in similar clusters, a phenomenon that indicates that the event recurred across human populations.

