## Supplementary Figures and Notes for "Long-read sequencing and structural variant characterization in 1,019 samples from the 1000 Genomes Project"

### Supplementary Notes

#### Supplementary Note 1

To allow capturing inversions of all sizes comprehensively in our resource, we explored inversion detection methods using simulations, by simulating inversions of varied sizes, ranging from 50 to 1000 bp. Simulating Oxford Nanopore Technologies (ONT) reads from a modified genome, we aimed for a coverage mirroring the coverage observed in actual samples - median coverage of 17X with reads averaging 20 kb in length. These reads were then aligned to a specified 250 kb segment of chromosome 1 from the hg38 reference genome using the minimap2 and NGMLR aligners, and as a next step, inversion detection was performed with Sniffles2 and Delly. **Fig. S34** shows that minimap2 exhibits a very high mismatch rate in regions containing inversions smaller than 1kb, whereas NGMLR alignments do not suffer from this drawback in small inversion regions. This superior performance of NGMLR, particularly in small and complex genomic regions, is attributed to its sophisticated scoring mechanism, tailored to accommodate the complexities of long reads[1]. Concurrently, **Fig. S46** illustrates that Delly is more effective than Sniffles2 in detecting inversions up to 1kb, suggesting a more suitable methodology for identifying small inversions.

#### Supplementary Note 2

**Targeted genotyping of challenging loci.** We used Locityper[2] to genotype a set of highly polymorphic and medically relevant loci for all 1,019 ONT datasets. The selected list of 270 target regions cover 15 Mb and overlap 463 protein coding genes, including 265 challenging medically relevant genes, characterized by the Genome in a Bottle Consortium[3]; 20 polymorphic MUC genes[4] and lipoprotein(a)-encoding *LPA* gene[5].

For each locus and each ONT dataset, Locityper produced two local haplotypes based on the sequencing data and its database of locus alleles. Additionally, we reconstructed haplotypes from the phased NYGC call set; and compared two sets of haplotypes against the phased whole genome assemblies for 8 HGSVC and 1 HPRC samples. As a measure of genotyping accuracy, we evaluated sequence divergence between pairs of actual and predicted haplotypes.

At 191/270 (70.1%) polymorphic regions Locityper haplotypes were at least 0.1% more accurate than NYGC-based haplotypes (≥1% improvement at 96 loci). In contrast, NYGC call set outperformed Locityper by 0.1% at only 7 loci (by 1% at 4 loci). Locityper improved genotyping accuracy by over 5% at 18 loci, covering 985 kb and completely encompassing 23 protein coding genes, including 8 mucin genes and tandem-repeat rich genes *GPI* and *LPA*. Therefore, our ONT based framework considerably outperforms the prior 1kg dataset for making genotype assessments in these medically relevant regions.

#### Supplementary Note 3

**Phasing Quality and Haplotype tagging of ONT reads using a high-coverage short read 1kG reference.** We used WhatsHap[6] to phase the raw genotypes of NYGC against the statistical phasing done by the same to assess the quality of the ONT reads. We observed an excellent agreement of the long-read based phasing to the NYGC statistical phasing[7] with an average switch error rate of 1.24% (**Fig. S5 - panel a**). The trio-based phasing and the long-read trio based phasing for the four trios also showed good agreement. The trio phasing had an average switch error rate of 0.84% and 0.21% for the parents and the children respectively. The long-read trio phasing had an average switch error rate of 0.93% and 0.35% for the parents and children respectively (**Fig. S5 - panel b**). We observe a phased block length with a median NG50 of 1,578,731 bp, as well as an excellent agreement with previous population based phasing[7], indicating that the phased panels from NYGC are a good resource to tag the reads and provide additional information to the SV discovery and SV genotyping tools.

We used WhatsHap's[6] haplotagging command, which uses the information from the given phased panel to tag the read coming either from haplotype 1 or haplotype 2. The ONT reads were haplotype-tagged (haplotagged) using the NYGC statistical phased VCF using WhatsHap's haplotag function. 69.9% of the ONT reads were tagged on average (**Fig. S47**) and utilised downstream by SVarp and Giggles to improve SV calling and genotyping.

#### Supplementary Note 4

SV **characteristic** breakdown for different variant categories. The final phased callset of the SAGA pipeline showed high abundance of low allele frequency variants with 58.78% of variants having minor allele frequency of less than 1%. Breaking this observation further down into variant categories, we observed 58.57% of insertions, 65.13% of deletions and 46.56% of putatively complex SVs had allele frequency less than 1% in their respective categories.

Using the variant annotations using SVAN (Methods), we further investigated the abundance of rare variants based on annotated class. All variants annotated as 'COMPLEX_DUP', 'DUP', 'INV_DUP', and 'DUP_INTERSPERSED' were considered as one category of 'DUP'. Similarly, all insertions annotated as 'solo', 'partnered', and 'orphan' were considered as 'MEI (Non-reference)' and deletions of similar annotations were considered as 'MEI (Reference)'. We observed that variants categorized as 'DUP', 'MEI (Non-reference)', 'NUMT', and 'PSD' had high abundance of rare alleles with 67.62%, 75.11%, 88.72%, and 85.92% of variants from each category respectively had MAF < 1%. Variants categorized as 'MEI (Reference)', 'VNTR Expansion' and 'VNTR Contraction' had 29.89%, 36.46%, and 34.92% of variants from respective categories with MAF < 1% (Fig. S31).

### Supplementary Figures

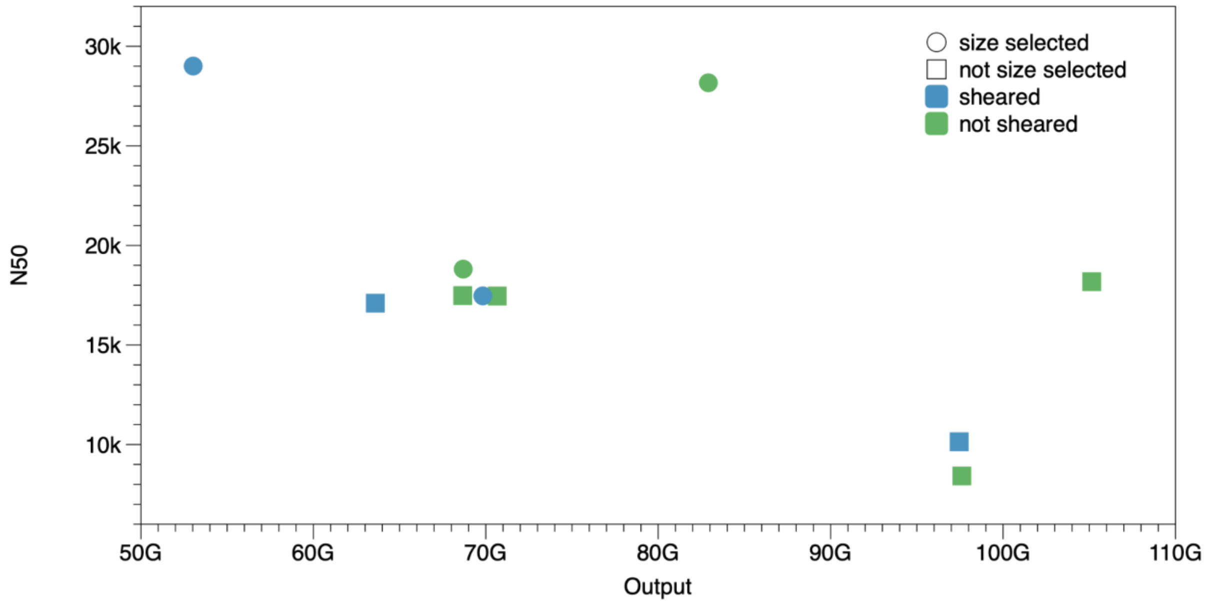


**Supplementary Figure 1:** Read-length N50 and output with and without size-selection for ≥25 kbp fragments and needle shearing of DNA.

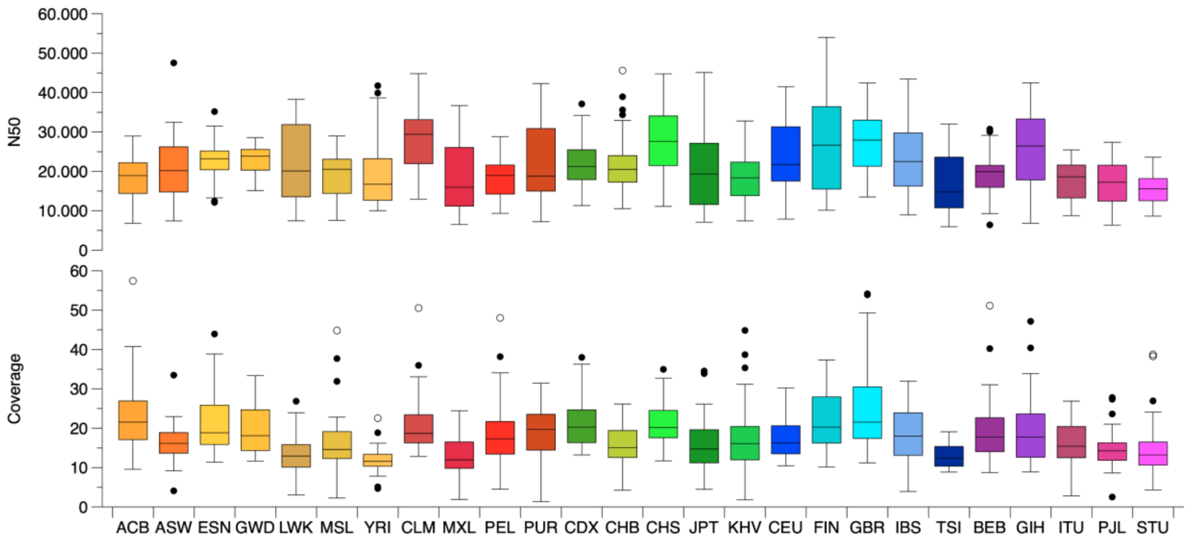


**Supplementary Figure 2**: Read-length N50 and fold-coverage for the 1,019 samples grouped by geographic location.

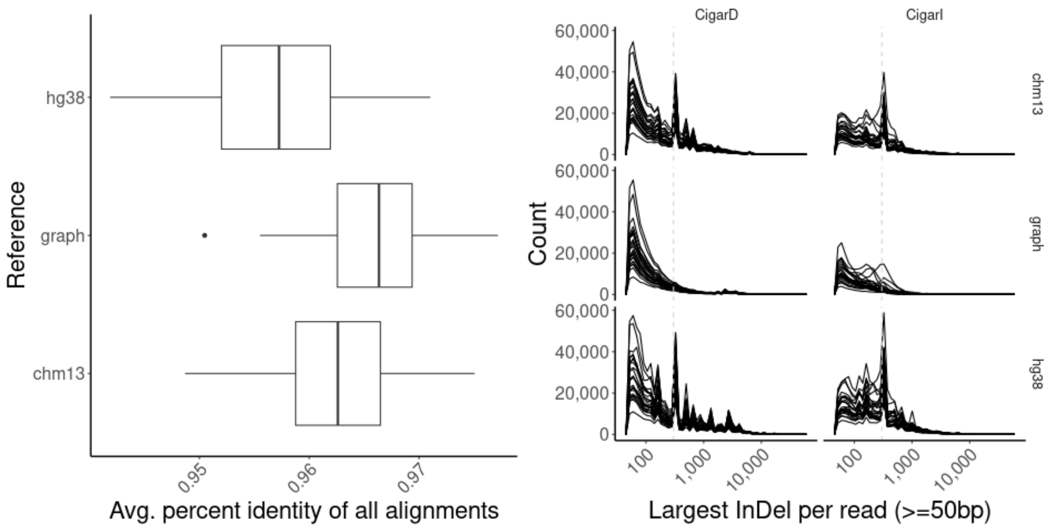


**Supplementary Figure 3: Left:** Average percent identity of all alignments (MAPQ > 0) of 25 random samples to the linear Genome Reference Consortium reference (hg38), the linear Telomere-to-Telomere consortium reference (chm13), and the pangenome minigraph reference from the HPRC (graph). **Right:** Distribution of the largest Cigar I (insertion) and Cigar D (deletion) operation for each read by sample. Vertical dashed line is at 300bp to indicate the expected ALU peak for linear reference genomes.

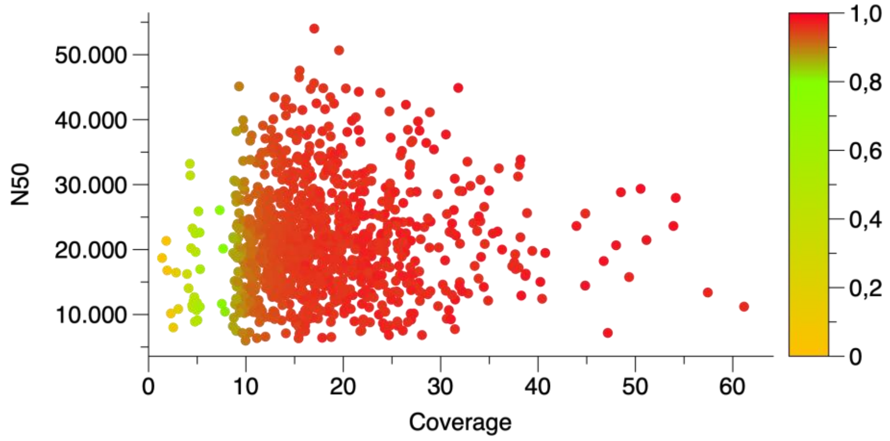


**Supplementary Figure 4:** Fold-coverage and read-length N50 for the 1,019 samples. Colors indicate the fraction of CHM13's bases covered at least five-fold.

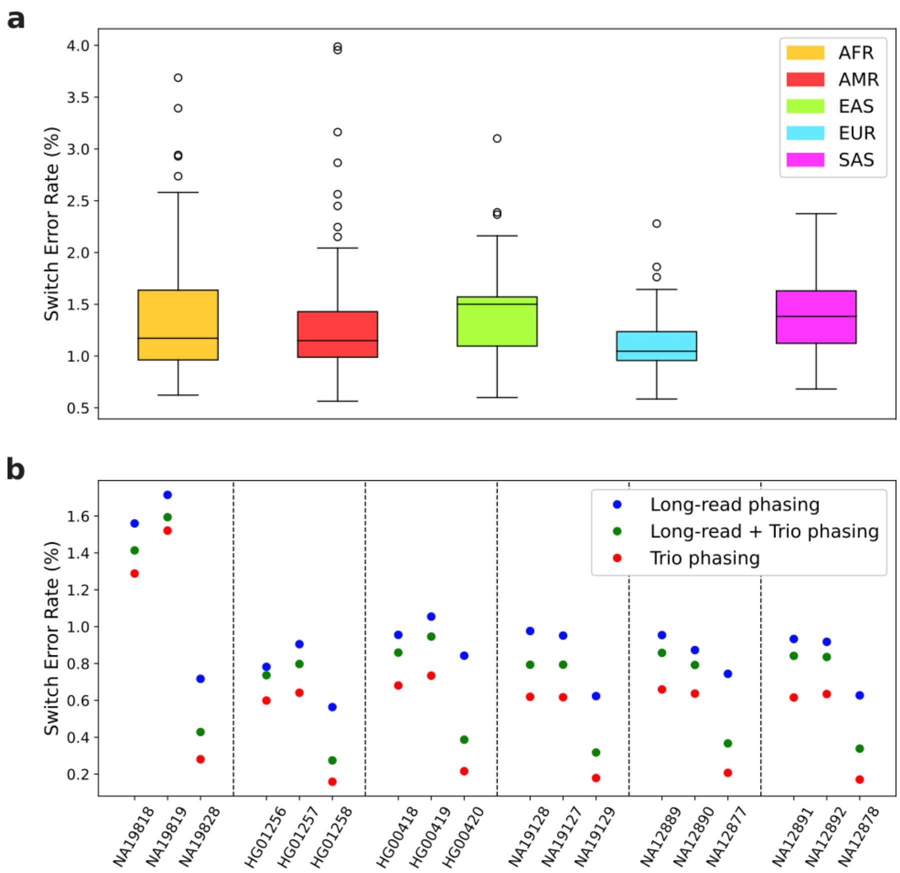


**Supplementary Figure 5:** Plotting the switch error rate (SER) between WhatsHap[6] phasing of the Byrska-Bishop et al., 2022 (Cell)[7] raw genotypes using the ONT reads from this study against the statistical phasing performed on the same genotypes. **a)** shows the SER for all the samples (grouped by population) for the comparison between long-read phasing and the statistical phasing. **b)** shows the SER for the samples in the 6 families for which ONT data is available. The SER is shown for the three different phasing strategies (long-read phasing, trio phasing and long-read trio phasing). The samples in each family are ordered father, mother and child.

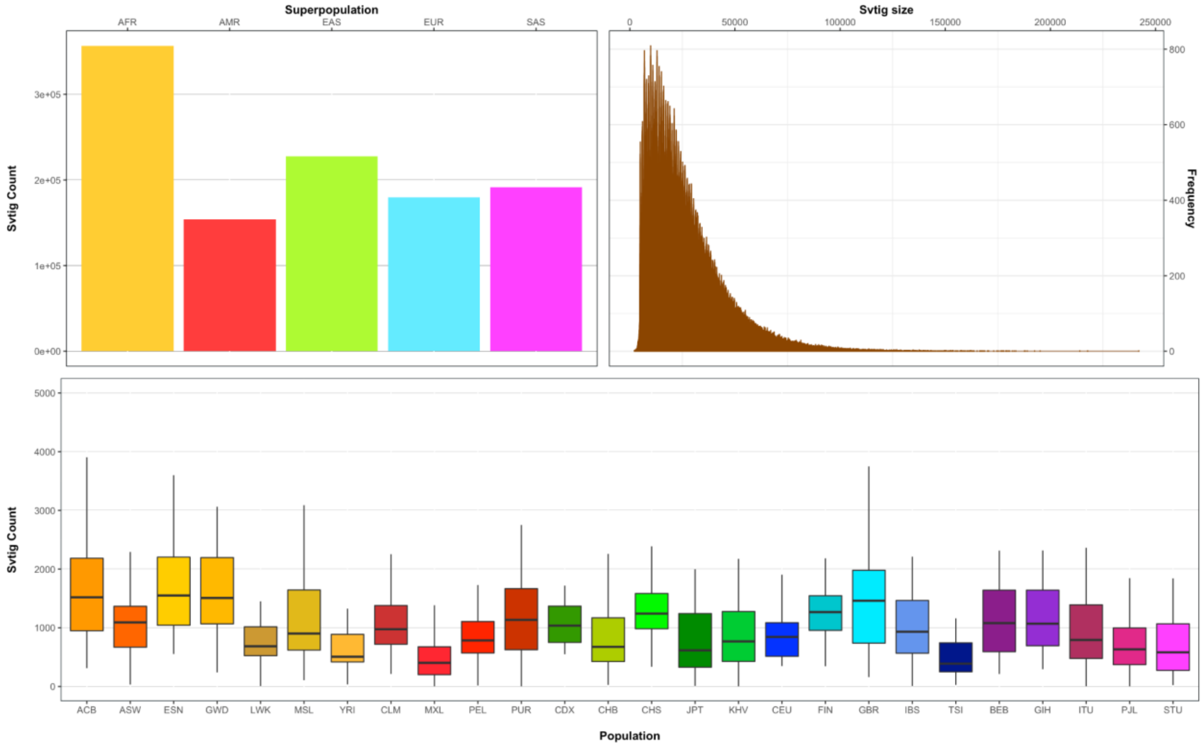


**Supplementary Figure 6:** Bar plot and the boxplot show svtig counts generated by SVarp per superpopulation and population respectively for 967 samples. The length frequency is also depicted by the histogram for the whole cohort. Note that numbers are based on the total number of svtigs in paternal and maternal haplotypes.

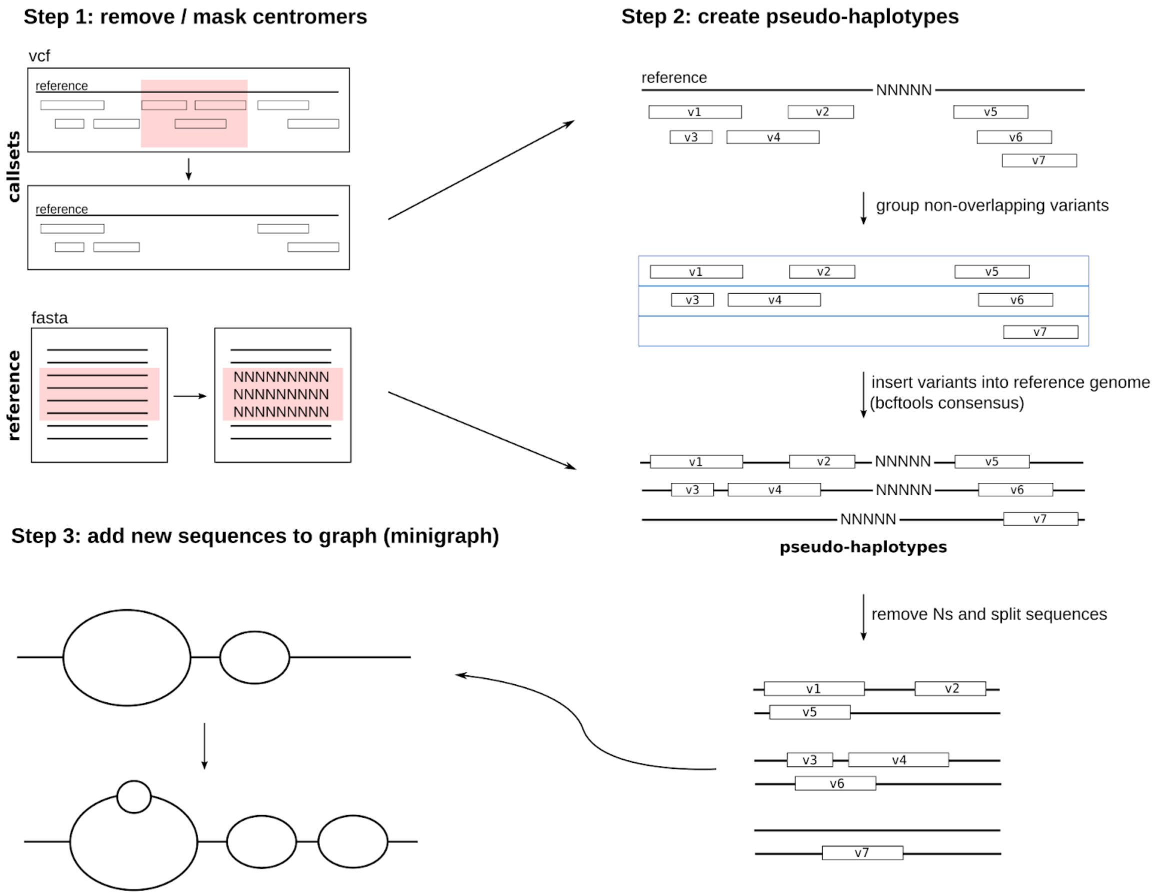


**Supplementary Figure 7**: Overview of our graph augmentation pipeline. **Step 1:** variant calls within centromere regions are removed and centromere regions are masked by Ns in the reference genome. **Step 2:** sets of non-overlapping variants are grouped and inserted into the reference genome to obtain "pseudo-haplotypes". **Step 3:** Pseudo-haplotypes are added to the graph using minigraph[8].

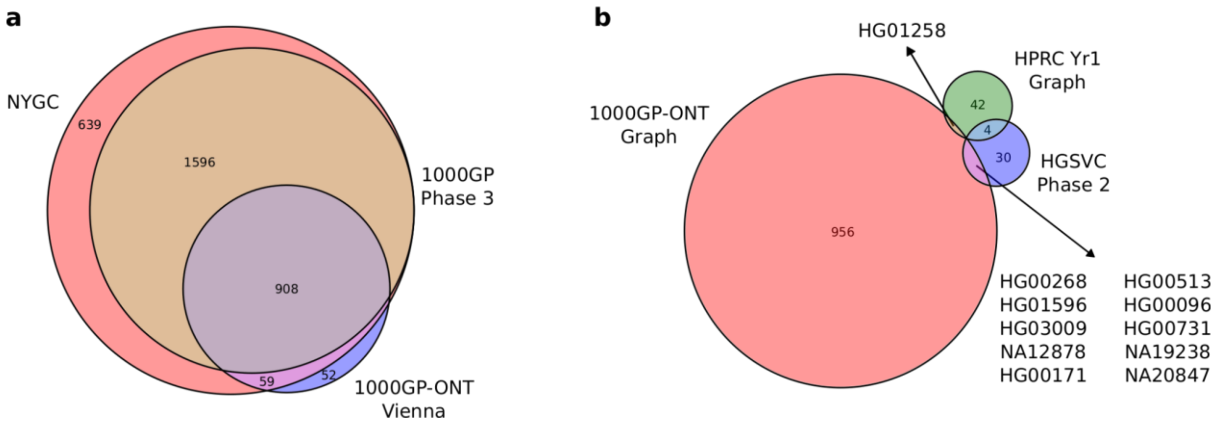


**Supplementary Figure 8: a)** Sample intersection of the 1019 samples of this study (1000GP-ONT Vienna) with the 2504 sample list of the 1000 Genomes Project Phase 3[17] and the New York Genome Center (NYGC)[7] study with 3202 samples (2504 samples along with 698 trios). **b)** For the Graph Methods, Giggles and SVarp, we utilise the NYGC phased panel and hence these methods are restricted to the intersection of the 1000 Genomes ONT Vienna Project and the NYGC sample set. This figure shows the intersection of this sample set with the HPRC Yr1[10] and HGSVC Phase 2[12] sample sets.

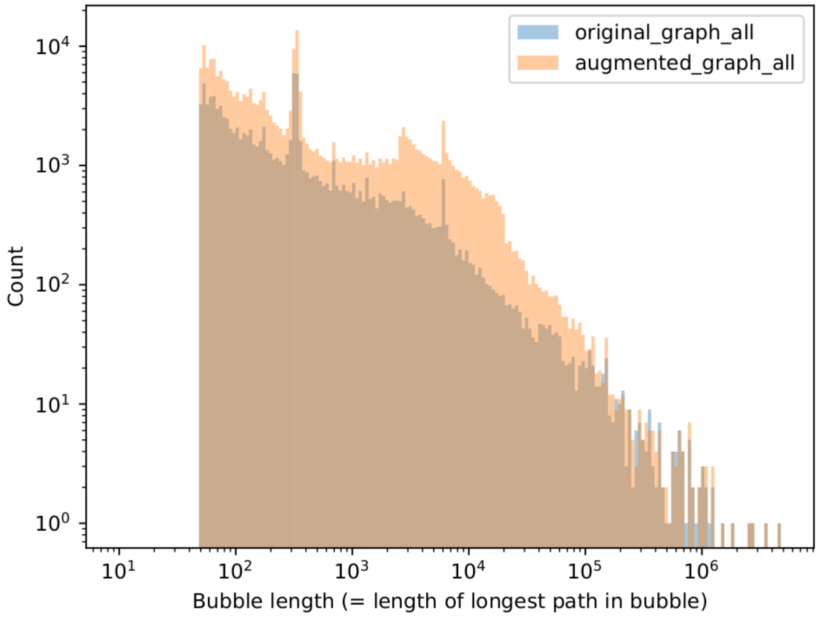


**Supplementary Figure 9:** Bubble-length histogram of the original and augmented graphs. The length of a bubble is defined as the length of the longest path through the bubble.

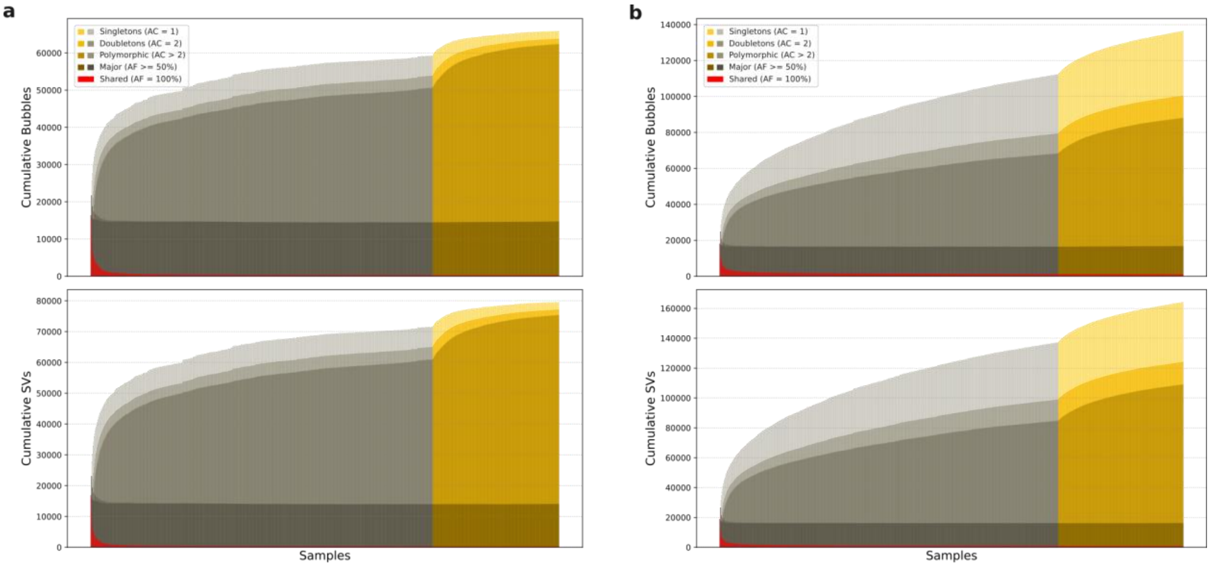


| | Shared | Major | Polymorphic | Doubletons | Singletons |
|---|---|---|---|---|---|
| Bubbles | 287 | 14,441 | 47,664 | 1,446 | 2,080 |
| SVs | 278 | 13,805 | 61,304 | 1,777 | 2,445 |

| | Shared | Major | Polymorphic | Doubletons | Singletons |
|---|---|---|---|---|---|
| Bubbles | 1,146 | 15,636 | 71,319 | 12,370 | 35,992 |
| SVs | 806 | 15,456 | 92,798 | 15,098 | 40,093 |

**Supplementary Figure 10:** Figure shows the cumulative growth curves for the SVs and the bubbles. **a)** shows the figures for the filtered genotypes of Giggles on the HPRC_mg and **b)** shows the figures for the final phased callset, which is the Giggles genotyped callset on the HPRC_mg_44+966 subsequently phased with ShapeIt5[11]. The tables below show the count of each category after all the samples have been considered. The figure has been made using the 908 samples from our callset which are unrelated. The yellow part of the figure is used to denote the addition of AFR samples while the grey part is for non-AFR samples. The non-AFR samples have been shuffled to reduce the effect of the non-AFR populations.

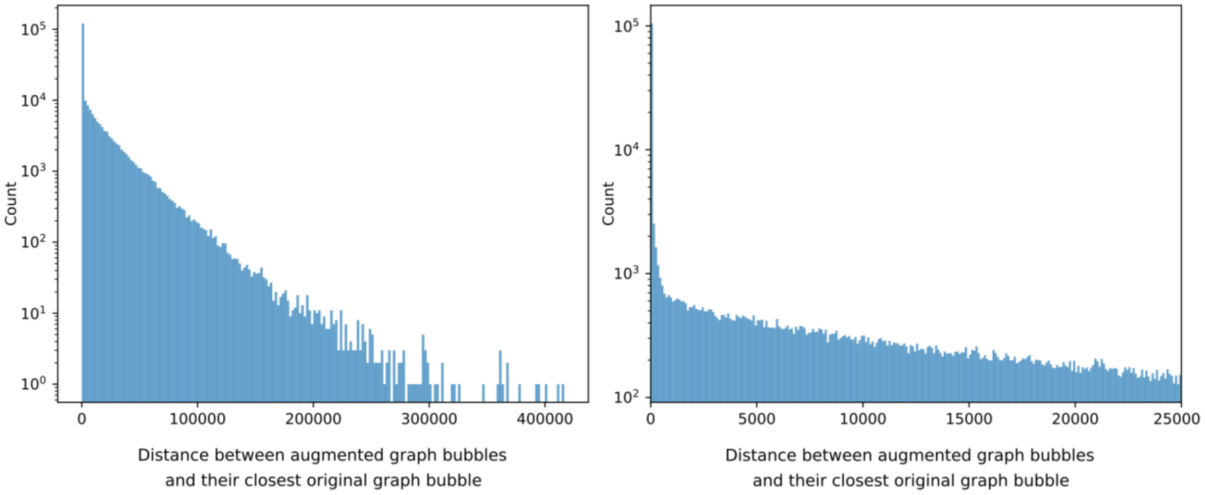


**Supplementary Figure 11:** Histogram showing the distances between bubbles in the augmented graph and their closest bubble in the original graph (in base pairs). The left panel shows the full histogram, the right panel shows only the part corresponding to distances up to 25,000 bp.

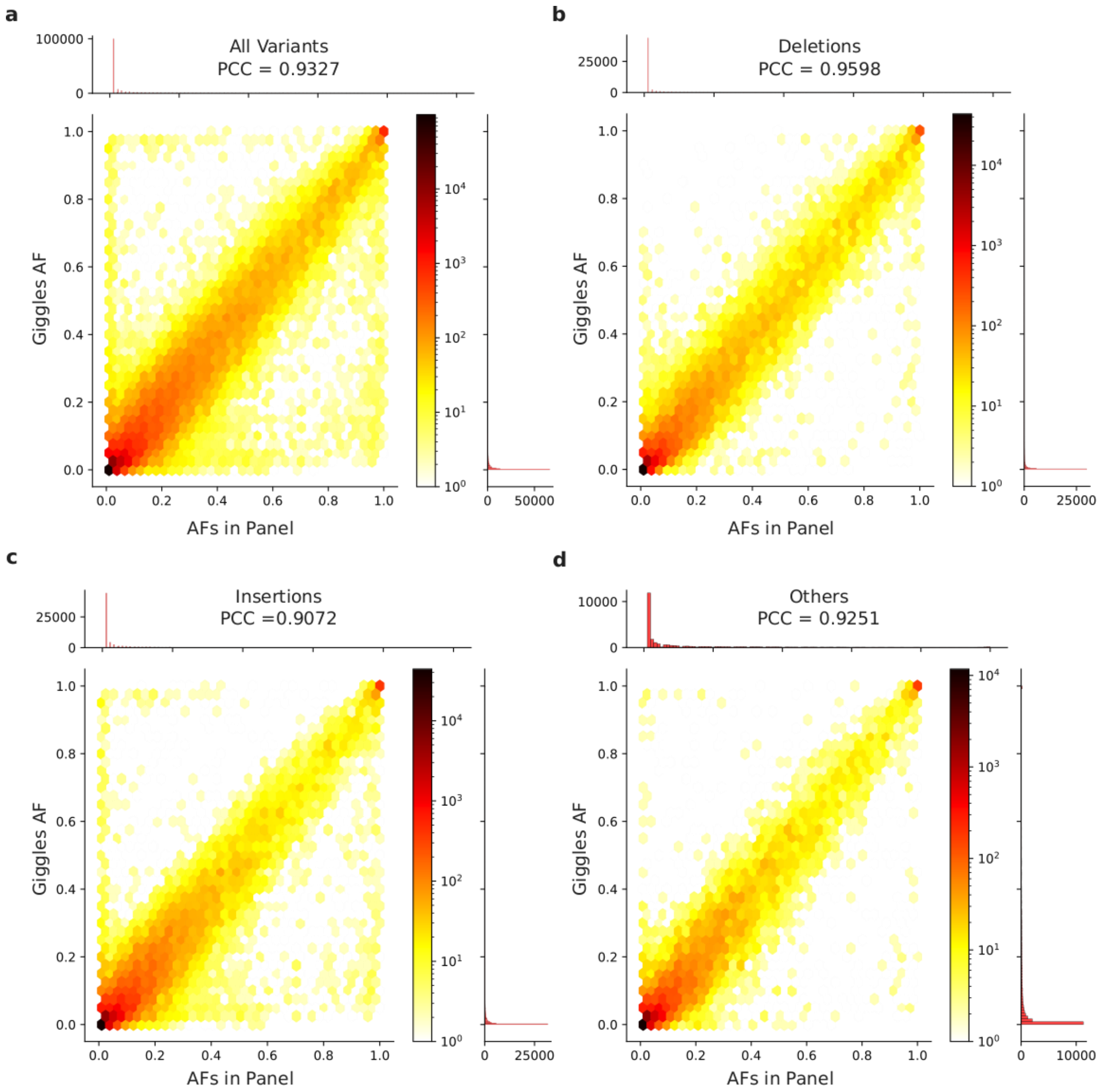


**Supplementary Figure 12:** The plot demonstrates the genotype quality of the genotypes by Giggles on the HPRC_mg_44+966 graph after filtering. Genotyping quality is shown here using a comparison of the allele frequency of an allele in the VCF panel (created using the HPRC[10] assemblies and the pseudohaplotypes of the SAGA framework) with the allele frequency of the same allele genotyped by Giggles in the callset (using only the 908 unrelated samples from our callset). The plot has been broken into variant types: **a)** shows the plot for all variants, **b)** shows deletions, **c)** shows insertions and **d)** shows the rest of the variants which strictly do not fall into deletions or insertions. The Pearson's Correlation Coefficient (PCC) for each plot has been provided in the figure.

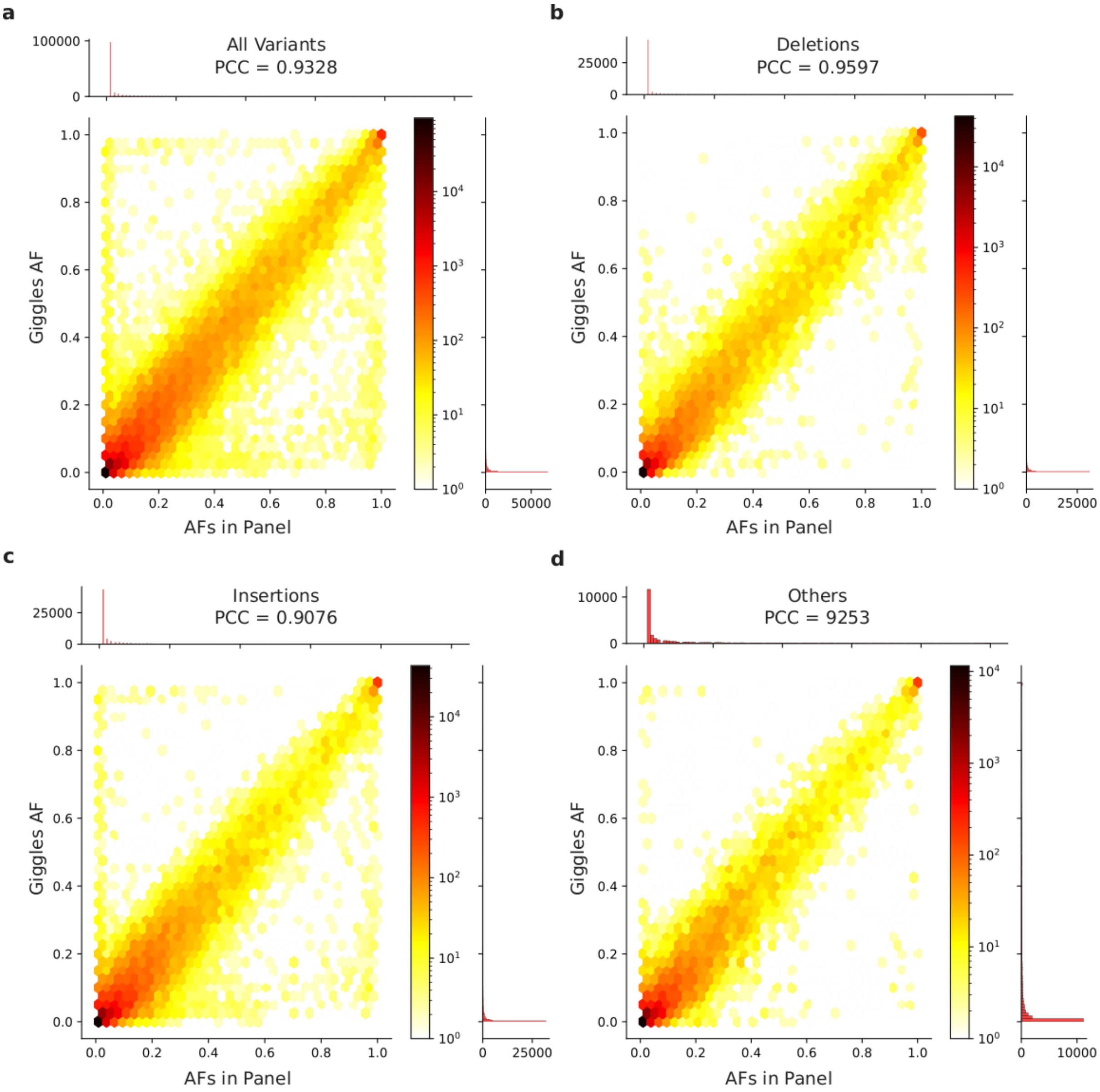


**Supplementary Figure 13:** The plot demonstrates the genotype quality of the genotypes by Giggles on the HPRC_mg_44+966 graph which have been phased using ShapeIt5[11] and filtered. Genotyping quality is shown here using a comparison of the allele frequency of an allele in the VCF panel (created using the HPRC[10] assemblies and the pseudohaplotypes of the SAGA framework) with the allele frequency of the same allele genotyped by Giggles in the callset (using only the 908 unrelated samples from our callset). The plot has been broken into variant types: **a)** shows the plot for all variants, **b)** shows deletions, **c)** shows insertions and **d)** shows the rest of the variants which strictly do not fall into deletions or insertions. The Pearson's Correlation Coefficient (PCC) for each plot has been provided in the figure.

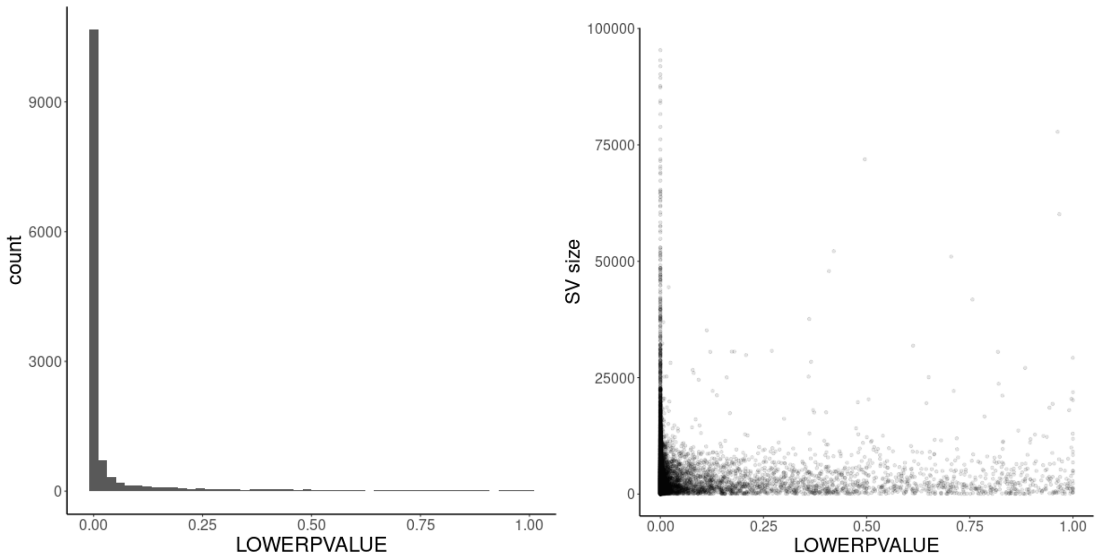


**Supplementary Figure 14:** Intensity rank sum (IRS) test[9] using SNP array probe intensity data. Left panel shows the p-value distribution for all deletions that can be assessed via IRS (13,788 deletions) and the right panel shows a scatter plot of p-value and SV size. Overall estimated FDR for deletions based on IRS is 8.06%.

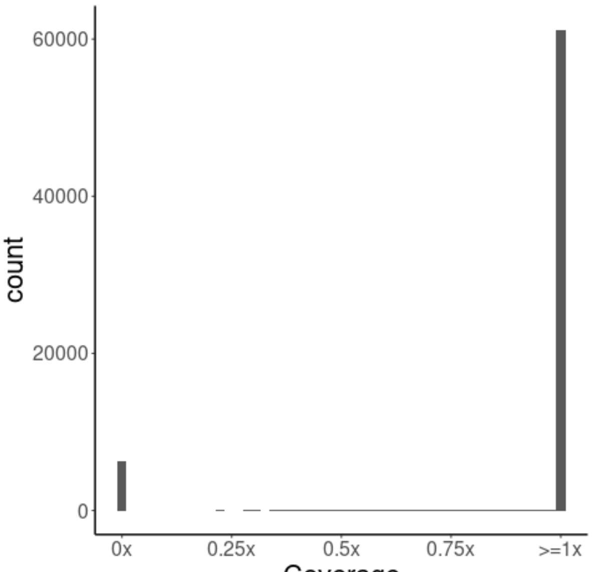


**Supplementary Figure 15:** Coverage analysis of the insertion sequences using the compacted de Bruijn graphs computed from prior short-read data[7]. For a singleton SV occurring in only one sample (allele count one) the expected coverage is 1x.

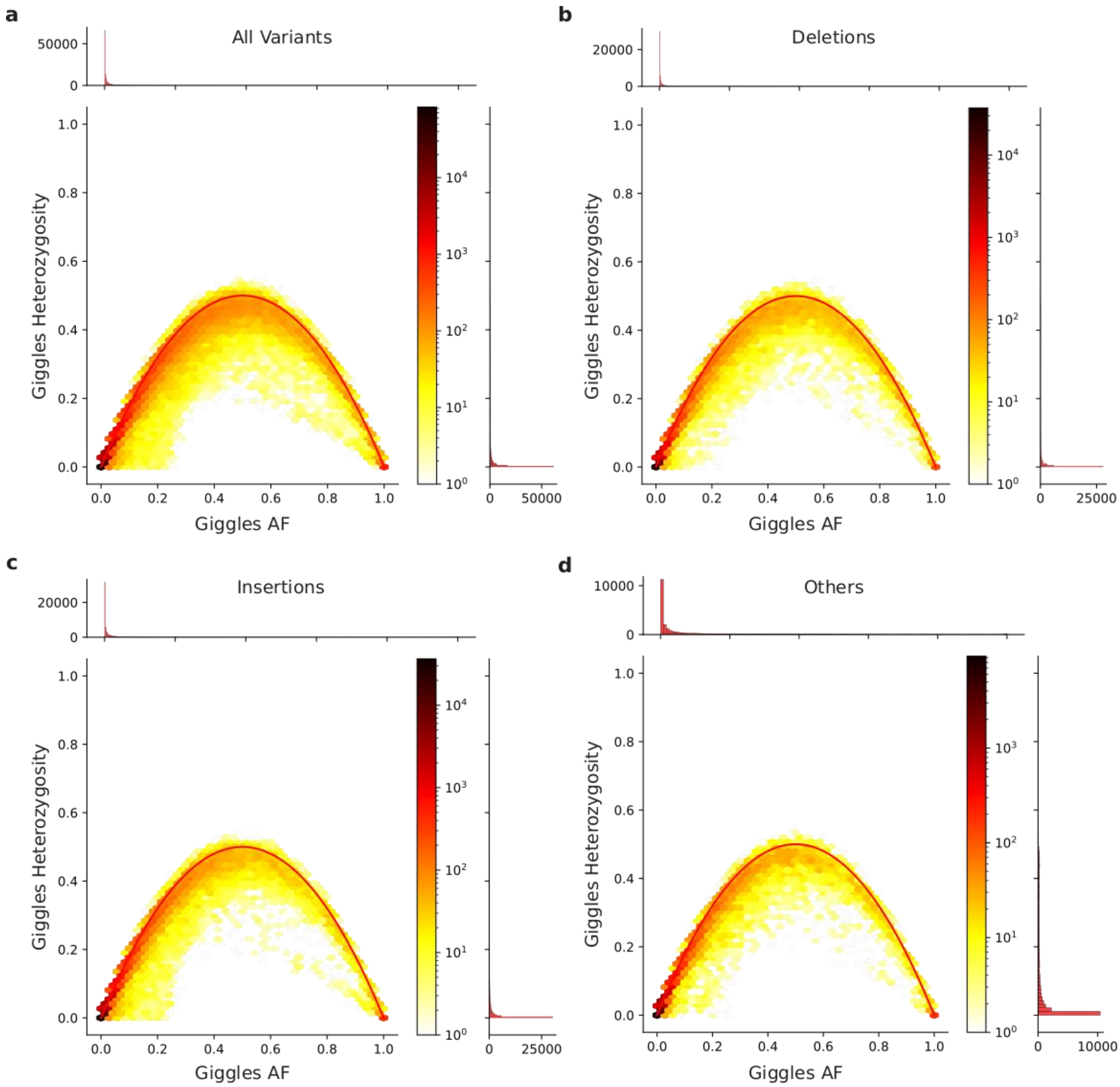


**Supplementary Figure 16:** The plot demonstrates the genotype quality of the genotypes by Giggles on the HPRC_mg_44+966 graph after filtering. Genotyping quality is shown here using a Hardy-Weinberg Equilibrium (HWE) plot given with the allele frequency of the genotyped allele and the percentage of samples heterozygous for that allele (using only the 908 unrelated samples from our callset). The plot has been broken into variant types: **a)** shows the HWE plot for all variants, **b)** shows deletions, **c)** shows insertions and **d)** shows the rest of the variants which strictly do not fall into deletions or insertions.

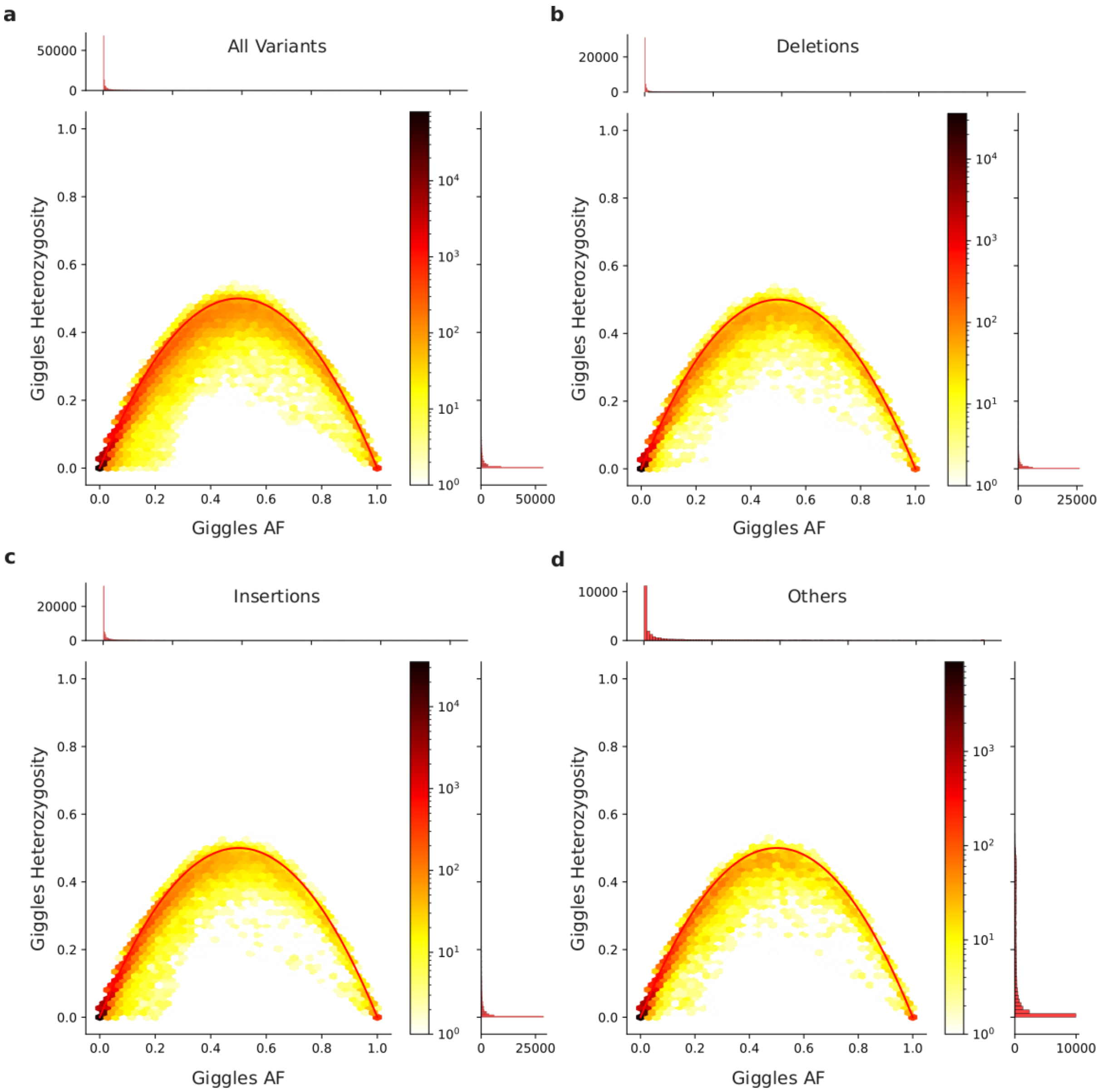


**Supplementary Figure 17:** The plot demonstrates the genotype quality of the genotypes by Giggles on the HPRC_mg_44+966 graph which have been phased using ShapeIt5 and filtered. Genotyping quality is shown here using a Hardy-Weinberg Equilibrium (HWE) plot given with the allele frequency of the genotyped allele and the percentage of samples heterozygous for that allele (using only the 908 unrelated samples from our callset). The plot has been broken into variant types: **a)** shows the HWE plot for all variants, **b)** shows deletions, **c)** shows insertions and **d)** shows the rest of the variants which strictly do not fall into deletions or insertions.

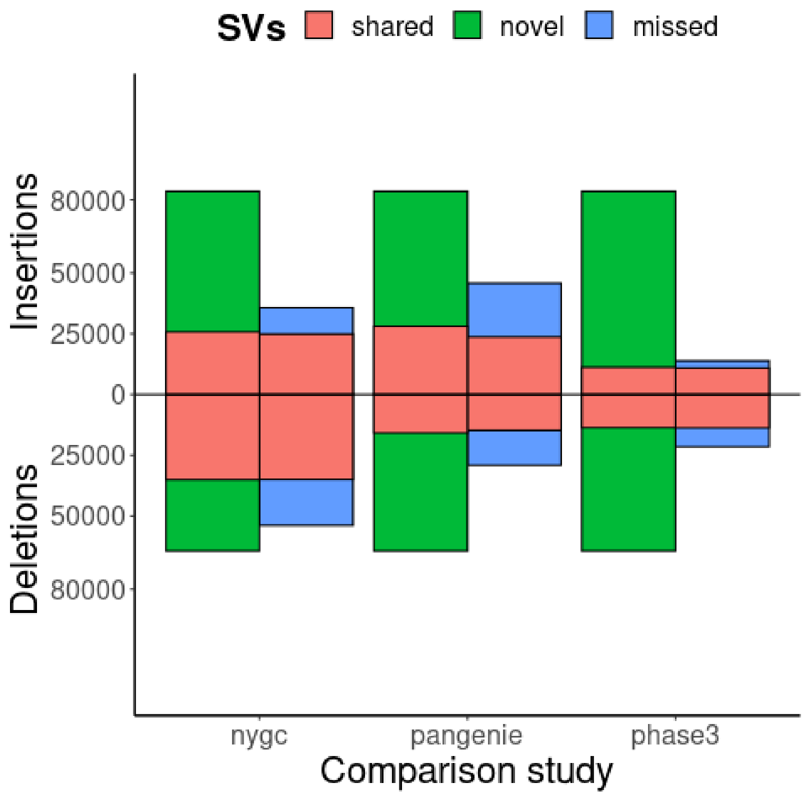


**Supplementary Figure 18:** The number of shared (red), novel (green) and missed (blue) SVs of our study (left bar) compared to prior SV studies (right bar) subsetted to samples present in our cohort. Comparison studies include deep-coverage short-read data generated by the New York Genome Center[7] (nygc), long-read data analyzed by the Human Genome Structural Variation Consortium project[12] and genotyped in the NYGC data using pangenie[13] (pangenie) and the 1kGP phase 3 SV callset[14] (phase3).

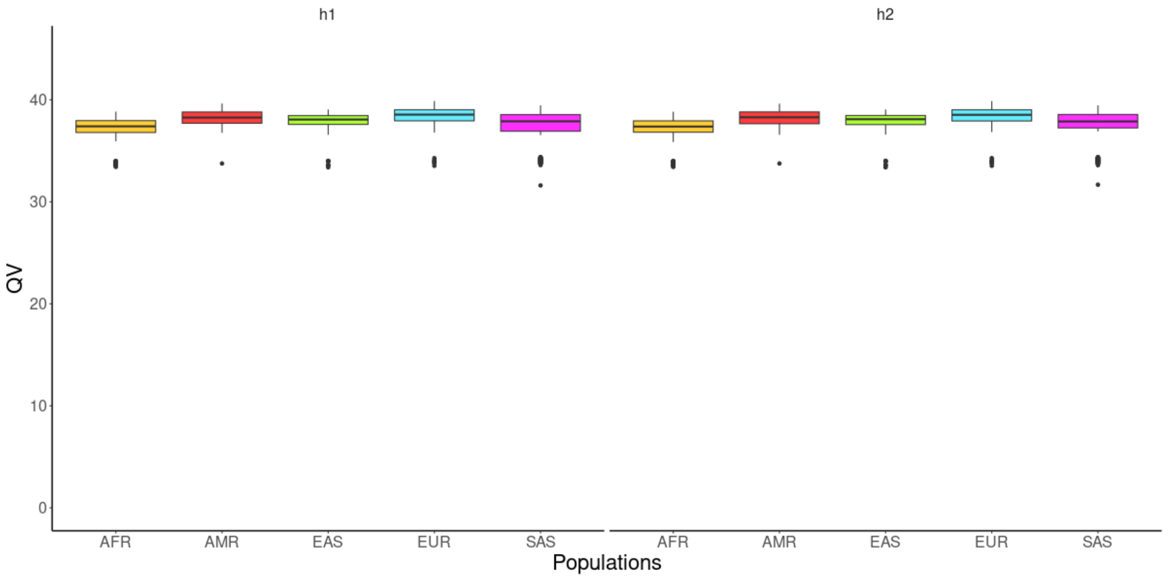


**Supplementary Figure 19:** Evaluation of variant completeness and phasing accuracy by integrating all variants (SNPs, InDels and SVs) into sample-specific haplotypes (h1 and h2) based on the CHM13 genome. Haplotype completeness and accuracy was evaluated using QV scores computed by yak[15] with k-mers of length 31 compared to the high-depth, short-read sequencing data. QV scores are lower compared to recently published high-quality HPRC assemblies[10] but in the range of high-depth *de novo* long-read assemblies[16].

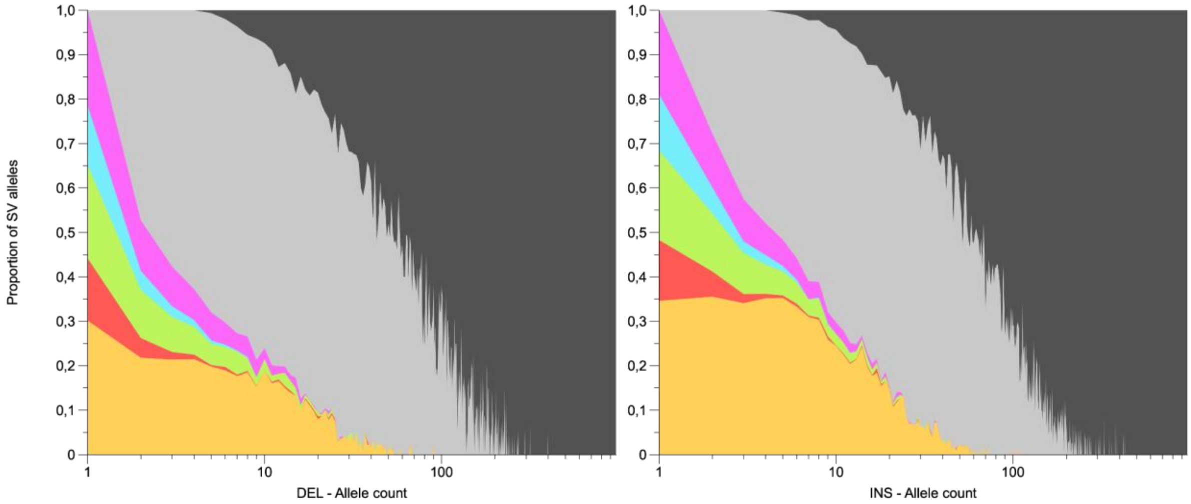

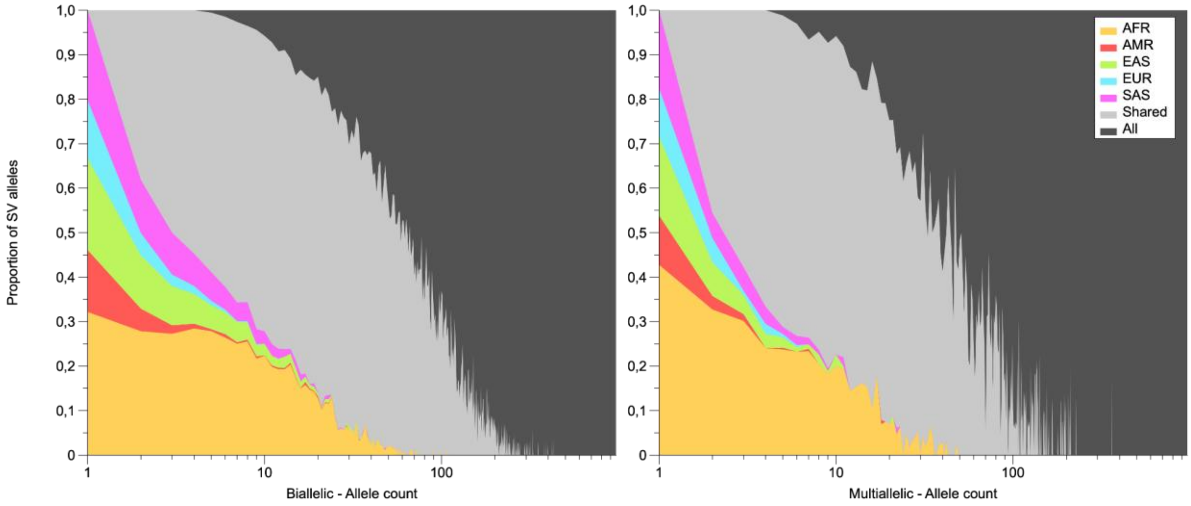


**Supplementary Figure 20:** SV allele sharing across continental populations. AFR, AMR, EAS, EUR and SAS exclusive to the respective continental group. Shared by at least two (and less then all) continental groups. All shared by all continental groups. Deletions (top left), insertions (top right), all biallelic SVs (bottom left), all multiallelic (SVs).

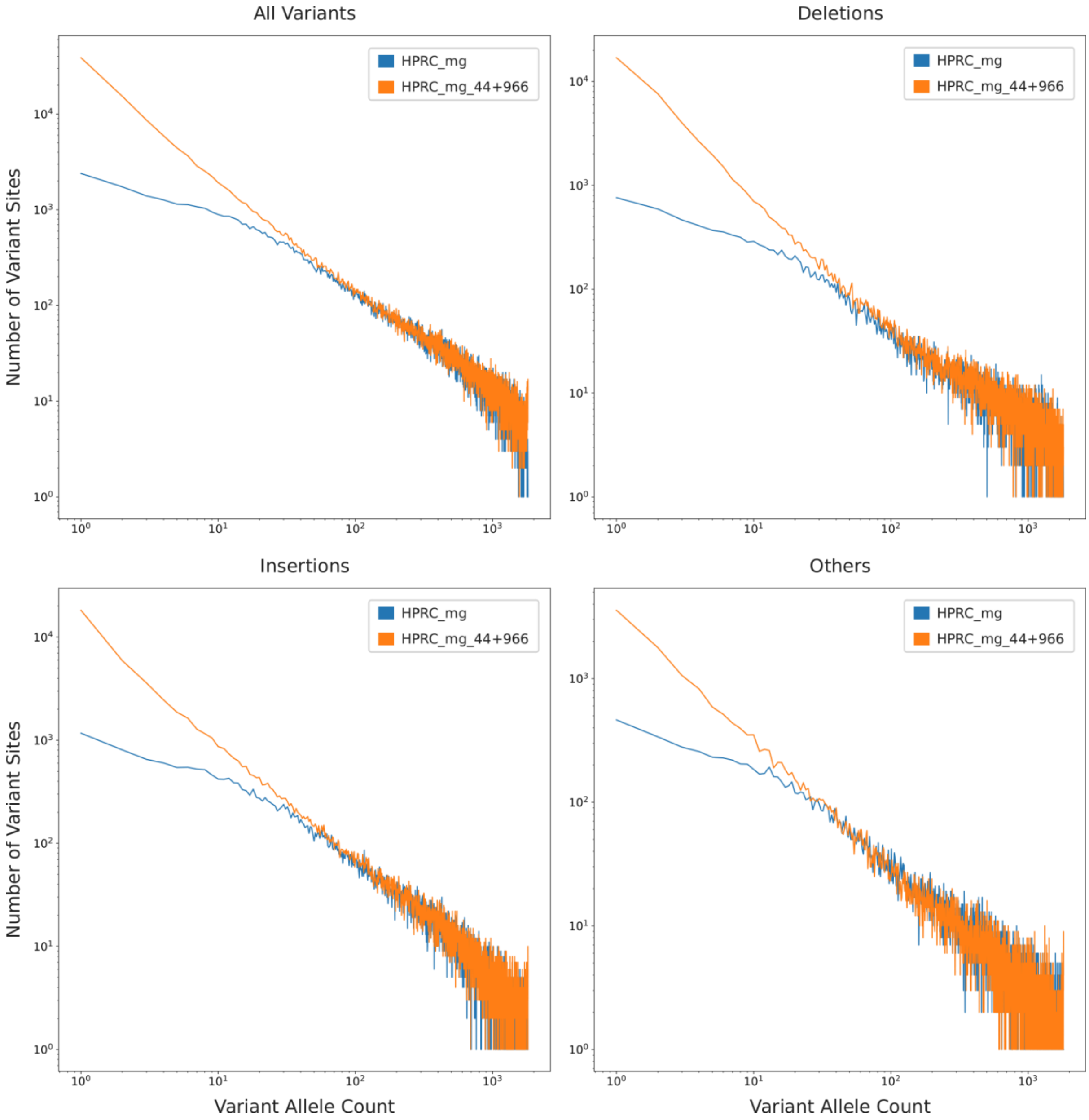


**Supplementary Figure 21:** The figure shows the relation between Variant Allele Count and the Number of Variant Sites with that allele count in the logarithmic space. In blue, the filtered genotypes of the Giggles callset on the HPRC_mg graph is shown and orange shows the final phased callset, which is the Giggles genotyped callset on the HPRC_mg_44+966 subsequently phased with ShapeIt5[11]. The plot has been broken into variant types: **a)** shows the plot for all variants, **b)** shows deletions, **c)** shows insertions and **d)** shows the rest of the variants which strictly do not fall into deletions or insertions.

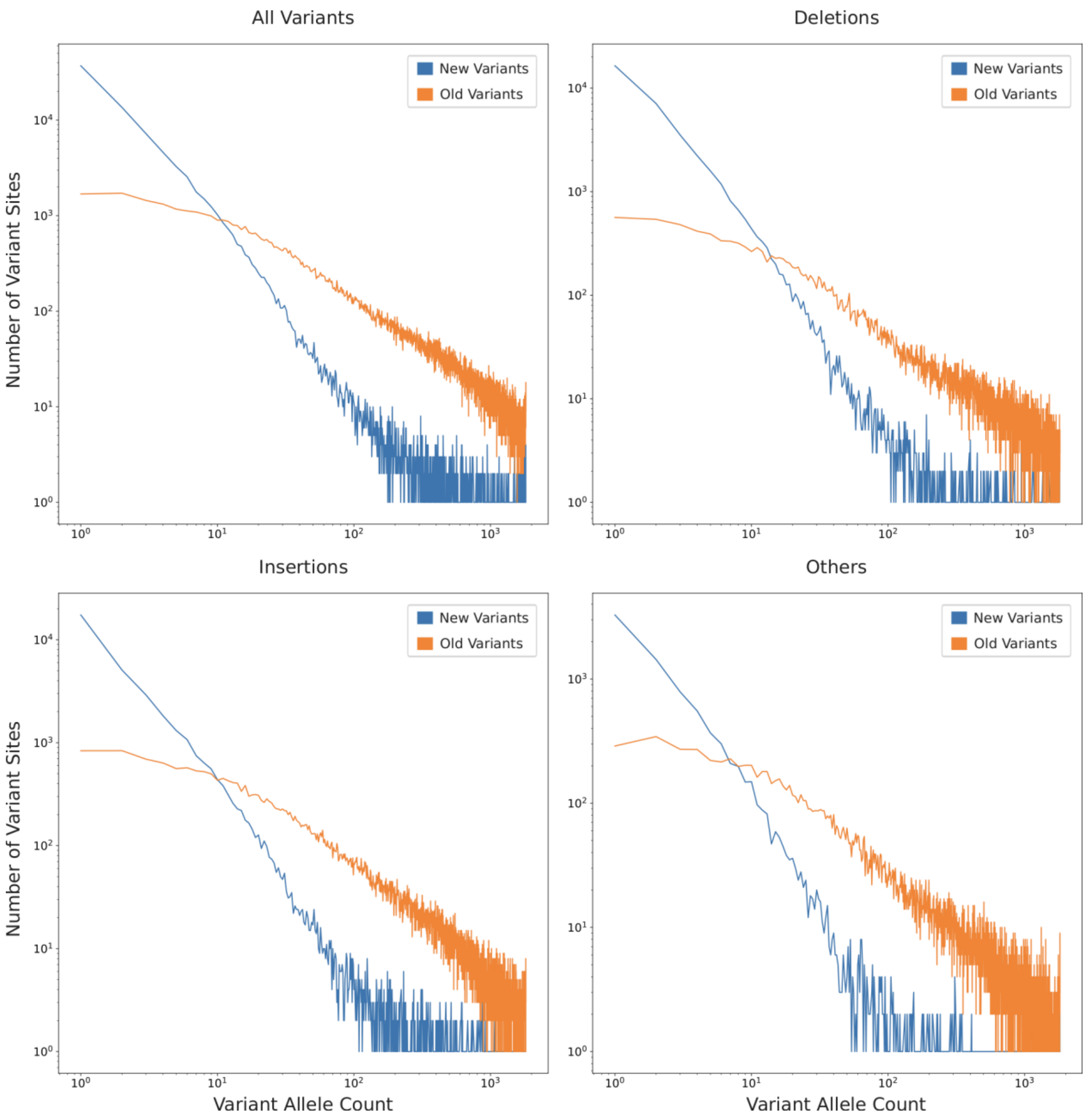


**Supplementary Figure 22:** The figure shows the relation between Variant Allele Count and the Number of Variant Sites with that allele count in the logarithmic space. The data shown here are the genotypes of the final phased callset, which is the Giggles genotyped callset on the HPRC_mg_44+966 subsequently phased with ShapeIt5[11]. The blue line is the new variants, determined by alleles exclusively found in the pseudo-haplotypes and the orange line is the old variants, determined by alleles exclusively found in the HPRC[10] assemblies. The plot has been broken into variant types: **a)** shows the plot for all variants, **b)** shows deletions, **c)** shows insertions and **d)** shows the rest of the variants which strictly do not fall into deletions or insertions.

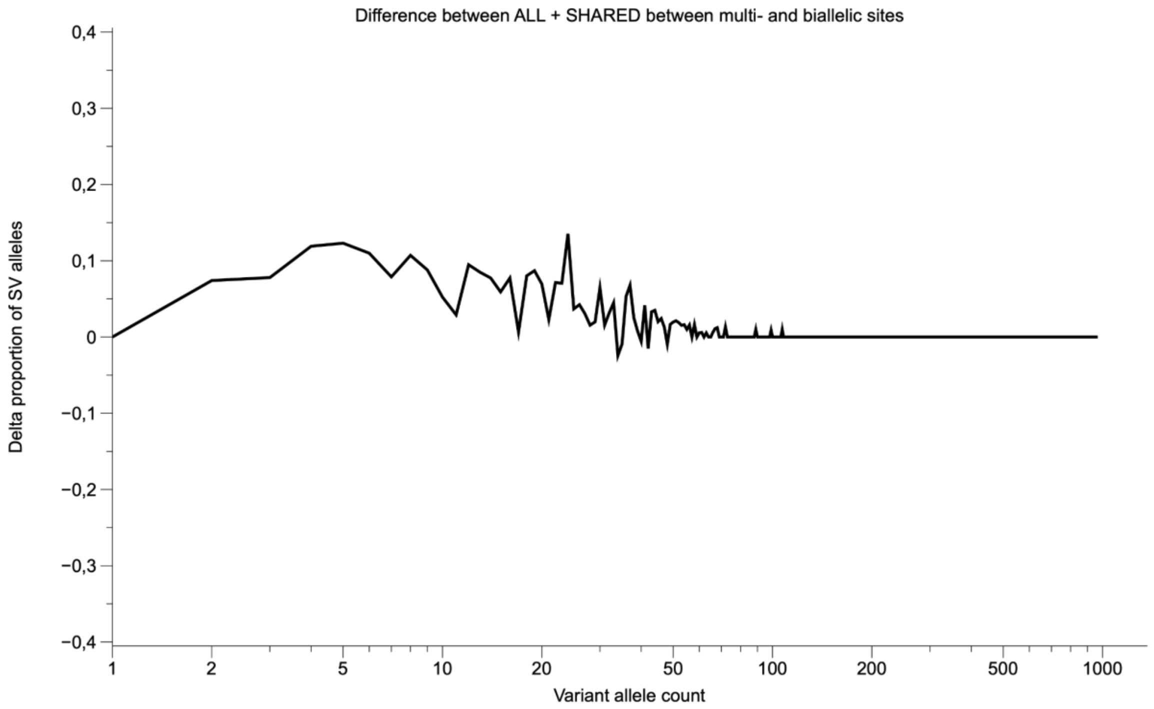


**Supplementary Figure 23:** Difference between the fraction of SVs shared across at least two continental populations (ALL = shared across all continental populations, SHARED = shared across at least two but not all continental populations) for biallelic and multiallelic sites. Multiallelic SVs have a higher propensity to be shared across continents than biallelic SVs.

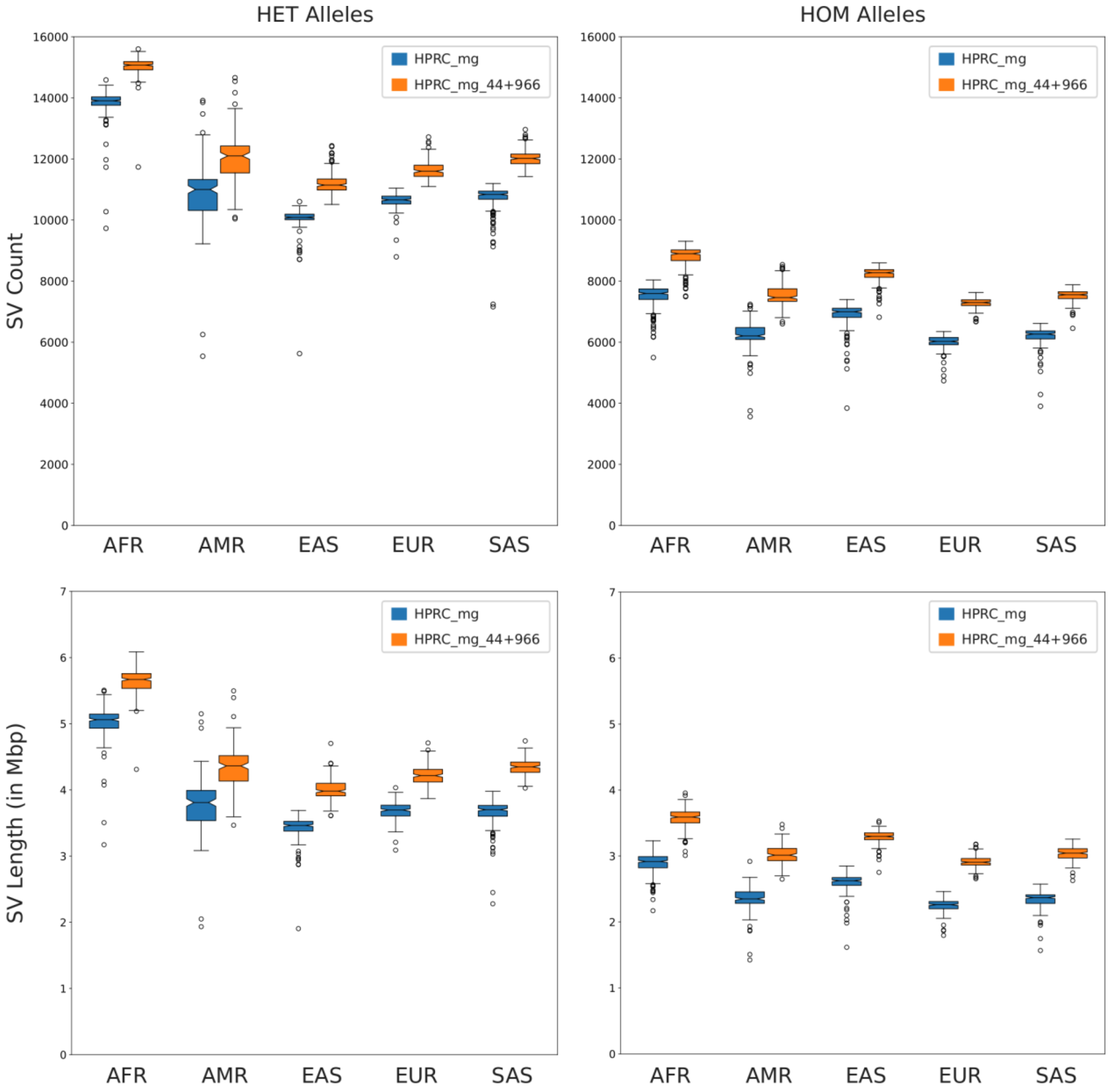


**Supplementary Figure 24:** The plot shows the distribution of SV counts for each sample (top row) and the cumulative length of the present SVs in the sample (bottom row). The variants have been divided into HETs and HOMs to denote heterozygous alleles (left column) and homozygous alternate alleles (right column). In blue, the filtered genotypes of the Giggles callset on the HPRC_mg graph is shown and orange shows the final phased callset, which is the Giggles genotyped callset on the HPRC_mg_44+966 subsequently phased with ShapeIt5[11].

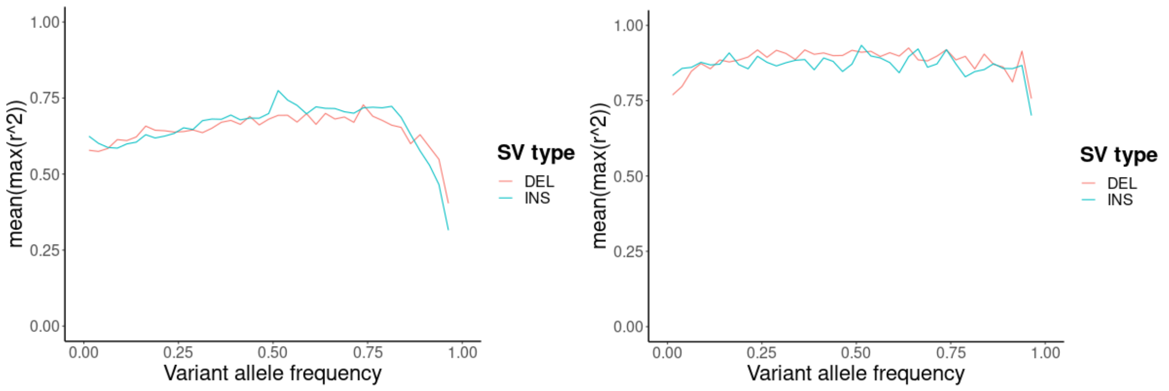


**Supplementary Figure 25:** Linkage disequilibrium (LD) of SVs (MAF >= 1%) with nearby single nucleotide polymorphisms (SNPs). The left panel shows all SVs whereas the right panel shows the subset of SVs in Genome in a Bottle high-confident regions of the CHM13 genome (2.3Gbp, 74.2%).

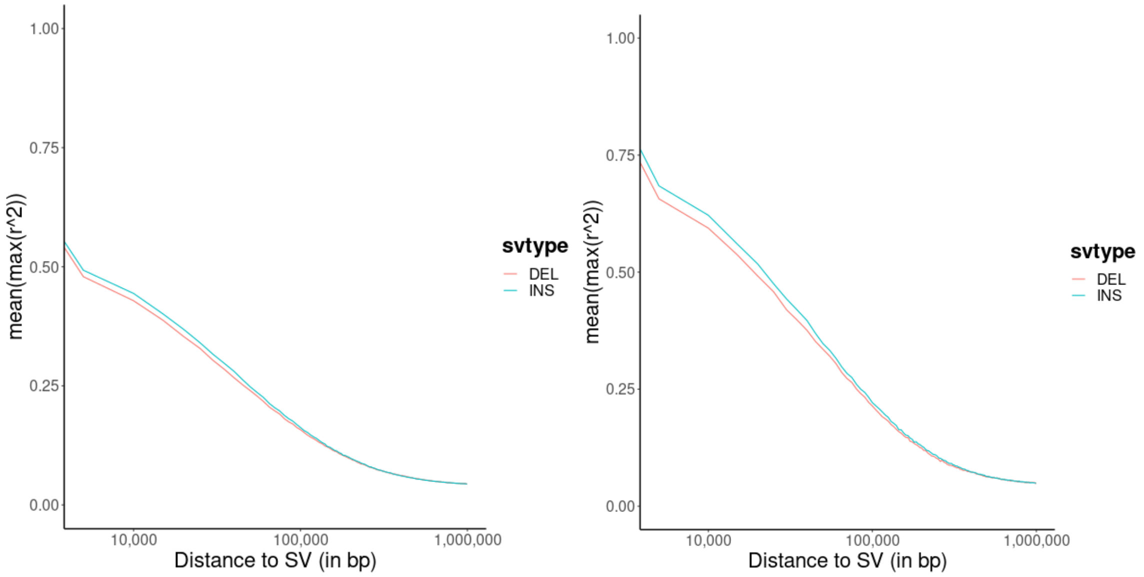


**Supplementary Figure 26:** Linkage disequilibrium (LD) of SVs (MAF >= 1%) with nearby single nucleotide polymorphisms (SNPs) as a function of the distance of the SV to the SNP. The left panel shows all SVs whereas the right panel shows the subset of SVs in Genome in a Bottle high-confident regions of the CHM13 genome (2.3Gbp, 74.2%).

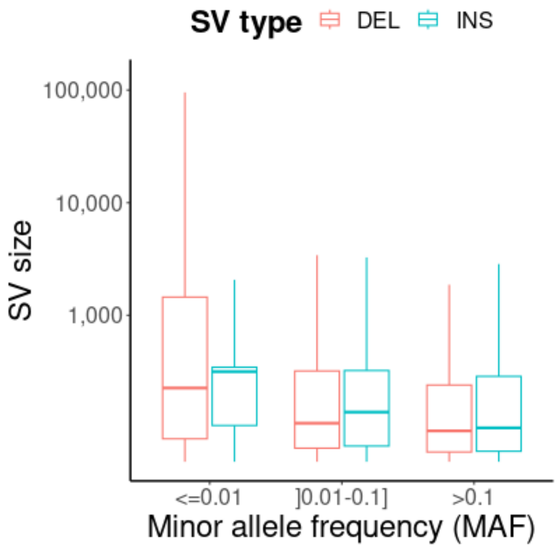


**Supplementary Figure 27:** SV size distribution by minor allele frequency (MAF) showing that common insertions and deletions tend to be shorter compared to rare SVs.

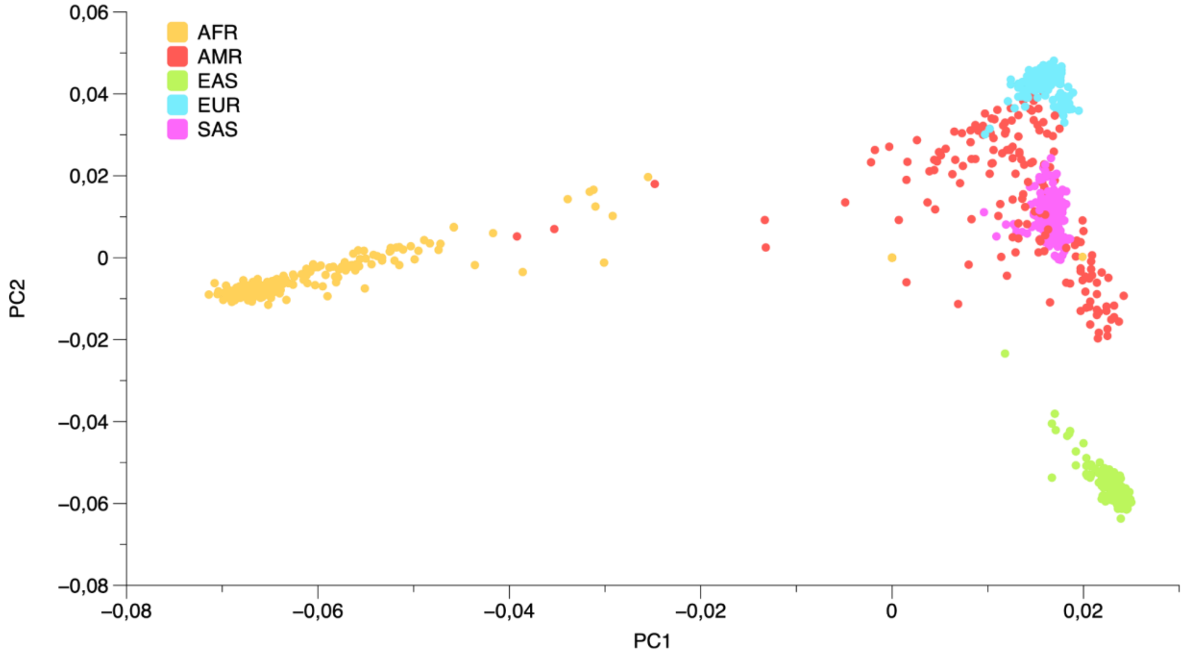


**Supplementary Figure 28:** Principal component analysis using insertions and deletions.

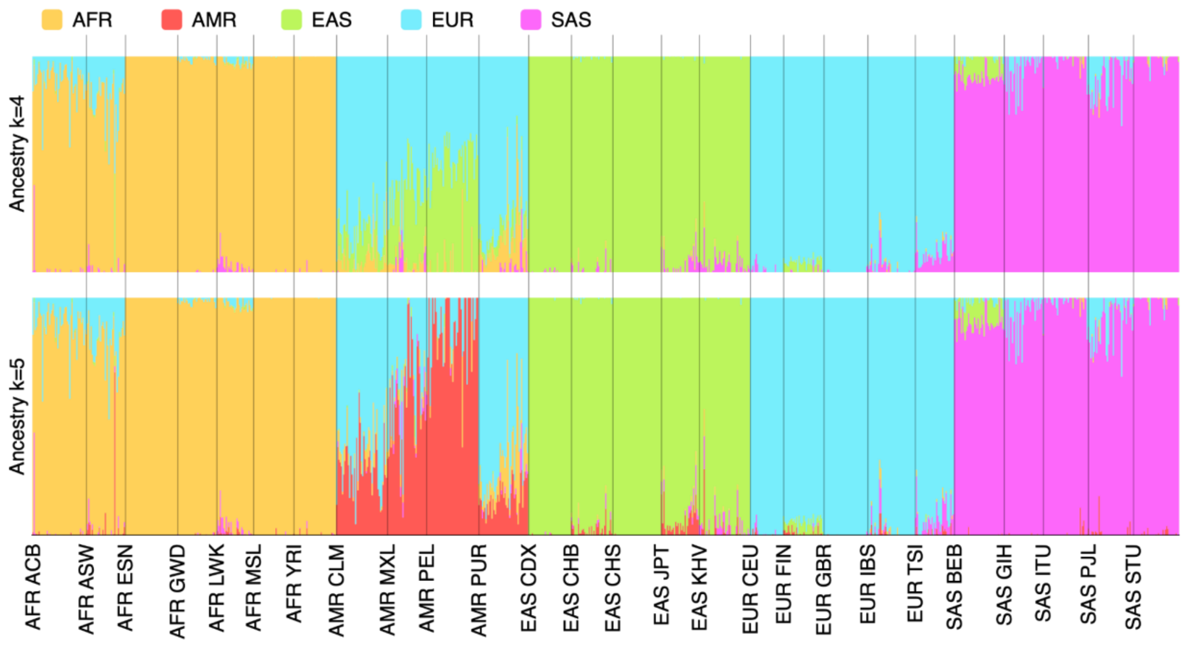


**Supplementary Figure 29:** SV-based admixing spectra using k=4 (top) and k=5 (bottom) reference populations.

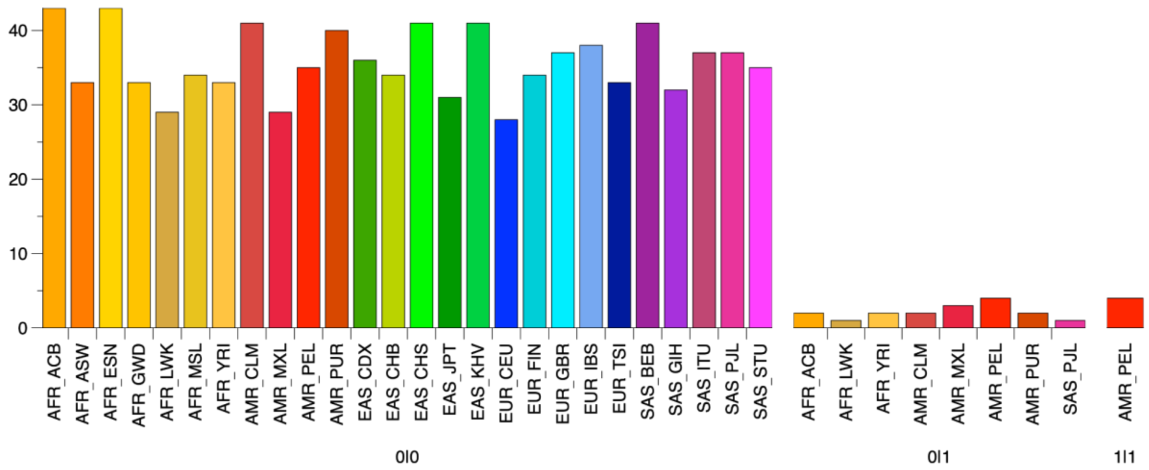


**Supplementary Figure 30:** Occurrences of SV *chr7-109295061-COMPLEX->s317119<s345452>s317121-246* (Fst 0.41) near *LAMB1* in the populations. The AMR populations in general show an enrichment of heterozygous copies. With homozygous copies occurring in PEL exclusively.

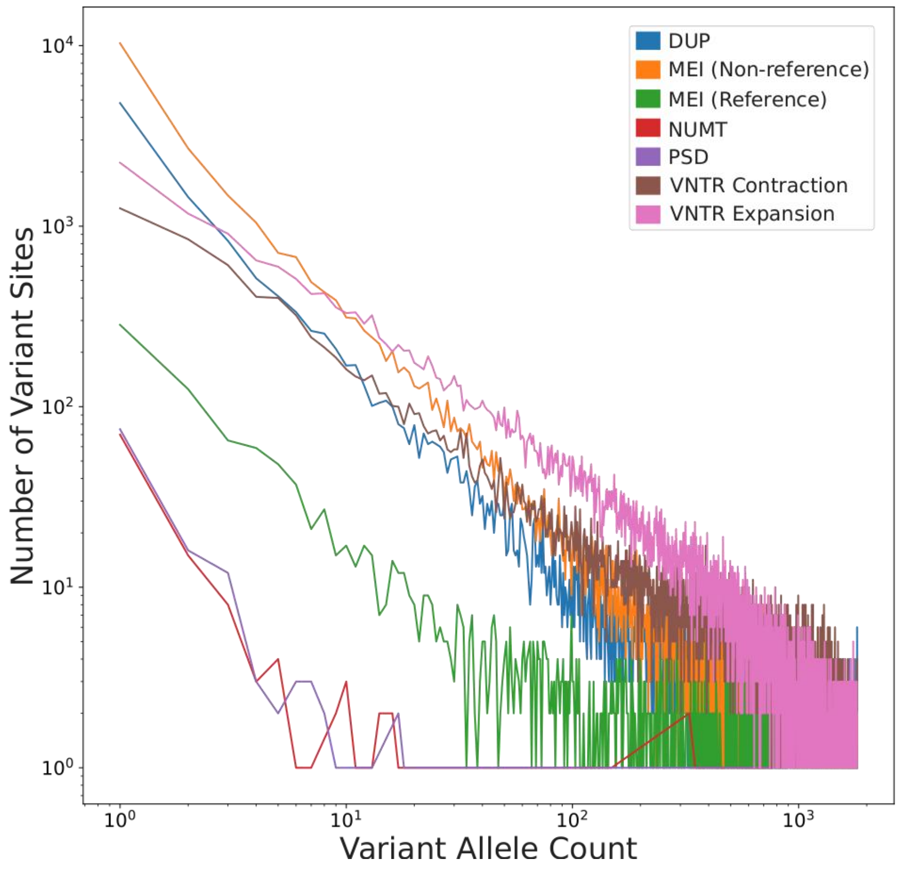


**Supplementary Figure 31:** The figure shows the relation between Variant Allele Count and the Number of Variant Sites with that allele count in the logarithmic space for the genotypes on the HPRC_mg_44+966 annotated by SVAN. Duplications (DUP), Mobile element insertions and deletions (MEI (non-reference) and MEI (reference), respectively), Nuclear mitochondrial DNA integration (NUMT), processed pseudogene integration (PSD), VNTR contraction and expansion.

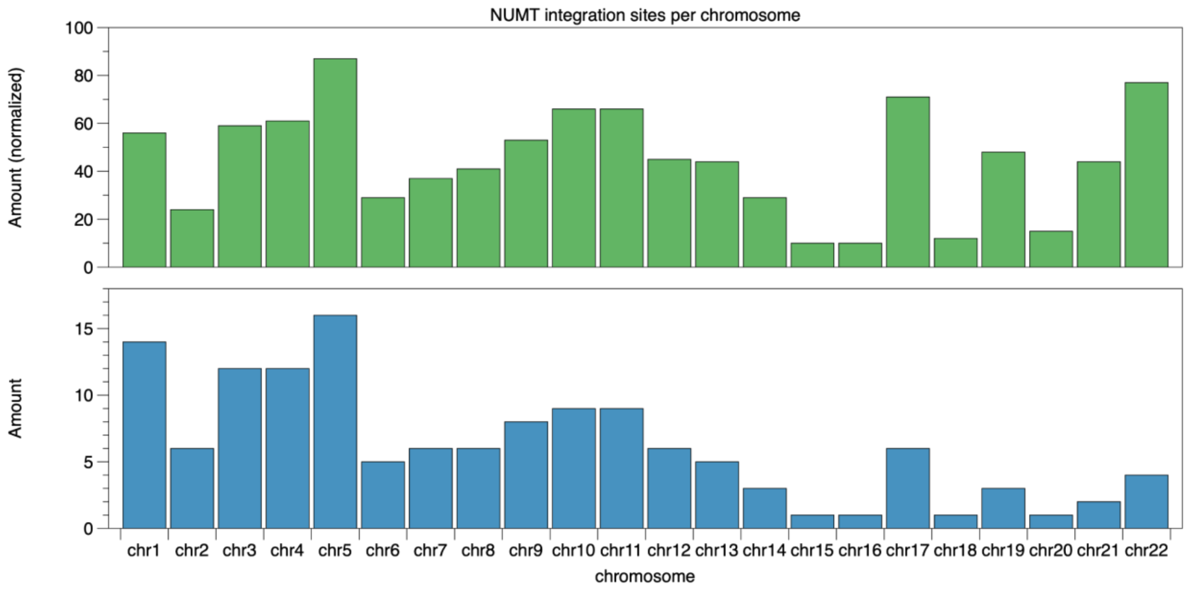


**Supplementary Figure 32:** Number of NUMT integration sites per chromosome. Bottom: number of sites. Top: number of sites normalized by chromosome length.

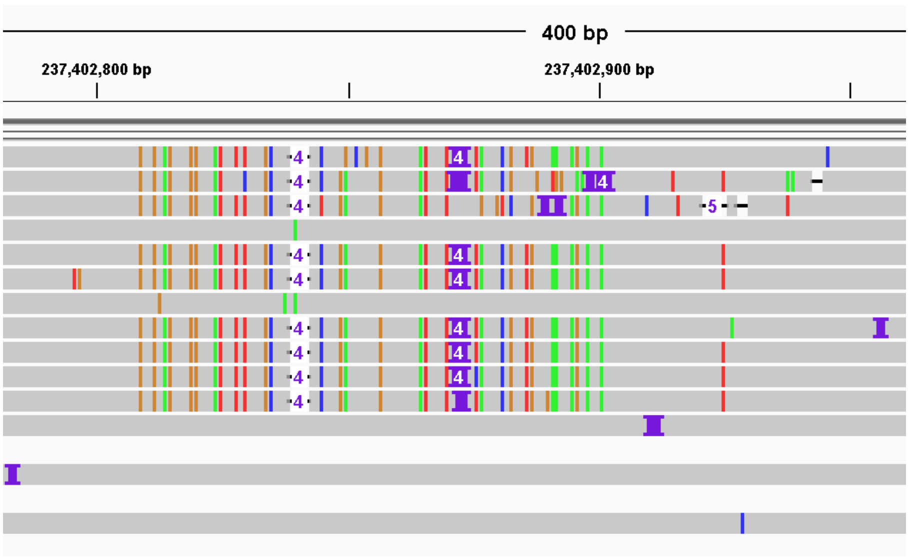


**Supplementary Figure 33**: Heterozygous inversion of length ~110 bp on chr1 in sample HG00107. Mapping with minimap2 to GRCh38 results in a forced high-mismatch alignment through the inversion.

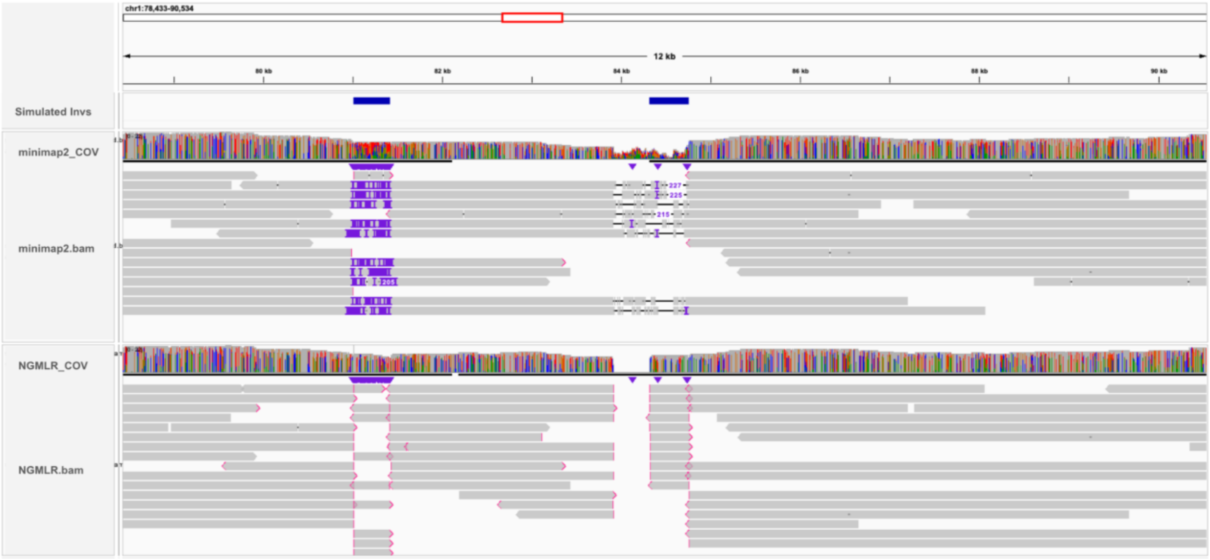


**Supplementary Figure 34: Comparative Analysis of NGMLR and minimap2 in small inversion detection.** Exemplary alignment outcomes by minimap2 and NGMLR within regions containing small inversions, approximately 500 base pairs each. It illustrates cases of a distinct inversion and an inversion adjacent to a small deletion. NGMLR demonstrates precise alignment in both instances, while minimap2 is characterized by the insertion of gaps and mismatches, thereby highlighting the superior performance of NGMLR in accurately aligning small inversions.

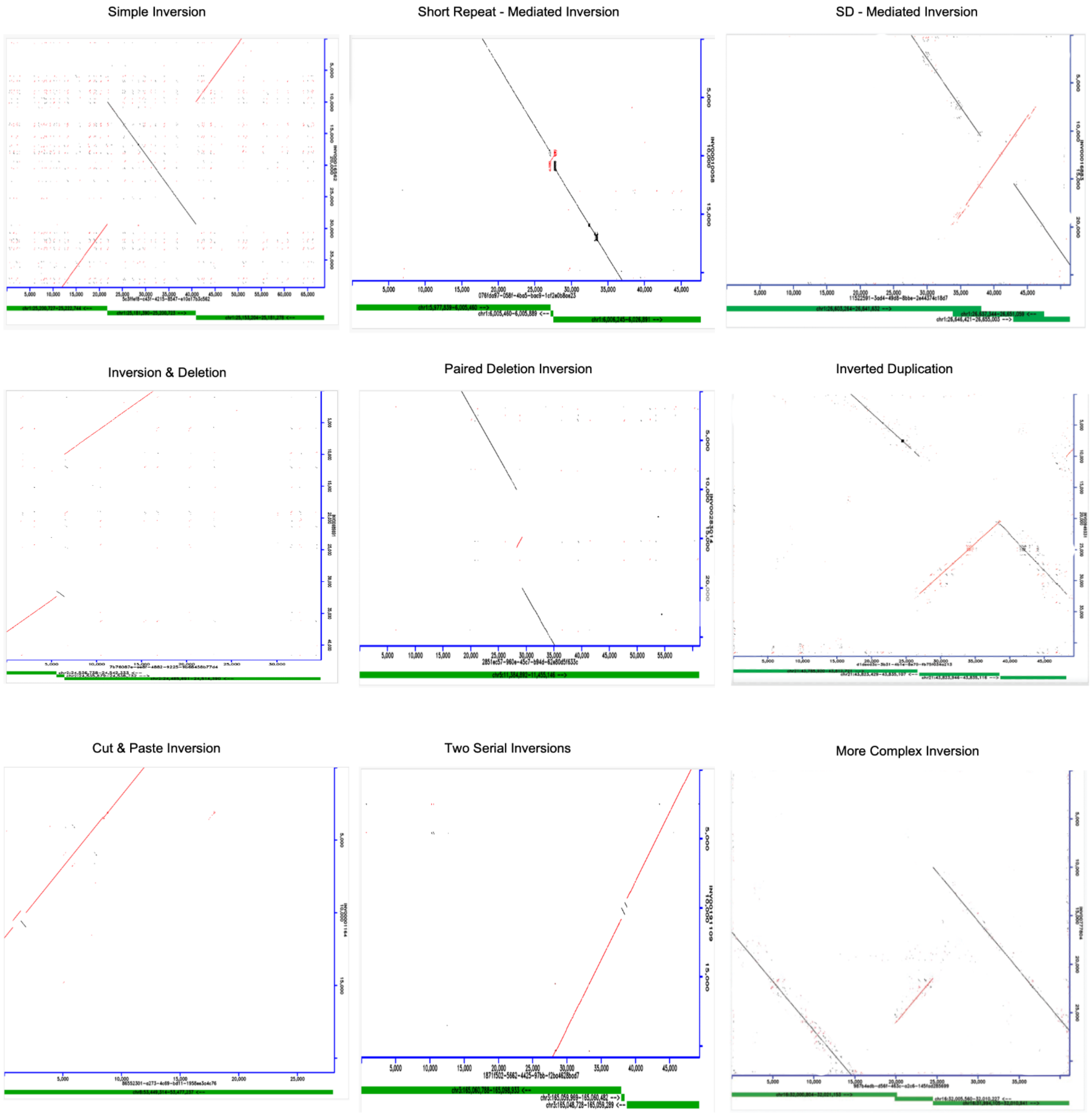


**Supplementary Figure 35:** Representative dot plots illustrating the categorization of inversions extracted from real sample data. The x-axis represents an ONT read in one of the samples in our resource for a specified genomic location, while the y-axis represents the corresponding region in the hg38 reference genome. Each candidate inversion locus was manually examined through dot plot analysis, facilitating the validation of candidate inversion locations and their classification into one of the eight distinct categories showcased in the graph.

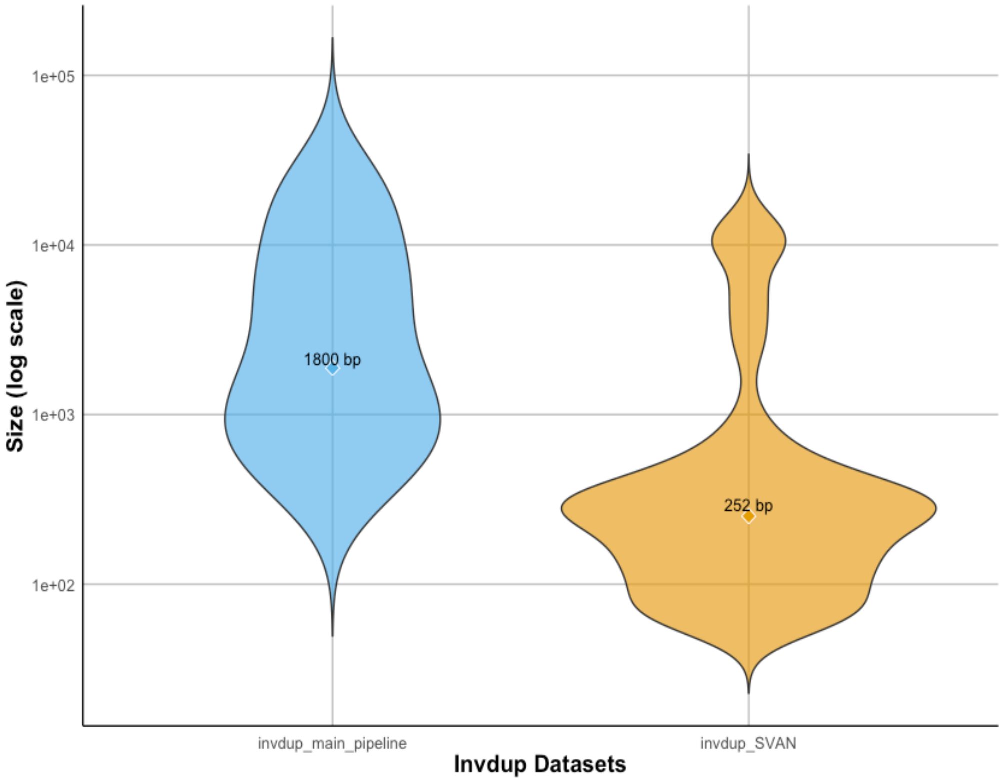


**Supplementary Figure 36:** Violin plots comparing the size distribution of inverted duplications detected through the main inversion detection pipeline versus SVAN. The x axis represents the two callsets, while the y axis represents the size of the inversions identified in each callset. The median size of inverted duplications from the main pipeline (1,8 kbps), represented in blue, significantly exceeds that of those detected by SVAN (252 bps), illustrated in orange, reflecting the enhanced sensitivity of Delly in detecting inversion polymorphisms larger than 500 bps. It should be noted that inverted duplications identified through SVAN exhibit a 13% false positive rate in the current dataset.

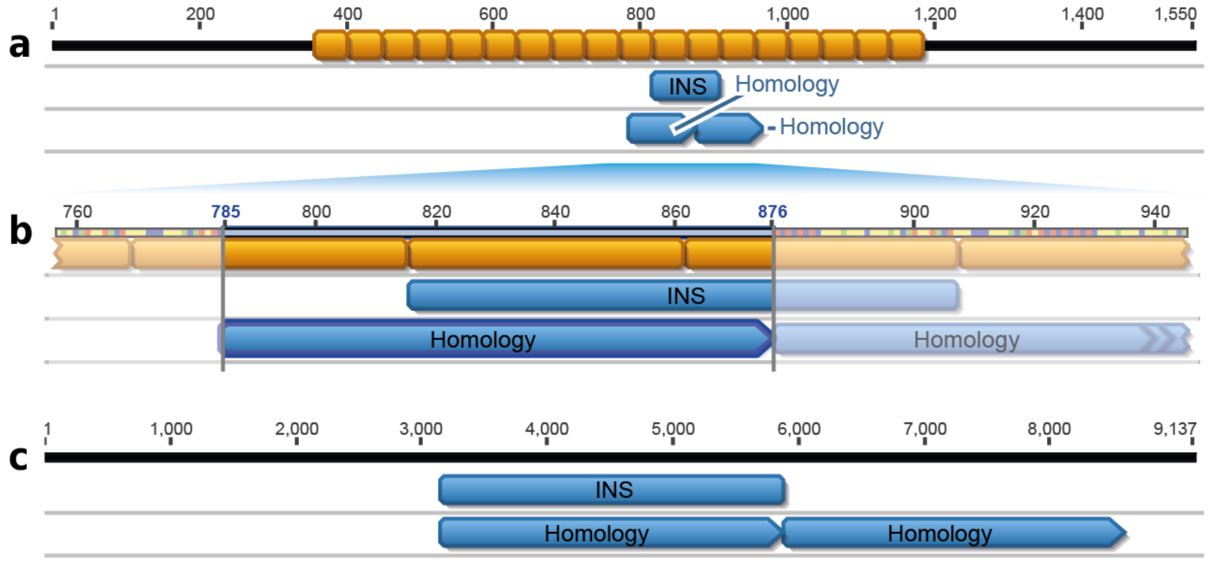


**Supplementary Figure 37: 92 bp VNTR insertion and 2.7 kb tandem duplication with detected homologies.** The top row in each panel shows a region of the T2T reference with the insertion sequence implanted. **a**) A 92 bp insertion (INS, turquoise) is inserted into a repeat (orange boxes represent one repeat unit). **b**) Detailed view of a): The insertion contains two repeat units and the detected homologous flanking sequences (Homology, turquoise) encompass a total of two repeat units as well. **c)** Tandem duplication of a 2.7 kb sequence results in the detection of 2.7 kb long homologous stretches spanning the insertion (left) and the duplicated segment (right).

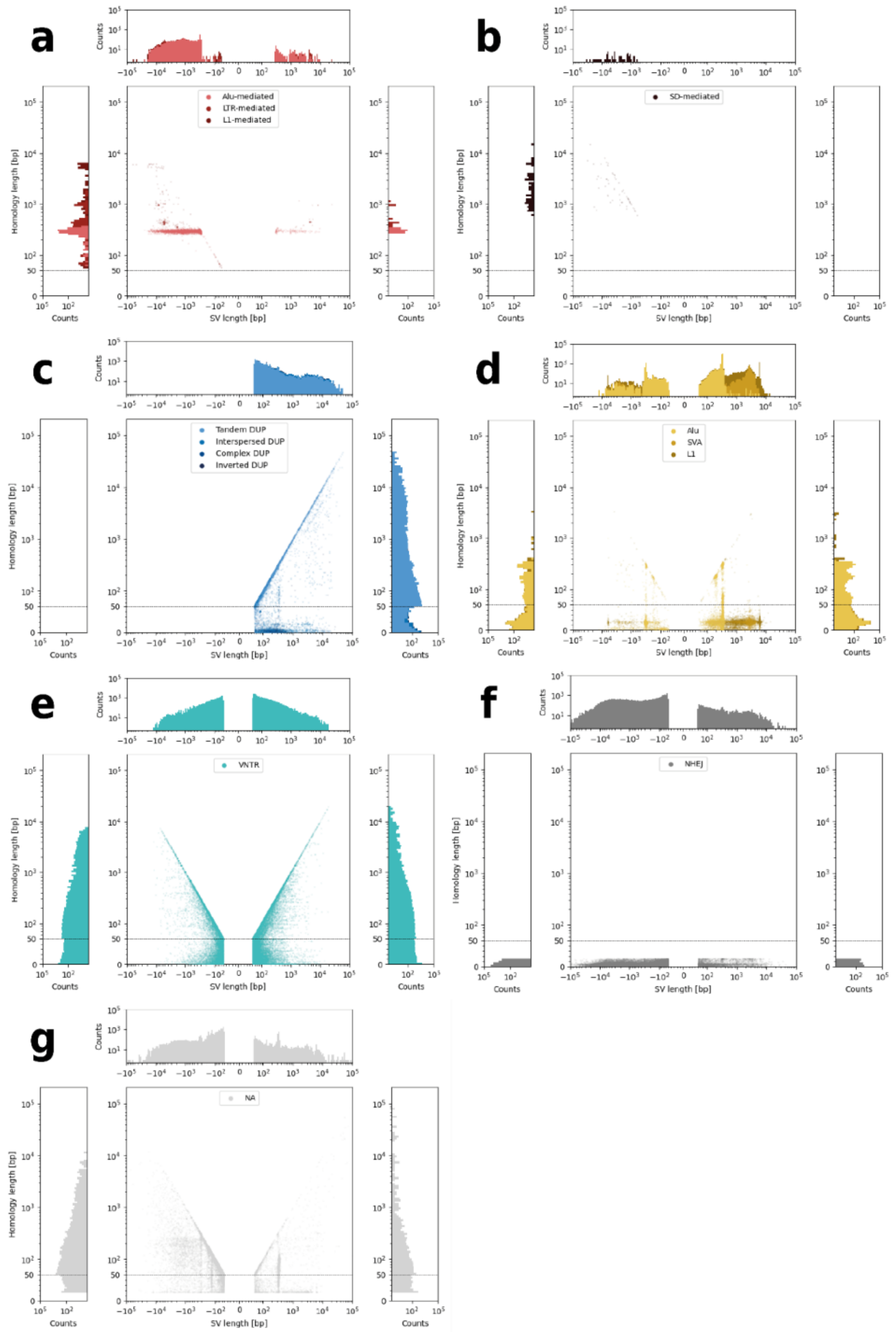


**Supplementary Figure 38: SV breakpoint homology and microhomology landscape in a thousand human genomes sequenced with long reads separated by SV annotation.** For all SV homology and microhomology was determined. SVs were annotated using the SVAN pipeline or leveraging repeat elements inside the detected homologous stretches at the SV breakpoints. Using this annotation procedure SV were grouped into **a)** repeat-mediated SV, **b)** segmental duplication-mediated SV, **c)** duplications, **d)** mobile elements, **e)** VNTRs, **f)** NHEJ-mediated SV and **g)** not-classified SV. For each group the SV length was scattered against the (micro)homology length (center plot). Adjacent histograms show the distribution of SV length (top) and homology for deletions (left) and insertions (right). The axes are linear from 0 to 50 bp and log-scale afterwards, which is denoted by a dashed line. The colors used correspond to Figure 5.

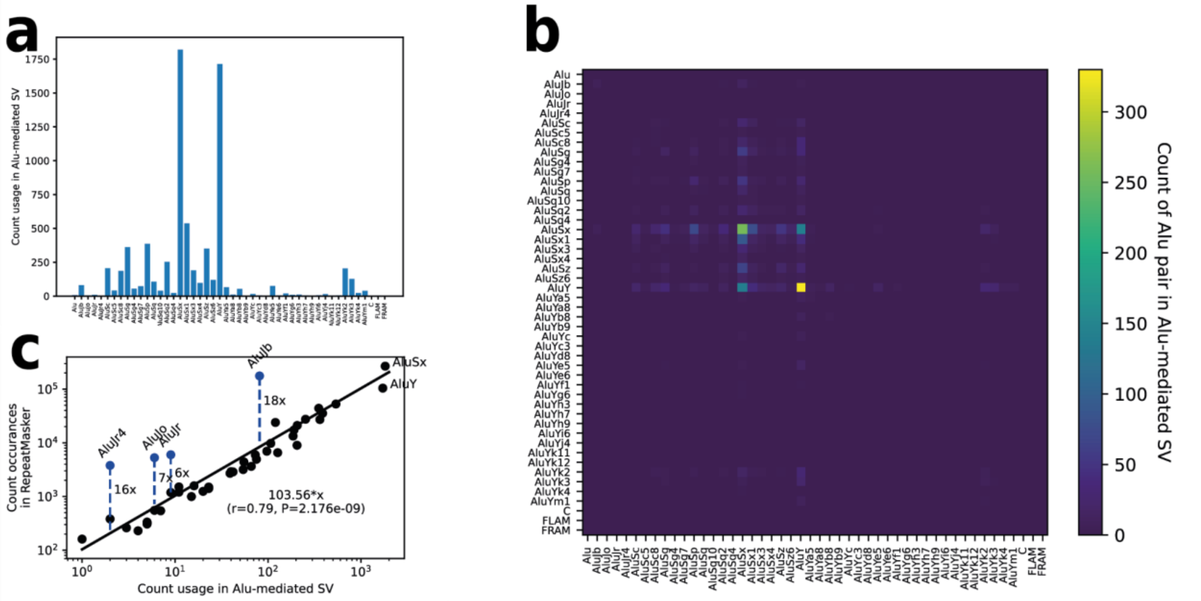


**Supplementary Figure 39: Alu element usage in Alu-mediated SV**. **a)** Counts of individual Alu-elements at breakpoint flanks. **b)** Heatmap showing the abundance of different Alu element pairs at the flanks of Alu-mediated SV. **c)** Scatter plot showing the relationship between the number of Alu elements at the flanks of Alu-mediated SV and inside the T2T RepeatMasker track. Members of the AluJ family are colored blue. Fold-change between AluJ occurrences in the T2T genome and their expected contribution to Alu-mediated SV (as predicted by the linear regression) is shown in labeled, dashed blue lines.

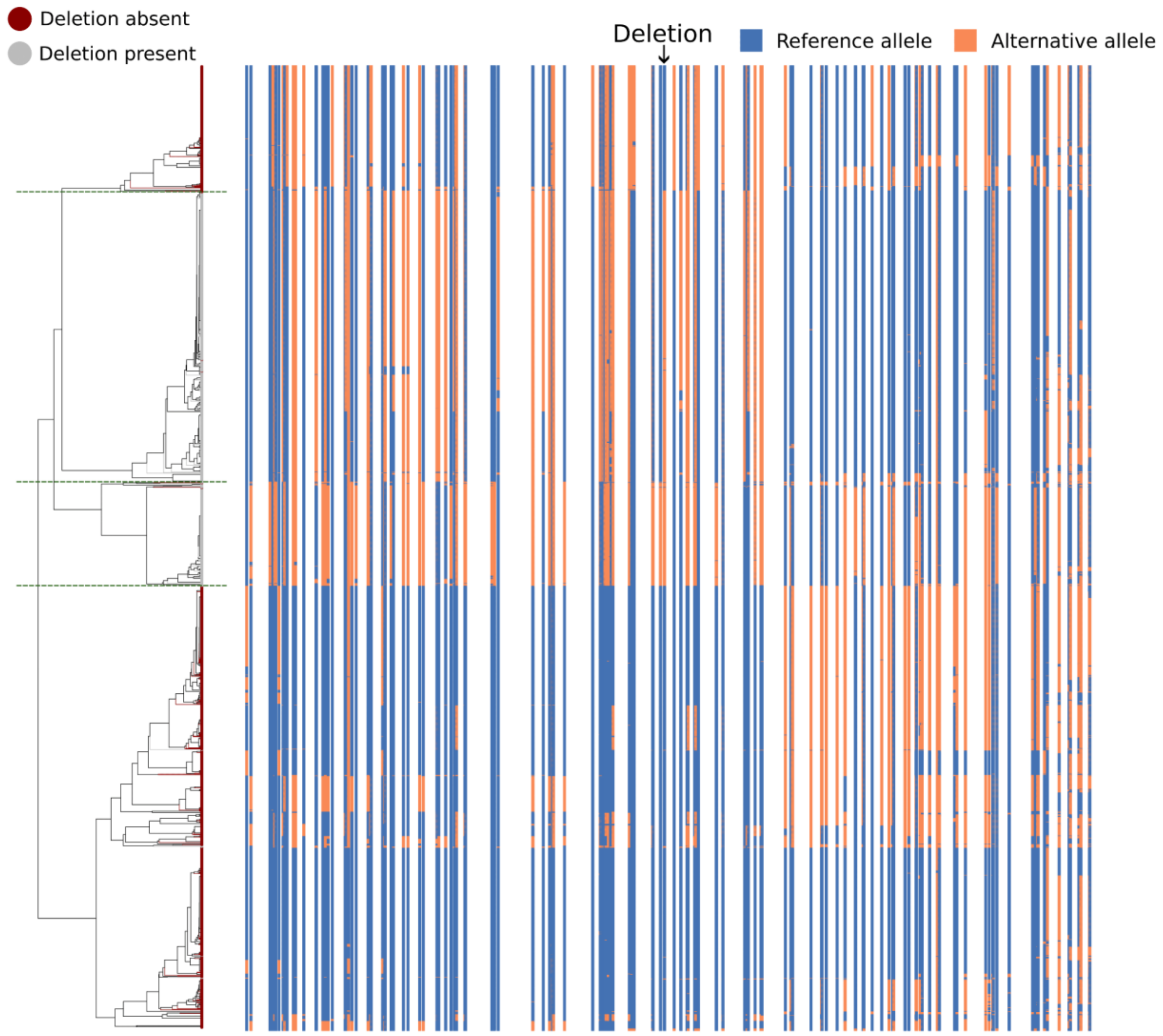


**Supplementary Figure 40:** A potentially recurrent 806 bp deletion at 12p13.3 mediated by an AluSx-AluY pair. Figure shows the variation of haplotypes in a 100kb window centered around the deletion and the relationship between haplotypes with (grey) and without the deletion (red). Dendrograms of haplotypes are plotted using a centroid hierarchical clustering method. Green dashed lines represent the separation of four haplotype groups shown in **Fig. 5g**. In each haplotype, reference and alternative alleles are shown in blue and orange, respectively.

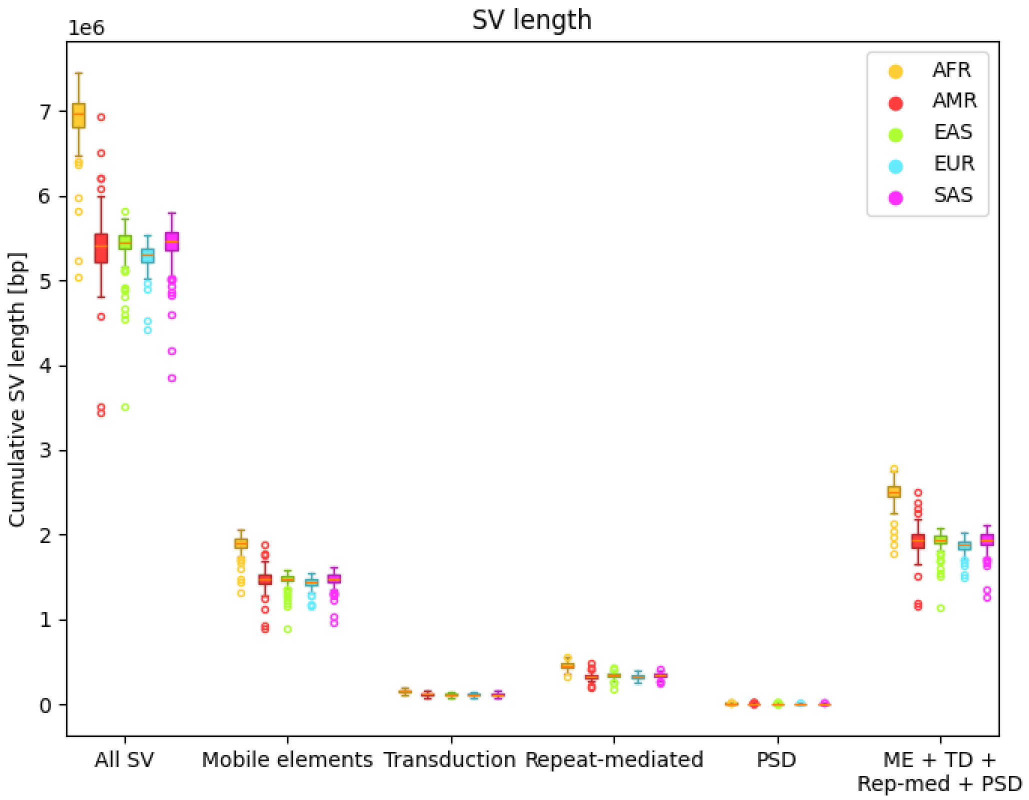


**Supplementary Figure 41: Cumulative SV length per sample for subsets of mobile-element associated SVs stratified by superpopulation.** From the callset, different SV subsets were extracted and the cumulative SV length per sample was determined by summing the length of each SV present in the sample. For the calculation of cumulative transduction size not the SV length but the transduction length determined by SVAN was summed. The results were stratified by superpopulation. ME: mobile elements, TD: transduction, PSD: pseudogene

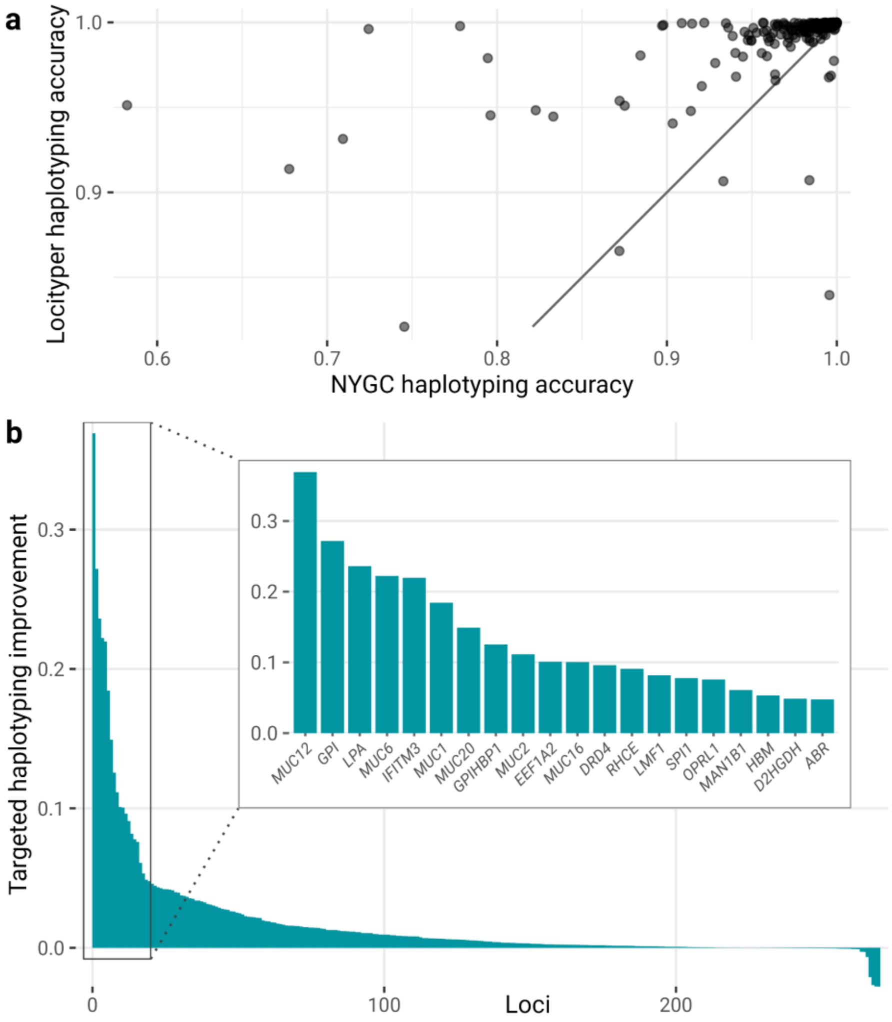


**Supplementary Figure 42: Targeted haplotyping accuracy across 270 medically relevant loci.** Haplotyping accuracy is calculated as sequence similarity between two predicted locus haplotypes and actual locus haplotypes, extracted from the whole genome assemblies for 1 HPRC and 8 HGSVC samples. **a)** Comparison of haplotyping accuracy for short read based NYGC call set and ONT based haplotypes, predicted by Locityper. **b)** Improvement in haplotyping accuracy (Locityper accuracy minus NYGC accuracy) across 270 loci. The inset shows 20 genes with the highest improvement in haplotyping accuracy.

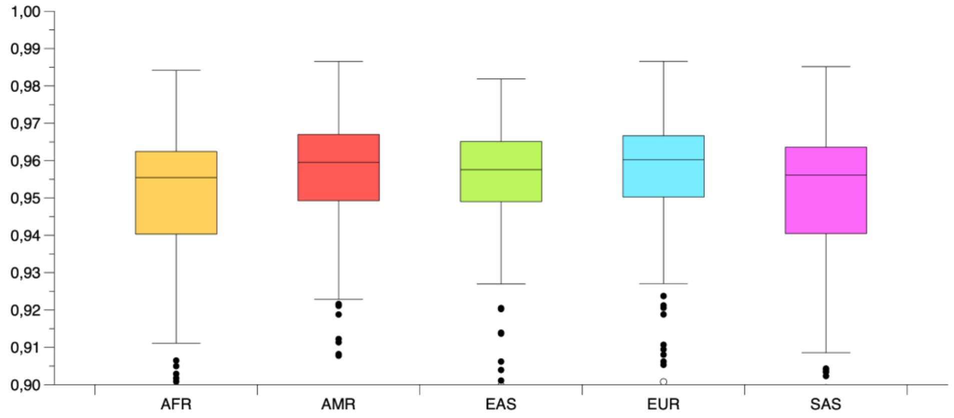


**Supplementary Figure 43:** Fraction of CHM13 bases covered at least five-fold. Y-axis truncated at 0.9.

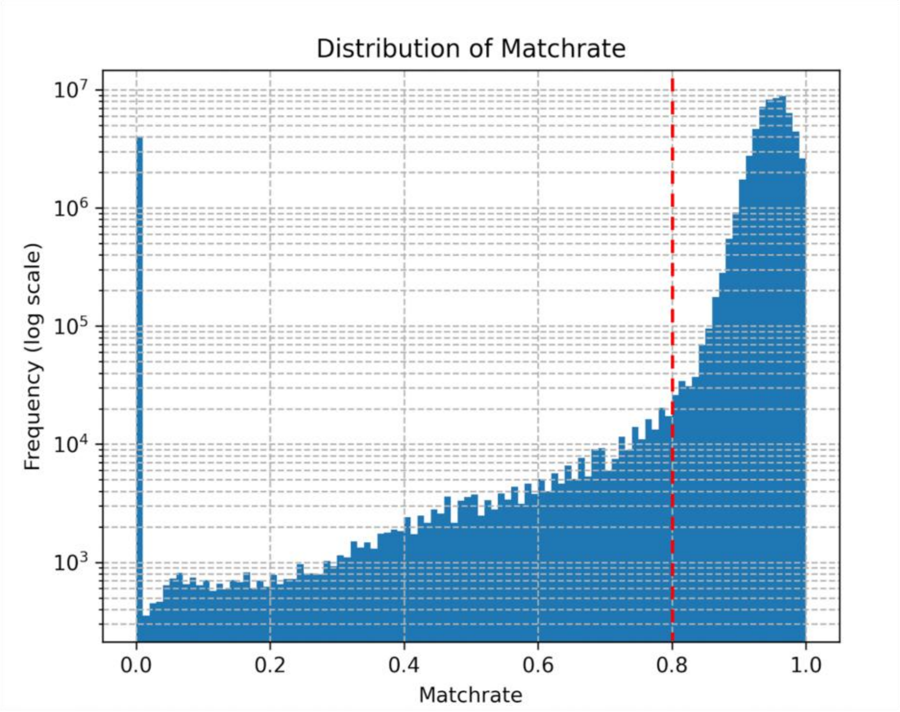


**Supplementary Figure 44: Distribution of Matchrate Indicating Regions Selected for Realignment.** The histogram displays the frequency distribution of match rates across genomic regions in NA12878, plotted on a logarithmic scale. A match rate threshold of 0.8 (delineated by the dashed red line) was established based on the observed distribution, where regions exhibiting a match rate below this value were identified as candidates for realignment. These regions, representing a significantly lower match rate compared to the majority, were subsequently realigned using NGMLR to enhance the accuracy of inversion detection. This thresholding approach ensures the prioritization of genomic regions most likely to benefit from the increased sensitivity and specificity of NGMLR in identifying small inversions.

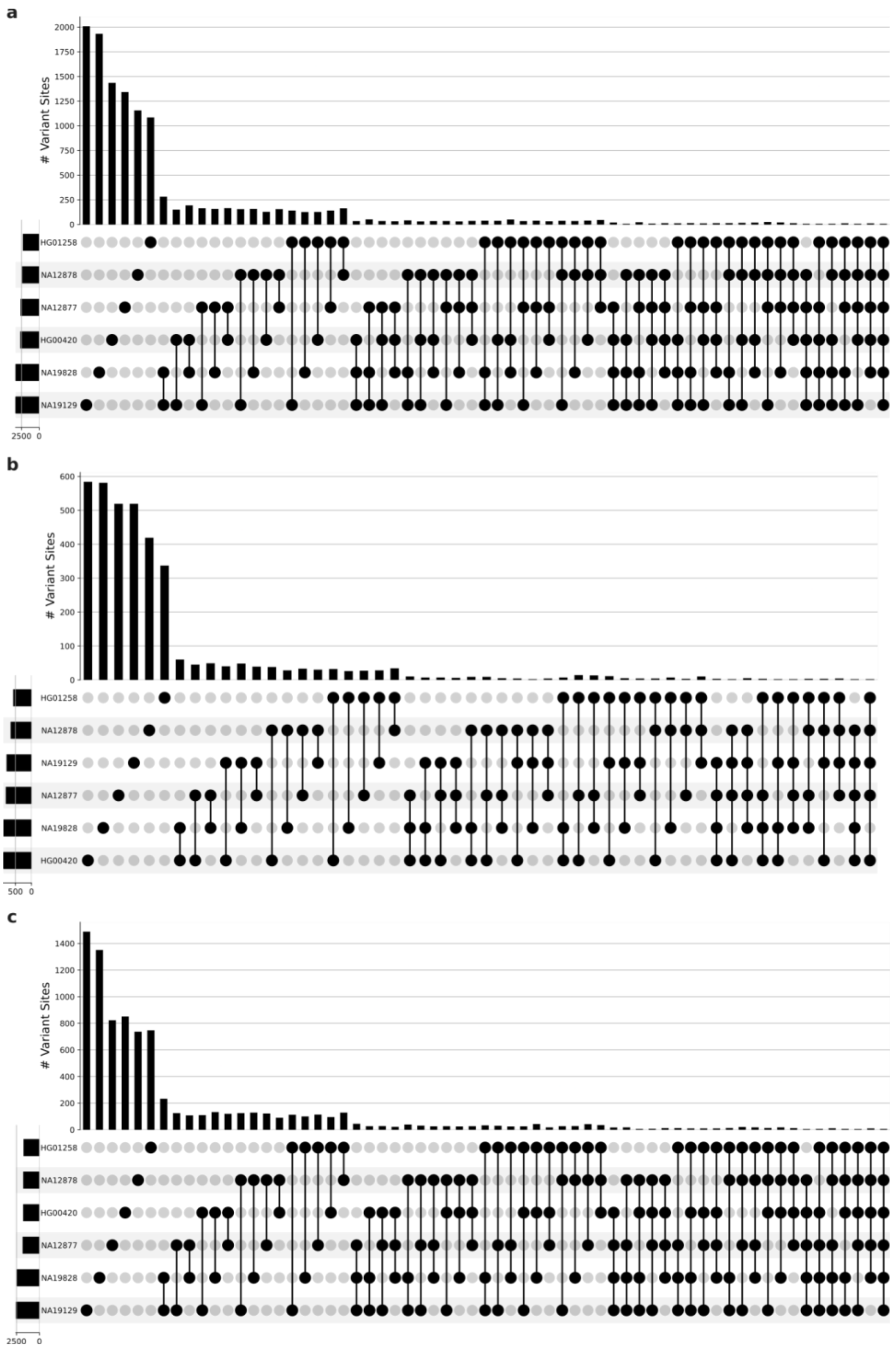


**Supplementary Figure 45:** The figure shows the intersection of the variants which failed Mendelian consistency as an upset plot. The genotypes for which the upset plot has been made come from Giggles genotyping on the HPRC_mg_44+966 graph which have been further filtered to give the strict set. **a)** shows the intersection of all variants, **b)** for variants from biallelic bubbles and **c)** for variants from multiallelic bubbles. The six families are represented by the child sample.

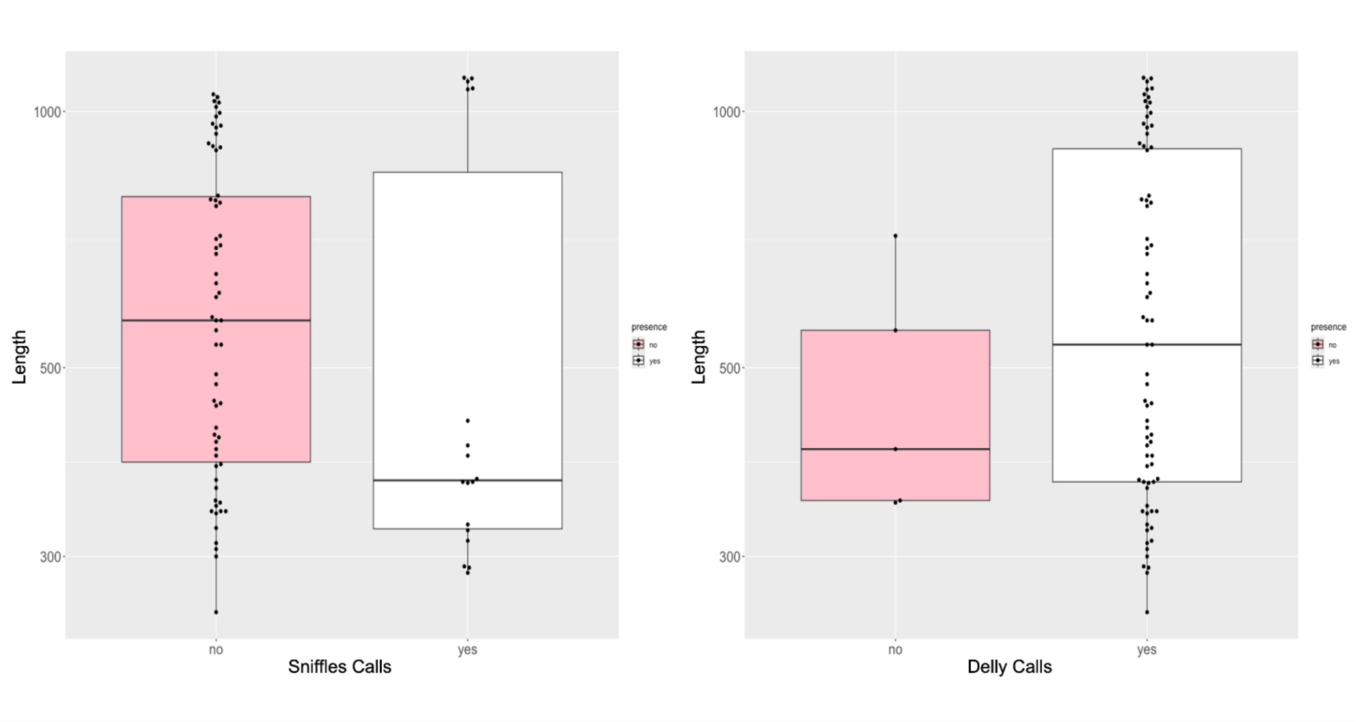


**Supplementary Figure 46: Performance of Delly and Sniffles in the detection of genomic inversions in simulated data.** Boxplot comparison of the inversion detection accuracy of Delly and Sniffles in simulated data for inversions spanning from 250 bps to 1,1 kb in size. The y-axis indicates the inversion length. The color coding represents the presence (white) or absence (pink) of accurate inversion detection, based on simulated benchmarks. As illustrated in the graph, Delly consistently identifies accurately inversions greater than 200 bps, whereas Sniffles only performs accurate inversion detection for inversions that exceed 1.1 kbps.

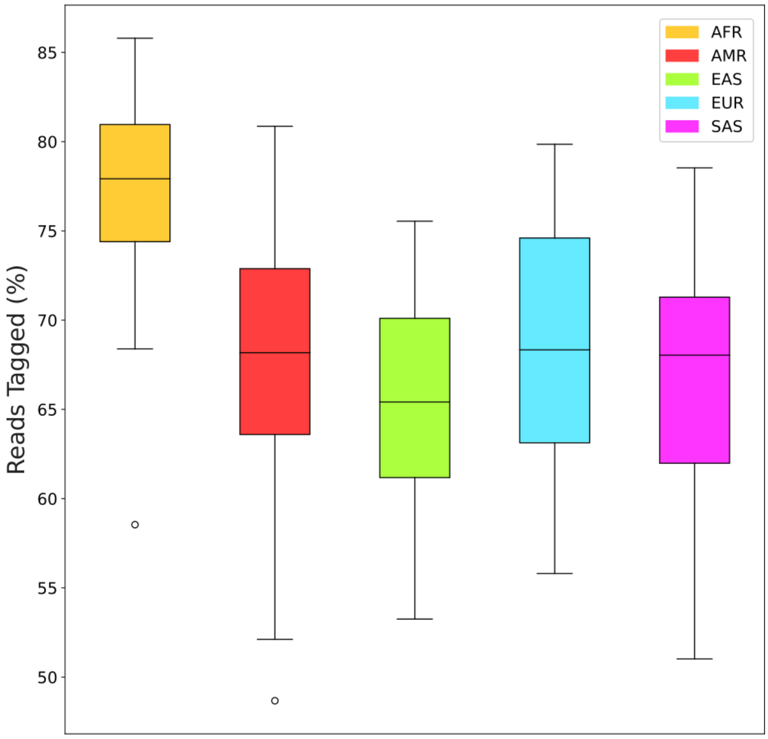


**Supplementary Figure 47:** The percentage of reads tagged by WhatsHap[6] haplotag command. The ONT reads for each sample are haplotype-tagged using the New York Genome Center (NYGC) statistical phased VCF[7].
